# Unconventional biocatalytic strategies orchestrate synthesis of the nucleoside analog sinefungin

**DOI:** 10.64898/2026.05.21.726688

**Authors:** Chi-Fang Lee, Tianhui H. Zhou, Songyi Xue, Lingyang Zhu, Wilfred A. van der Donk, Michael F. Freeman

## Abstract

Sinefungin is a potent nucleoside antimetabolite of *S*-adenosylmethionine (SAM), yet its biosynthesis has remained unclear for decades. Here we detail the identification and characterization of the complete sinefungin biosynthetic gene cluster (BGC) from *Streptomyces incarnatus* NRRL 8089. In vitro and in vivo analyses demonstrate that the defining carbon–carbon (C–C) bond is formed not by the long-hypothesized PLP-dependent process, but by a vitamin B_12_-dependent radical SAM enzyme. Using isotope-labeled cofactors and substrates, we provide evidence that the adenosyl group of sinefungin atypically originates from adenosylcobalamin via a homolytic *S*_*H*_2 substitution, establishing a rare instance where adenosylcobalamin is enzymatically consumed during the reaction. Furthermore, the pathway utilizes a cryptic phosphorylation-dephosphorylation strategy to control intermediate processing and substrate recognition. We also characterize two peptide aminoacyl-tRNA ligases (PEARLs) that append alanines onto the nucleoside scaffold using tRNA-activated amino acids. The PEARLs act directly on small molecules rather than macromolecular substrates, with one PEARL capable of iterative elongation. Finally, we leverage these enzymes in a reduced multi-enzyme cascade to biosynthesize sinefungin. Together, these findings redefine radical-mediated C–C bond formation and pearlin enzyme versatility, unlocking biocatalytic possibilities to produce amino acid-nucleoside conjugates.

**Graphical Abstract:** 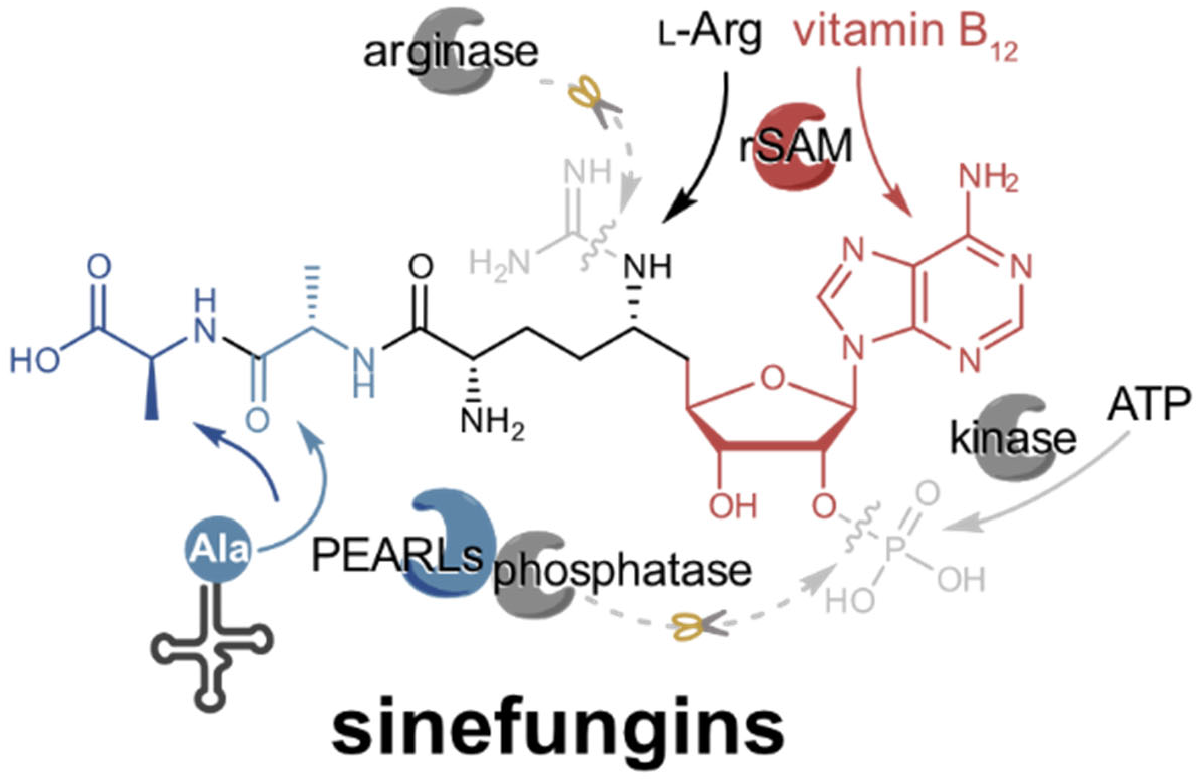

## Introduction

Sinefungin is a nucleoside antimetabolite of the universal methyl donor *S*-adenosylmethionine (SAM).^1^ SAM is an essential cofactor for all forms of life that plays crucial roles in epigenetics, aging, DNA repair, and modulation of protein function.^2,3^ In contrast to the plethora of natural products mimicking essential cofactors such as ATP,^4^ sinefungin is the only known natural SAM pan antimetabolite aside from the SAM breakdown products *S*-adenosylhomocysteine (SAH) and methylthioadenosine.^5^ Sinefungin (**1**; A-9145, 32232 RP) competitively inhibits SAM-binding enzymes through structural mimicry via methylene-amine substitution of the methyl-sulfonium (**Figure 1a**). Consequently, sinefungin initially showed promising bioactivities as an anti-fungal,^6^ anti-viral,^7^ anti-cancer,^8^ and anti-parasitic agent,^9^ including showing nanomolar activity against the causative agent of malaria, *Plasmodium falciparum*.^10^ Nonetheless, development of sinefungin as a therapeutic has been limited due to the nephrotoxic activity of the natural metabolite^11^ and challenges in its synthesis to efficiently install the correct configuration at *C*6’.^12^

**Figure 1.**
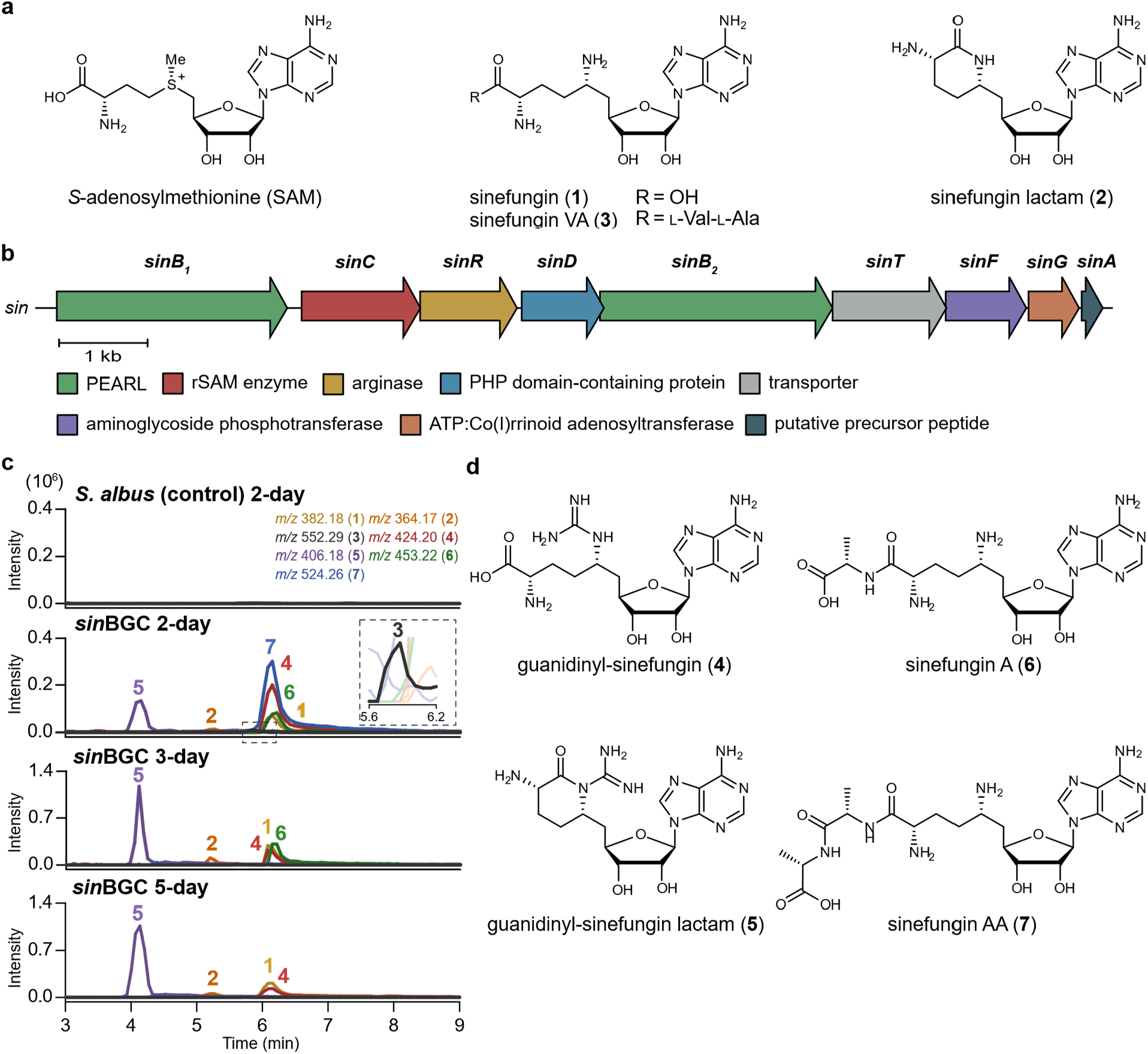
Identification of the sinefungin biosynthetic gene cluster and new derivatives. **a**, Chemical structures of SAM and previously reported sinefungin analogs (**1**–**3**). **b**, Genetic organization of the *sin* BGC from *S. incarnatus* (scale bar, 1 kb). **c**, Selected LC-MS extracted ion chromatograms (positive electrospray ionization; EIC(+)) of metabolite extracts from *S. albus* (negative control) and the *sin* BGC expression *S. albus* after 2, 3 and 5 days of fermentation. Inset provides a magnified view of the trace in the boxed retention region showing compound **3**. Data are representative of three independent biological replicates. **d**, Chemical structures of new sinefungin derivatives (**4**–**7**) identified in this study.

Sinefungin (**1**; A-9145, 32232 RP) and sinefungin lactam (**2**; compound II) were isolated in the early 1970s from *Streptomyces griseolus*^1,13,14^ and soon after from *Streptomyces incarnatus*^15,16^ (**Figure 1a**). Early biosynthetic feeding studies showed high incorporation rates of ^14^C-adenine and adenine-containing nucleosides/tides, and conflicting reports of either ornithine or arginine incorporation into the natural product.^17–20^ Cell-free assays with partially purified proteinaceous fractions revealed additions of the cofactor pyridoxal phosphate (PLP) and Mg^2+^, Mn^2+^, and Co^2+^ metal ions enhanced the production of sinefungin.^16,21^ These data lead to the hypothesis that a PLP-dependent enzyme forms the key carbon– carbon (C–C) bond between adenosine and ornithine/arginine.^16^ Despite further studies and the sequencing of the *S. incarnatus* NRRL 8089 genome in 2015,^22^ details concerning sinefungin biosynthesis have remained a mystery for over 50 years.

Here we report the identification and characterization of the sinefungin biosynthetic gene cluster from *S. incarnatus* NRRL 8089. Through a comprehensive series of in vivo and in vitro analyses, we identify a suite of new and previously isolated sinefungin-related metabolites. Additionally, we provide evidence for the activities of all encoded biosynthetic enzymes within the cluster and propose a biogenic pathway for sinefungin and its congeners. Amongst our findings, we reveal a cryptic phosphorylation and dephosphorylation strategy for sinefungin maturation. We also show activities of two PEARL enzymes that create amino acid derivatives of the sinefungin scaffold using tRNA-conjugated amino acids. Unlike all known PEARLs that act only on peptide or protein substrates, the PEARLs act on small nucleoside metabolites, with one enzyme capable of catalyzing iterative amino acid extensions. Most strikingly, we reveal the key C–C bond formed during the biosynthesis is catalyzed not by a PLP-dependent process, but via a vitamin B_12_-dependent, radical-SAM enzyme. Through a series of coupled in-vitro experiments producing labelled precursors and products, we provide compelling evidence that the sinefungin adenosyl group originates from adenosylcobalamin (AdoCbl) and not SAM. Our results agree with and expand on a very recent disclosure on sinefungin biosynthesis involving the *snf* BGC from *Streptomyces* sp. K05-0178.^23^ Altogether, our findings expand current knowledge of biocatalytic strategies to create nucleoside natural products.

## Results and Discussion

### Identification of the sinefungin biosynthetic gene cluster in *S. incarnatus*

We first revisited the hypothesis that the key C–C bond between ornithine and adenosine in sinefungin was catalyzed by a PLP-dependent enzyme. Analysis of the *S. incarnatus* genome revealed >70 putative PLP-dependent encoded enzymes that gave little insight into which loci may be involved in sinefungin production. Next, we focused on whether ornithine or arginine was more directly incorporated into the natural product. We posited that should arginine be a more direct substrate, an arginase would likely be found in the biosynthetic gene cluster to remove the guanidino group of arginine or an arginine-containing precursor.^24,25^ Only two arginase homologs were identified in the *S. incarnatus* genome, with one located in a nine-gene cassette amongst several genes commonly found in natural product biosynthetic pathways (**Figure 1b, Table S1**). In fact, gene *sinA* bore hallmarks of a ribosomally synthesized and post-translationally modified peptide (RiPP) precursor, including a potential double glycine (VAGG) motif often involved in proteolytic maturation.^26,27^ RiPPs are natural products first translated as peptide precursors and then post-translationally modified on their C-termini prior to proteolytic maturation, yielding the final natural product.^28^ Curiously, the C-terminal sequence of *sinA* encodes the sequence RVA, corresponding strikingly well to the related metabolite sinefungin-Val-Ala (**3**, sinefungin VA) isolated from *Streptomyces* sp. K05-0178.^29^ Additionally, two PEARL enzymes (SinB_1_ & B_2_) are encoded in the BGC. Pearlins, the products of PEARL-containing BGCs, are members of a new RiPP family that use tRNA-activated amino acids to append and reload amino acids often onto RiPP precursor peptides.^28,30,31^ These observations lead us to believe this BGC may be responsible for sinefungin biosynthesis, despite the lack of proteins predicted to use the cofactor PLP.

Indeed, heterologous expression of the putative gene cluster (*sin* BGC) in *Streptomyces albus* resulted in sinefungin production, as well as multiple derivatives, over several days of fermentation (**Figure 1c**). Purification in conjunction with mass spectrometric and NMR analyses identified guanidinyl-sinefungin (**4**), guanidinyl-sinefungin lactam (**5**), sinefungin lactam (**2**), sinefungin-Ala (**6**, sinefungin A), sinefungin-Ala-Ala (**7**, sinefungin AA) along with sinefungin (**1**) and trace amounts of sinefungin VA (**3**) (**Figure 1d; see Supporting Information for full characterization data**). We note that sinefungin VA has not been previously detected in *S. incarnatus*, and that sinefungin A (**6**) and sinefungin AA (**7**) are new amino acid-nucleoside conjugates not previously isolated from *S. incarnatus* or any other source. Repeated experiments consistently revealed compounds **3, 6**, and **7** were produced early during fermentations, and later lost, with concomitant increases in compounds **1, 2, 4**, and **5**. These data demonstrate the *sin* BGC is responsible for sinefungin biosynthesis, and that the larger sinefungin amino acid derivatives may be further processed or serve as intermediates in the pathway.

### Removal of individual genes from the *sin* BGC leads to accumulation of pathway intermediates

To determine whether sinefungin is a RiPP natural product, we deleted *sinA* in our *S. albus sin* BGC fermentations. Surprisingly, the metabolic profile remained unaffected upon removal of *sinA* from the *sin* BGC expression strain (**Figure 2**). This result suggests sinefungin is not a RiPP metabolite and is biosynthesized via an alternate pathway. In response, we performed a series of top-down (**Figure 2**) and bottom-up (**Figure S1**) heterologous expressions along with in-vitro enzymatic assays using purified recombinant proteins (**Figure S2**) to unravel the details of the pathway. Only a single gene (*sinC*), when deleted from the *sin* BGC operon, resulted in the loss of all identified sinefungin-related metabolites during *S. albus* coexpressions as monitored by liquid chromatography coupled with mass spectrometry (LC-MS). Conversely, guanidinyl-sinefungin (**4**) was universally produced in all top-down *sin* BGC expressions except for the *sinC*-deleted expression strain. These data suggest *sinC* acts first in the pathway and may be responsible for the key C–C bond formation between arginine and the adenosyl group. In addition, *sinR* (arginase) and *sinF* (kinase) deletions only produced guanidinyl-sinefungin (**4**) during heterologous expressions and thus were also likely to act early in the pathway. Surprisingly, the arginase SinR displayed no in-vitro activity with either L-arginine or guanidinyl-sinefungin (**4**) (**Figure a**). Guanidinyl-sinefungin (**4**) was, however, a substrate for the kinase SinF, and SinF efficiently phosphorylated the 2’-hydroxyl of **4** to yield 2’-phosphoguanidinyl-sinefungin (**8**) during in-vitro assays supplemented with Mg^2+^ and ATP (**Figure b**). Although not observed in the *sin* BGC heterologous expression experiments, compound 8 represents an on-pathway intermediate because SinR assays with **8** yielded the new compound 2’-phosphosinefungin (**9**), as determined by NMR spectroscopy (**see Supporting Information**). These observations revealed a 2’-phosphate requirement for SinR to hydrolyze the guanidinium of its substrate (**Figure 3a, 3c**).

**Figure 2.**
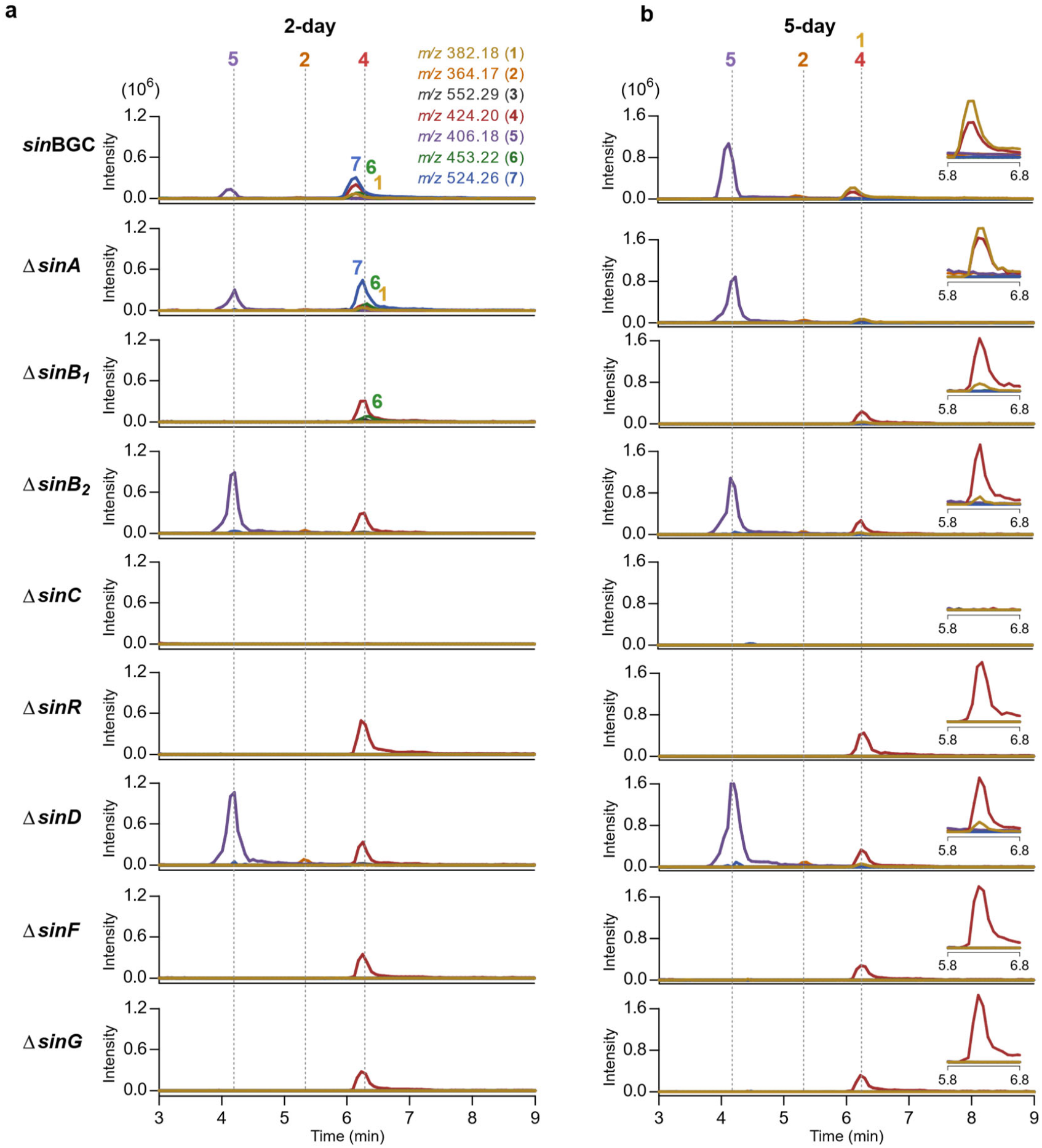
Analysis of metabolites from top-down expression of the *sin* BGC in *S. albus. a, b*, Selected LC-MS extracted ion chromatograms (EIC(+)) of extracts from *S. albus* harboring the indicated gene deletion variants of the *sin* BGC after 2 days (**a**) and 5 days (**b**) of fermentation. Insets provide magnified views of peaks between 5.8 and 6.8 min. Data are representative of three independent biological replicates.

**Figure 3.**
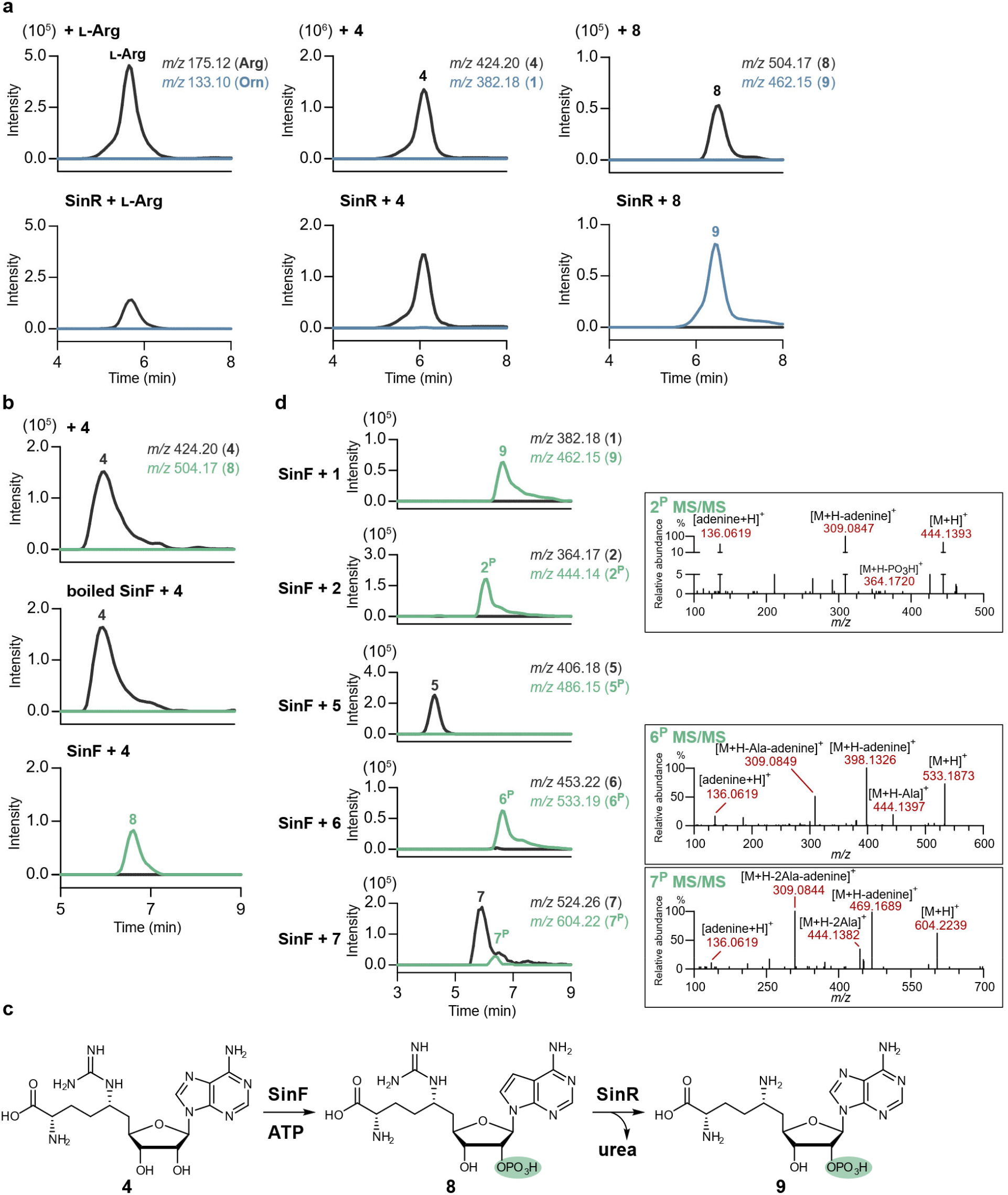
Discovery of a cryptic phosphorylation step by SinF required for SinR substrate recognition. **a**, LC-MS analysis of SinR activity toward different substrates. SinR shows no activity toward L-Arg or **4** but specifically converts the phosphorylated intermediate **8** to **9**. Substrate and product EIC(+) traces are shown in black and blue, respectively. **b**, LC-MS analysis of SinF activity with **4**. SinF phosphorylates **4** to generate **8** in the presence of ATP. The activity of SinF is abolished when the enzyme is heat-denatured (boiled). Substrate and product EIC(+) traces are shown in black and green, respectively. **c**, The proposed biosynthetic order of SinF and SinR. SinF catalyzes 2’-phosphorylation of **4** to generate **8**, which SinR then recognizes and converts to **9. d**, Substrate specificity of SinF. LC-MS chromatograms (left) show SinF activity toward analogs **1, 2, 6**, and **7** with significantly lower conversion rates, and no detectable activity toward substrate **5**. Corresponding LC-MS/MS fragmentation spectra (right) confirm the identity of the phosphorylated products.

Cryptic phosphorylation in nucleoside biosynthetic pathways has been previously reported, and our data is also consistent with the absence of detectable phosphorylated intermediates in these previously studied systems.^32^ Phosphorylations in nucleoside pathways have been postulated to provide a molecular handle for downstream enzymes and/or to prevent premature leakage of intermediates from inside the cell.^32^ We propose an additional rationale for cryptic phosphorylation in sinefungin biosynthesis: to prevent premature sinefungin formation within the cell. Essentially all metabolites detected and purified from our *sin* BGC expressions were isolated from culture supernatant and not the cell mass. Phosphorylation of guanidinyl-sinefungin prevents in vivo production of sinefungin upon hydrolysis of the guanidinium by SinR prior to SinB_1_ activity (**see below**).

Next, we focused on the origins and biosynthetic timings of the new sinefungin amino acid derivatives sinefungin A (**6**) and sinefungin AA (**7**). Careful analysis of the *sinB*_*1*_ and *sinB*_*2*_ gene-deletion expressions revealed a loss of sinefungin A and sinefungin AA production in the Δ*sinB*_*2*_ strain, with a loss of only sinefungin A in the Δ*sinB*_*1*_ strain (**Figure 2**). Complementing the top-down deletion experiments, bottom-up *S. albus* expressions of *sinC, sinR, sinF, sinB*_*2*_, and the putative transporter *sinE* produced sinefungin A (**6**), but not sinefungin AA (**7**), early during fermentations (**Figure S1**).

### In vitro reconstitution of the activity of SinD, SinB_1_ and SinB_2_

Thus, PEARLs SinB_2_ and SinB_1_ were suspected of performing the first and second amino acid extensions respectively, yet the manner for how this may be accomplished was unclear. All pearlins to date have required a protein or large peptide substrate for activity, and all have shown strict specificity for the composition of the protein precursor/peptide C-termini and for the amino acids that are appended.^31,33^ Early in-vitro experiments testing SinB_1_/B_2_ amino-acid extensions of SinA variants failed to produce any products, consistent with SinA being unnecessary for heterologous sinefungin biosynthesis. Intriguingly, a bioinformatic search revealed three homologous *sin* BGC clusters that encode natural fusions of putative *sinD* and *sinB*_*2*_ homologs (accession numbers: WP_256791495, WP_190021322, WP_382036511). These observations led us to assess the activity of SinD and any concerted activities with SinB_2_. In-vitro reactions of SinD alone verified phosphatase activity with 2’-phosphoryl sinefungin derivatives **8** and **9** (Error! Reference source not found.). However, similarly to the kinase SinF (**Figure 2c**), the phosphatase SinD shows promiscuity for multiple substrates (Error! Reference source not found.), and therefore the precise biosynthetic timings for either enzyme cannot be pinpointed through in-vitro substrate profilings alone. We then extended our bottom-up expressions to include SinD. Two-day *S. albus* expressions of SinC/R/F/B_2_/E/D resulted in elevated production of sinefungin A (**6**) compared to expressions without SinD. Unexpectedly, we also observed sinefungin AA (**7**) produced in the experiments (**Figure S1**). These data insinuated SinB_2_ can iteratively ligate two alanines onto this nucleoside scaffold, which would be a first for the PEARL enzyme family.

In response, we performed a series of in-vitro coupled-enzyme assays combining SinF, SinD, SinB_1_, and/or SinB_2_ with different substrates to verify the activities and unravel the biosynthetic timing of the PEARLs. Four in-vitro transcribed isoforms of *S. incarnatus* tRNA^Ala^s were used in combination with the *S. incarnatus* alanine tRNA synthetase to load the tRNAs with L-Ala-2,3-^13^C_2_ for the in-vitro assays. Sinefungin (**1**) was converted to sinefungin A (**6**) in overnight reactions with SinB_2_, SinF, and SinD with all other necessary components and cofactors (**Figure 4**; **Figure S4**). SinF inclusion was necessary for conversion of **1** to **6**; addition of SinD enhanced production of sinefungin A (**6**), comparatively. Of note, sinefungin A (**6**) was not produced when SinB_2_ was replaced with SinB_1_ in these reactions, nor if sinefungin (**1**) was the substrate in the absence of kinase SinF (**Figure 4**). These data reveal SinB_2_ likely accepts 2’-phosphosinefungin (**9**) as a native substrate, and not sinefungin (**1**), to conjugate alanine and produce sinefungin A (**6**) in concert with SinD. In line with our bottom-up in-vivo results, small quantities of sinefungin AA (**7**) were also observed in SinB_2_- and SinF-coupled in-vitro reactions, confirming the ability of SinB_2_ to iterative incorporate two alanines. Nonetheless, addition of SinB_1_ to the reaction significantly enhanced the production of sinefungin AA (**7**) (**Figure 4**).

**Figure 1.**
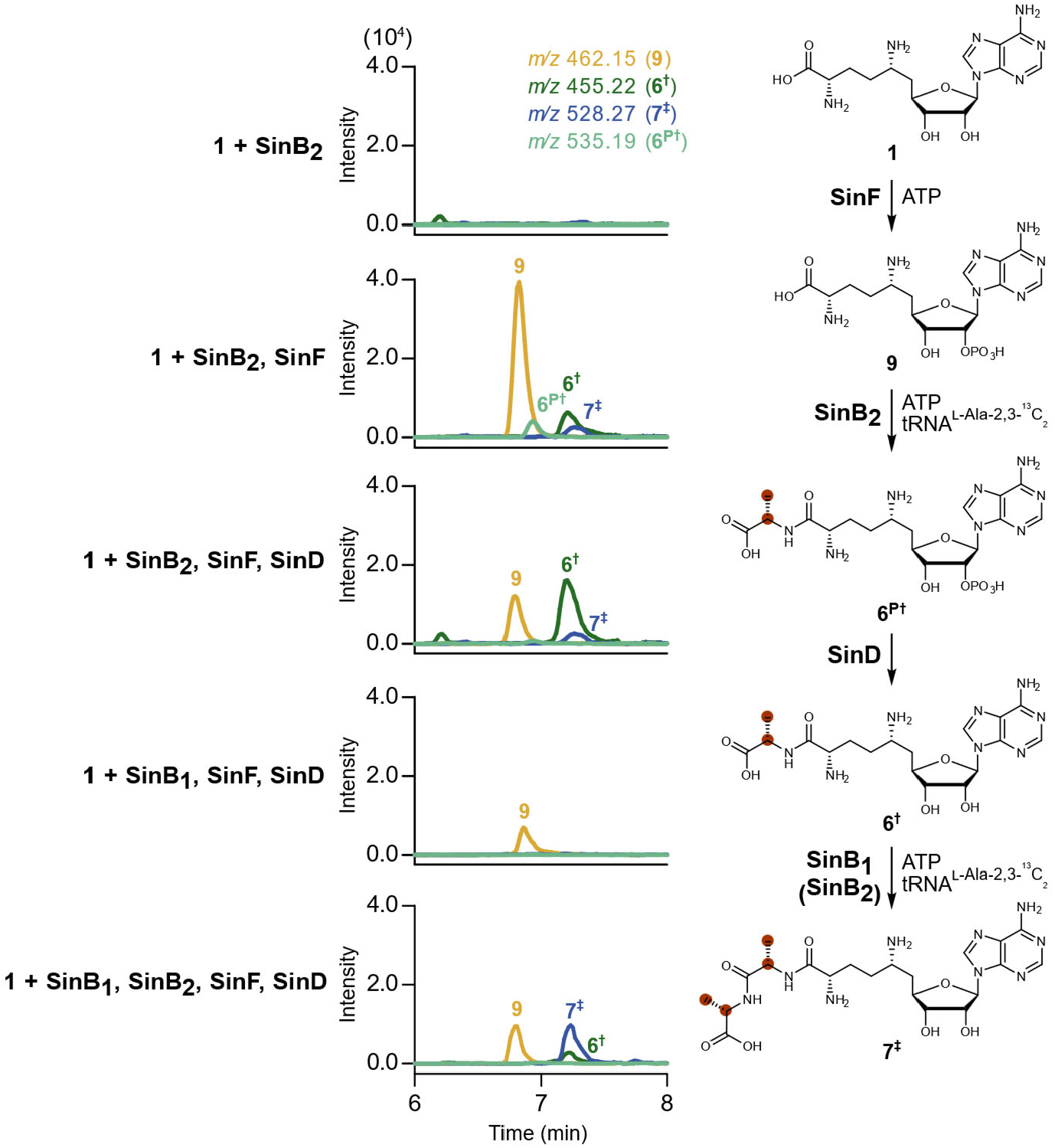
In vitro characterization of the amino acid extensions catalyzed by SinB_2_ and SinB_1_. Enzymatic reactions using **1** as the starting scaffold and tRNA^Ala^s pre-charged with L-Ala-2,3-^13^C_2_ (isotope positions indicated by red circles). Incubation of **1** with SinB_2_ alone shows no conversion, while co-incubation with SinB_2_ and SinF results in the formation of **9** as well as small amounts of isotope-labeled phosphorylated sinefungin A (**6**^**P†**^), isotope-labeled sinefungin A (**6**^**†**^), and dually labeled sinefungin AA (**7**^**‡**^). Addition of SinD in SinB_2_ assays shifts conversion of the phosphorylated intermediate **6**^**P†**^ to **6**^**†**^, while inclusion of SinB_1_ efficiently shifts the cascade towards **7**^**‡**^. SinB_1_ is unable to accept **1** as a substrate in assays including SinF and SinD. Isotopically labeled atoms are colored red.

To evaluate SinB_1_ activity separately from SinB_2_, sinefungin A (**6**) was chemically synthesized and tested in vitro using four *S. incarnatus* tRNA^Ala^s and tRNA alanine synthetase to load L-Ala-2,3-^13^C_2_; sinefungin AA (**7**) was successfully produced under these conditions. Unlike SinB_2_, addition of SinF and/or SinD did not boost product formation with SinB_1_, suggesting neither the presence of SinD nor phosphorylation of sinefungin A (**6**) is required for conversion to sinefungin AA (**7**) (**Figure S5**). However, from these data, the precise biosynthetic timing of SinD activity with respect to SinB_1_ cannot be fully resolved. Mechanistic and structural details concerning interactions between SinB_2_ and SinD are the subject of future investigations. Collectively, the iterative activity observed with SinB_2_ and the substrate profiles for SinB_1_ and SinB_2_ differentiate the *sin* BGC PEARLs from all other characterized pearlin examples by not requiring protein or large peptide substrates. In support of these findings, recent bioinformatics and structural analyses of PEARLs suggest their evolution originates from ATP-GRASP enzymes that also do not require protein or peptide substrates for activity.^34^ The existence of the RiPP-like precursor SinA in the *sin* BGC remains unanswered; future work may uncover whether it is an evolutionary remnant or has an alternative function within the cluster.

### Proposed pathway for sinefungin biosynthesis

Our cumulative data allows us to propose a biogenic pathway for sinefungin biosynthesis (**Figure**). SinC catalyzes the first key C–C bond-forming step in the pathway through SAM-derived radical-initiated formation of guanidinyl-sinefungin (**4**) from L-arginine and adenosylcobalamin (regenerated by the Co(I)rrinoid adenosyl transferase SinG). SinF then phosphorylates **4** to produce 2’-phosphoguanidinyl-sinefungin (**8**) prior to hydrolysis of the guanidinium by SinR to afford 2’-phosphosinefungin (**9**). SinB_2_, in concert with SinD, catalyzes tRNA^Ala^-mediated formation of 2’-phosphosinefungin A prior to the phosphatase activity of SinD to produce sinefungin A (**6**). SinB_1_ then ligates an additional L-alanine onto **6** to create sinefungin AA (**7**), which we surmise is the most biosynthetically advanced metabolite produced in vivo prior to export by the major facilitator superfamily transporter SinE. Maturation of sinefungin AA to sinefungin likely arises from extracellular proteolytic activities. Amino acid-nucleoside conjugates, including the nikkomycins,^35^ microcins,^36^ and polyoxins,^37^ act as trojan horse antibiotics to increase cellular uptake via various oligopeptide permeases followed by intracellular release of the warhead by peptidases; **6** and **7** may have similar effects which is a subject of future study. Lactams **2** and **5** appear to be shunt products formed by a yet unidentified enzyme(s). We note the lactam metabolites were never observed in our in-vitro reactions, and it is unclear if the *sin* BGC biosynthetic enzymes participate in their formation or whether one or more host-encoded enzymes are responsible.

The successful identification of the sinefungin biosynthetic machinery provides a streamlined alternative to the multi-step chemical synthesis. We demonstrate that four enzymes are sufficient to convert the core intermediate **4** into sinefungin (**1**). By leveraging *S. albus* as a cellular factory to produce the core intermediate **4**, we bypass the most challenging synthetic hurdle. The in vivo-derived material was then converted to sinefungin (**1**) through a single-pot, multi-enzyme cascade featuring SinF, SinR, and SinD (**Error! Reference source not found**.). The near-complete conversion of **4** into **1** highlights the robustness of these biocatalysts and establishes a modular framework to produce sinefungin without the need for complex chemical protecting group strategies.

### An unusual utilization of adenosylcobalamin

To further investigate the enzymatic machinery responsible for the assembly of the sinefungin core, we focused on the radical-mediated mechanism of SinC. SinC is predicted as a cobalamin-dependent, radical SAM (rSAM) enzyme; enzymes of this class are known to catalyze a diverse array of radical-mediated transformations, including C–C bonds at unactivated carbon centers.^38^ Notably, *sinG*, also present in the gene cluster, is annotated as an ATP:corrinoid adenosyltransferase (ACAT) and is likely responsible for generating adenosylcobalamin (a form of vitamin B_12_) to provide the SinC cofactor.^39,40^ *S. albus* naturally produces vitamin B_12_ ^41^ and also encodes a homolog (WP_003950891) with 66% sequence identity to SinG. We verified SinG can generate adenosylcobalamin via *sinG* genetic complementation of an adenosylcobalamin auxotroph of *Salmonella enterica* that is a triple ACAT knockout (**Error! Reference source not found**.).^42^ Based on these observations, we posited the key C–C bond formation in sinefungin biosynthesis is catalyzed by SinC with support from SinG, rather than the long-held hypothesis that the C–C bond is catalyzed by a PLP-dependent enzyme.

Substantial efforts to produce and purify soluble SinC in concert with solubility tags or coexpressions with loci known to aid in iron-sulfur (Fe-S) cluster assembly and cobalamin uptake (*isc, suf, btu* operons) failed.^43–45^ In response, we engineered a soluble SinC variant (SinC*) with a total of 51 amino acid substitutions. The design was based on a consensus sequence of putative SinC variants found in related BGCs, with a focus on protein surface residue substitutions for charged and polar amino acids (**Error! Reference source not found. Table S5**). The activity of SinC* was first validated in vivo with replacement of SinC in full gene cluster coexpressions in *S. albus* (**Figure S9a**). Subsequently, soluble SinC* was produced and purified from *E. coli* coexpressions with the *isc* (FeS assembly)^43^ and *btu* (B_12_ uptake)^45^ operons under microaerobic induction conditions followed by anaerobic protein purification (**Error! Reference source not found**.). Fe-S cluster reconstitution was performed prior to overnight SinC* in-vitro assays supplemented with SAM, adenosylcobalamin, L-arginine, and the reductants NADPH and methyl viologen. Production of guanidinyl-sinefungin (**4**) in these assays confirmed SinC forms the key C–C bond in sinefungin biosynthesis. (**Figa**) SinC* does not accept D-arginine or L-ornithine as substrates under these conditions (**Figure S9b**). Of note, neither full-length SinA, nor SinA truncations ending in RVA, RV, or R, were substrates for SinC* under these conditions (data not shown), consistent with the SinA deletion results described above..

Radical SAM-mediated adenosylation reactions are rare in the literature, and all known examples adenosylate sp^2^ carbon centers in non-natural substrates.^46,47^ While examples of 4,5-dehydroarginine have been observed in other natural product pathways,^48,49^ we did not suspect the SinC reaction to proceed through a double-bond intermediate. Indeed, incubation of SinC* with ^2^H-L-arginine resulted in a net 1-Da loss during production of **4**, providing strong evidence that SinC is abstracting a single hydrogen during the adenosylation reaction and not proceeding through a double-bond intermediate (**Figure 6a**).

**Figure 5.**
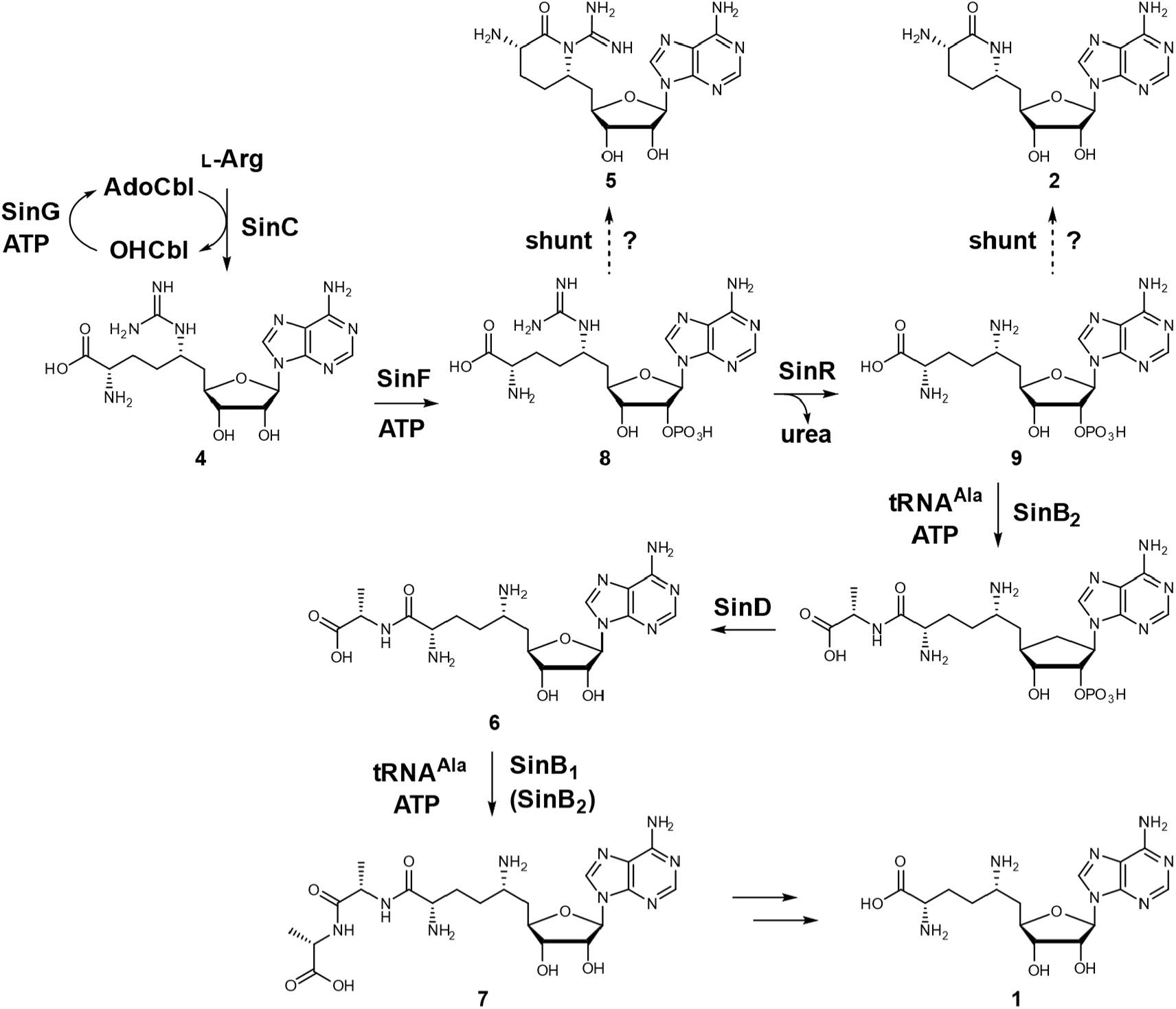
Proposed sinefungin biosynthetic pathway. The pathway is initiated by SinC via the radical SAM-dependent coupling of L-Arg and adenosine from AdoCbl generated by SinG. Intermediate **4** is 2’-phosphorylated by SinF to generate **8**, followed by urea removal by SinR to form **9**. Subsequently, the first alanine is appended by SinB_2_ and the second by either SinB_1_ or SinB_2_ with the assistance of SinD. The resulting tripeptide derivative is proposed to be cleaved by an unknown peptidase to yield sinefungin (**1**). Dashed arrows indicate putative shunt products **2** and **5**.

**Figure 6.**
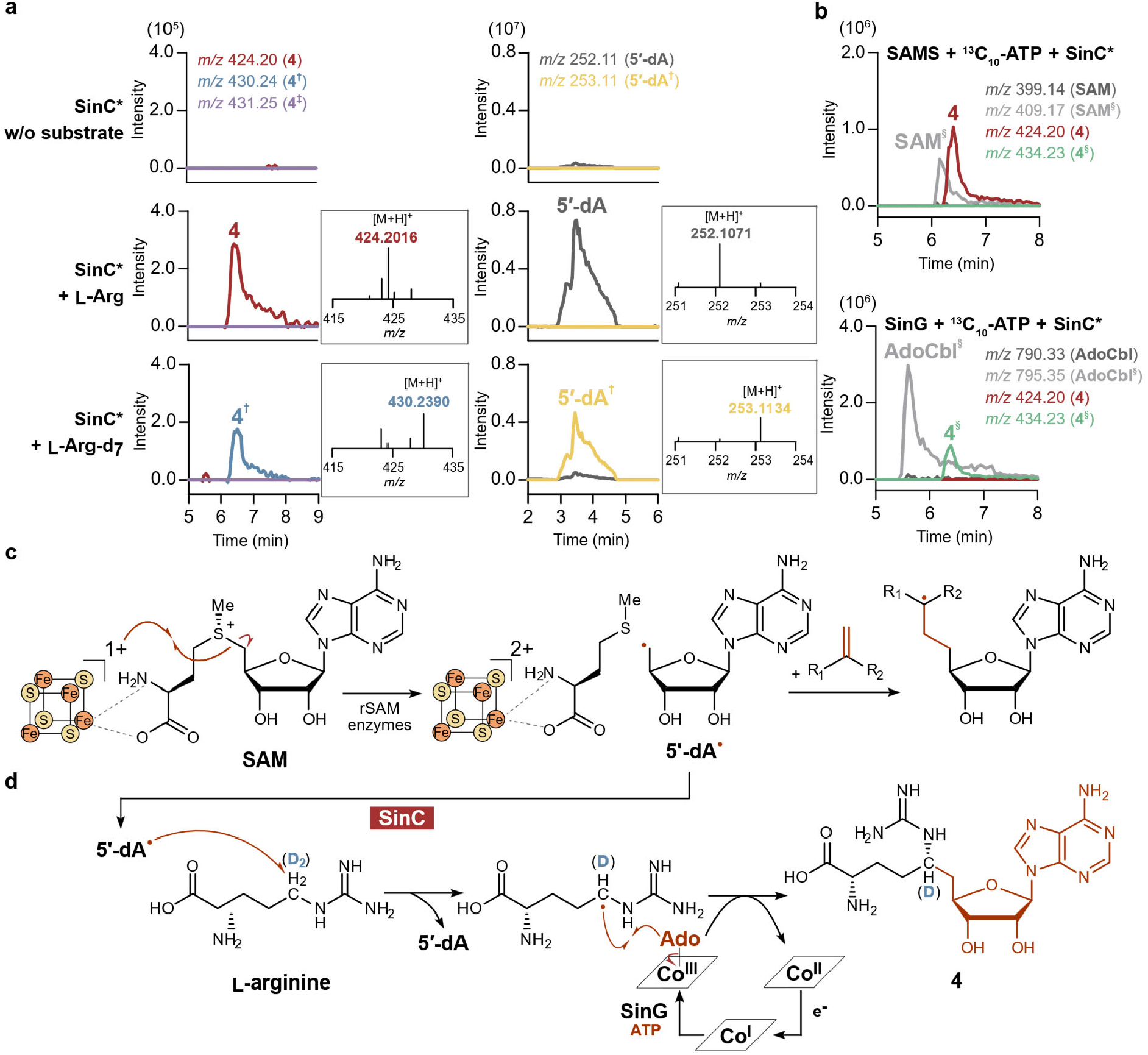
Mechanistic interrogations of SinC* with isotope-labeled substrates and cofactors. **a**, SinC**\*** reactions with L-arginine and uniformly deuterium-labeled L-arginine. Selected EIC(+) values are *m/z* values of **4** (red) and its deuterated forms **4**^**‡**^ (purple) or **4**^**†**^ (blue) corresponding to the loss of a single deuterium. High-resolution LC-MS of the product peaks are provided in the insets. **b**, Coupled reactions of SinC* and SAMS (top) or SinG (bottom) with uniformly ^13^C-labelled ATP. Selected EIC(+) values are *m/z* values of cofactors without (dark grey; SAM or AdoCbl, respectively) and with (light grey; SAM^§^ or AdoCbl^§^, respectively) ^13^C-ATP incorporation. Selected EIC(+)s for **4** is in red and **4** with ^13^C-ATP incorporation (**4**^**§**^) is in green. The *m/z* value of the byproduct 5′-dA shows a shift from *m/z* 252.11 (dark grey) to 253.11 (5′-dA^§^; yellow) when L-arginine-d_7_ is used.

The metabolic source of the adenosyl group was also unclear. Since SAM and adenosylcobalamin are typically used as 5′-deoxyadenosyl radical initiators, both cofactors used by a single enzyme have not been reported previously. Adenosylcobalamin-dependent enzymes typically regenerate the cofactor during their full catalytic cycle, yet radical derailment is known to occur in some systems.^50^ We ran coupled-enzyme assays to produce either ^13^C-SAM or ^13^C-adenosylcobalamin in the presence of SinC* to follow subsequent incorporation into **4**. The I137V variant of *Bacillus subtilis* SAM synthetase^51^ was independently expressed, purified, and added along with ^13^C-ATP and other necessary components to SinC* in-vitro reactions. In parallel, SinG was anaerobically purified and incubated with ^13^C-ATP, hydroxocobalamin, and the reductant dithiothreitol to produce ^13^C-adenosylcobalamin in SinC* assays that used a methyl viologen and NADPH reduction system (**Figure S9c**). Mass spectrometric analyses of the reaction products revealed ^13^C-label incorporation into **4** only when ^13^C-adenosylcobalamin was biocatalytically produced in vitro, and not when ^13^C-SAM was available (**Figb**). Thus, these data provide strong evidence that the adenosyl group in sinefungin is derived from adenosylcobalamin. Our mechanistic proposal aligns with bimolecular homolytic *S*_*H*_2 substitution reactions,^52^ akin to the use of methylcobalamin in B_12_-dependent radical SAM methyltransferases. To our knowledge, this is the only known enzyme to consume adenosylcobalamin via biosynthetic incorporation of the adenosyl group into a metabolite (**Figc, 6d**). This observed stoichiometric use of adenosylcobalamin also provides a satisfactory explanation for the presence of a second copy of the gene encoding ATP:Co(I)rrinoid adenosyltransferase in the *sin* BGC, since this pathway requires significantly more adenosylcobalamin than in a typical *Streptomyces* strain that uses the cofactor catalytically.

Upon final preparations of this manuscript, Ushimaru and Abe and coworkers reported the discovery of the sinefungin gene cluster in *S. incarnatus* NRRL 8090.^23^ The cluster was identified by sequencing the sinefungin-producing strains *Streptomyces griseolus* NRRL 3739 and *Streptomyces* sp. K05-0178 followed by comparative genomics analyses with *S. incarnatus*. The authors also reported the initial activities of several enzymes, including the key B_12_-dependent rSAM enzyme, by combining results attained from *S. incarnatus* and from homologous pathways in *S*. sp. K05-0178 and *Streptomyces cellostaticus* NBRC 12849. The two bodies of work nicely complement each other and are largely in agreement. Our work significantly expands upon the full metabolic profile observed in *S. incarnatus*, including the new derivatives sinefungin A and AA, and provides a more complete interrogation of all biosynthetic enzymes in the pathway.

## Conclusion

After decades of uncertainty, details concerning the biosynthesis of the SAM antimetabolite sinefungin are finally becoming clear. The unanticipated radical-mediated key C–C bond formed in the pathway broadens our perspective on how adenosylcobalamin can be employed by vitamin B_12_-dependent enzymes. In addition, the discovery of iterative PEARL enzymes acting on small-molecule scaffolds further highlight the plasticity of these catalysts for amino acid and peptide chemistry. All in all, these findings emphasize the value of basic science to unearth new enzyme activities that can serve as a foundation for future biotechnology applications to benefit human health.

## Acknowledgments

We would like to thank the following people for their generous gifts of materials: S.J. Booker for *isc* and *btu* expression plasmids pDB1282 and pBAD42-BtuCEDFB; A. Vagstad and J. Piel for SAMS expression plasmid pET28a-SAMS I317V; J.C. Escalante-Semerena for *Salmonella enterica* JE23613 (Δ*pduO*,Δ*cobA*, Δ*eutT*); and M. J. Smanski for *Streptomyces albidoflavus* J1074 and expression plasmid pSYH-synP. We are truly thankful to C.A. Townsend for the donation of his Coy anaerobic chamber. We also thank A.R. Buller for helpful discussions.

## Author Information

**Notes:** The authors declare no competing financial interest.

## Associated Content

The Supporting Information is available free of charge at XXX.

Additional experimental methods, data, Tables S1-S13 with accession numbers, primers, and NMR assignments, Schemes S1-S4 with workflows, and Figures S1-S49 (PDF)

