## Supporting Information for "Unconventional biocatalytic strategies orchestrate synthesis of the nucleoside analog sinefungin"

#### Experimental Procedures

##### Materials

All chemicals were purchased from Sigma-Aldrich, synthetic genes from Twist Biosciences, enzymes from New England Biosciences (NEB), and primers from Integrated DNA Technologies unless otherwise stated. *Streptomyces incarnatus* NRRL 8090 was purchased from Agricultural Research Service (ARS), USDA. *Streptomyces albidoflavus* J1074 (*S. albus*) and expression vector, pSYH-synP, were generous gifts from Dr. Michael Smanski. *Salmonella enterica* JE23613 (Δ*pduO*,Δ*cobA*, Δ*eutT*) was a generous gift from Dr. Jorge C. Escalante-Semerena. Plasmids pDB1282 (ISC operon expression) and pBAD42-BtuCEDFB (B12 uptake) were generous gifts from Dr. Squire J. Booker. Plasmid pET28a-SAMS I317V (SAM synthetase I317V) was a generous gift from Dr. Anna Vagstad and Dr. Joern Piel. Chromatographic data were processed using GraphPad Prism (v.5.04), and all figures were prepared using Affinity Designer (v.1.10.6.1665). Specific analysis software used for metabolomics and mass spectrometry are detailed in their respective sections.

##### Cloning and Plasmid Construction

The *sin* BGC (11.8 kb) was amplified from *Streptomyces incarnatus* genomic DNA extracted using the Monarch Spin gDNA extraction kit (NEB) via PCR with Q5 DNA polymerase (NEB). The expression vector pSYH-synP, which contains a *Streptomyces* synthetic promoter (synP),[1] was linearized by PCR using Q5 polymerase. All PCR reactions were pergormed according to the manufacturer's protocols with the following cycling conditions: initial denaturation at 98 °C for 30 s; 30 cycles of denaturation at 98 °C for 10 s, annealing at optimized temperatures for 20 s, and extension at 72 °C (time optimized for fragment length); followed by a final extension at 72 °C for 2 min. Vector and insert fragments were assembled using the NEBuilder® HiFi DNA assembly cloning kit (NEB). The resulting *sin* BGC expression construct further served as the template for top-down and bottom-up expression plasmid assemblies.

For protein expression in *E. coli,* gene sequences were codon-optimized and synthesized by Twist Biosciences (**Table S2**). Synthetic genes were cloned into linearized pET28 vectors (via PCR amplifications or NdeI/XhoI digestions) using the NEBuilder® HiFi DNA assembly cloning kit.

All assembled constructs were transformed into *Escherichia coli* TOP10 cells by electroporation for propagation. Plasmids were isolated using the GeneJET plasmid miniprep kit (Thermo Scientific) and sequence-verified by ACGT – DNA Sequencing Services or Plasmidsaurus. Primer sequences used in this study are listed in **Table S3**.

##### *Sin* BGC heterologous expression in *Streptomyces albidoflavus* J1074 (*S. albus*)

Media formulations were as follows: Mannitol Soy Flour (MS) agar (20 g/L mannitol, 20 g/L soy flour, 20 g/L agar; ISP2 (4 g/L yeast extract, 10 g/L malt extract, 4 g/L glucose, 20 g/L agar, pH

7.2); TSB (30 g/L Tryptic soy broth); 2×YT: 16 g/L tryptone, 10 g/L yeast extract, 5 g/L NaCl, pH 7.0); ISM3 (15 g/L yeast extract, 10 g/L malt extract, 20 g/L glucose, 1 g/L $MgSO_4 \cdot 7H_2O$, 0.3 g/L $FeCl_3 \cdot 6H_2O$, pH 7.0).

To prepare *S. albus* spores, TSB cultures were inoculated with a toothpick and shaken at 220 rpm for 2 days at 30 ºC, spread onto MS agar, and incubated at 30 ºC for 5–7 days. Spores were harvested in 2×YT, filtered through a 40-µm cell strainer, and adjusted to ~5 × $10^8$ spores/mL in 1:1 (v:v) 50% glycerol and 2×YT for storage at -80 ºC. For conjugation, freshly prepared or thawed spores (washed twice with 2×YT), were pre-germinated in 2×YT overnight at 4 ºC, heat-shocked at 50 ºC for 10 min, and incubated at 30 ºC for 3-4 h.

Expression plasmids encoding the *sin* BGC (including wild-type, deletion mutants, and homolog replacements) were introduced into *E. coli* ET12567/pUZ8002 via electroporation. A negative control plasmid encoding enhanced green fluorescent protein (eGFP) in the same pSYH-synP backbone was processed in parallel. After an hour of recovery at 37 ºC, cells were transferred to LB supplemented with apramycin (50 µg/mL), kanamycin (50 µg/mL), and chloramphenicol (25 µg/mL), and cultured for 22-24 hours at 37 ºC with shaking at 220 rpm. The *E. coli* cells were collected, washed twice with antibiotic-free LB, and resuspended in 2 mL LB for use as donor cells.

For intergeneric conjugation, activated *Streptomyces* spores (250 µL) and *E. coli* donor strains (500 µL) were mixed and plated on MS agar containing 10 mM $MgCl_2$. After overnight incubation at 30 ºC, plates were overlaid with 1 mL sterile water containing apramycin (1 mg/mL) and nalidixic acid (1 mg/mL). Exconjugants were allowed to develop and were then re-streaked on ISP2 agar containing apramycin (50 µg/mL) for secondary selection. Genomic integration was verified by PCR.

For expressions, confirmed exconjugants were inoculated in 10 mL TSB with apramycin (50 µg/mL) and cultured for 2 days at 30 ºC with shaking. Seed cultures (1 mL) were transferred to 100 mL ISM3 media with apramycin (50 µg/mL) in 500-mL Ultra-Yield flasks and incubated at 30 ºC with shaking at 220 rpm. Cultures were harvested at various time points for metabolite analysis.

**Metabolite Extraction and Chemical Analysis**

For metabolite extraction, 2 mL of cultures were centrifuged to separate the supernatant and cell pellet. Supernatants were partitioned three times with equal volumes of ethyl acetate (EtOAc). The aqueous phase was dried using a SpeedVac and subsequently extracted with 80% methanol (MeOH). Cell pellets were also extracted with MeOH. All MeOH extracts were dried, resuspended in 50% acetonitrile (ACN), and analyzed by high-resolution, high-pressure liquid chromatography−mass spectrometric analysis (LC–MS/MS).

LC-MS/MS was performed on a Thermo LTQ Orbitrap Velos mass spectrometer coupled to a Dionex RSLC HPLC system (Thermo Scientific) using a hydrophilic interaction chromatography

(HILIC) column (XBridge Amide, 3.5 µm, 130 Å, 150 × 4.6 mm) (Waters). Mobile phases solvent A ($H_2O$) and solvent B (95% ACN) both contained 0.1% formic acid and 5 mM ammonium formate. A linear gradient from 95% to 23% solvent B was applied over 8 min at a flow rate of 1.0 mL/min. Mass spectrometric detection was performed in positive mode electrospray ionization, and targeted MS/MS analyses were performed using higher-energy collisional dissociation (HCD) with a normalized collision energy of 20%. Raw data were visualized and processed using Thermo Xcalibur Qual Browser (v.4.5.474.0). For comparative metabolomics and the identification of unknown signals across data sets, Metaboseek (https://metaboseek.com/) was employed as previously described.[2]

$^{1}H$, $^{13}C$, $^{31}P$ and two-dimensional (2D) NMR spectra were acquired on a Bruker Avance NEO 600 MHz spectrometer equipped with a 5-mm BBO Prodigy probe, a Bruker Avance III HD 500 MHz spectrometer equipped with a 5-mm BBFO CryoProbe at the NMR Laboratory, University of Illinois Urbana–Champaign, or a Bruker Avance NEO 600 MHz spectrometer equipped with a 5-mm CryoProbe at the Minnesota NMR Center (MNMR), University of Minnesota.

**Isolation of compounds 1, 2 and 5–7**

Compounds **1** and **5** were purified from 2-day expression cultures in ISM3 of *S. albus* expressing *sin* BGC. Compounds **2**, **6**, and **7** were purified from 3-day cultures of *S. albus* expressing *sin* BGC-Δ*sinD*. In general, 1 L of expression culture was harvested by centrifugation, and the supernatant was partitioned three times with equal volumes of EtOAc. The aqueous layer was dried in vacuo and extracted with 1 L 80% MeOH. After removing precipitates, the extract was applied to an activated cation-exchange column packed with AmberChrom 50WX2 resin ($H^+$ forms, 100–200 mesh, 250 g, 270 × 45 mm) (DuPont de Nemours; purchased from Sigma-Aldrich). The column was washed with water, followed by sequential elution with 1 M and 2 M $NH_4OH$. Fractions containing target compounds were identified by LC–MS/MS and were then either directly purified by HPLC on an Atlantis Silica HILIC column (5 µm, 250 × 10 mm) (Waters) or first passed through a 50 g RediSep Gold C18Aq column (Teledyne ISCO) prior to HILIC purification. Detailed purification schemes are provided in **Schemes S1 &S2.**

**SinC* engineering design**

A collection of putative SinC homologs were compiled using the cblaster module of CAGECAT[3,4] with NCBI accession IDs WP_208903854, WP_208903148, WP_208903150, WP_208903151, WP_208903152, WP_208903153, WP_208903154, and WP_208903155 as queries. A Clustal Omega sequence alignment of SinC and 98 putative homologs was then made to create a SinC consensus sequence using the software Geneious 2025.2.2. An alphafold2[5] SinC model was used to cross-reference SinC surface-exposed residues that differed from the consensus sequence. Residues modeled with possible intra-molecular contacts were discarded as targets, as well as those near entry to, or within the putative active site. Amino acid substitutions were made based on the consensus sequence or other highly represented residues at each position, prioritizing soluble, charged and polar amino acids. The final engineered variant,

SinC*, had a total of 51 amino acid substitutions including a histidine tag replacing residues at the C-terminus (**Figure S8**).

**Overexpression and purification of SinC*- $His_6$**

Plasmid pET28-*sinC**-$CHis_6$ was co-transformed into *E. coli* LOBSTR BL21(DE3) cells with plasmid pDB1282 harboring the ISC operon[6] and pBAD42-BtuCEDFB for B12 uptake.[7] A single colony was grown overnight in 15 mL LB medium (Research Products International) supplemented with ampicillin (100 µg/mL), spectinomycin (50 µg/mL), and kanamycin (50 µg/mL) at 37 ºC with shaking at 220 rpm. The overnight culture was used to inoculate 1.5 L of Terrific Broth (TB) (Research Products International) in a 2 L flask. Cells were grown at 37 ºC with shaking at 220 rpm, and when an $OD_{600}$ of 0.4 was reached, ferric ammonium citrate (50 µM), L-cysteine (150 µM), hydroxocobalamin (OHCbl) (1.3 µM), and L-arabinose (0.2%) were added to the medium. The cultures were further grown at 37 ºC and 180 rpm until the $OD_{600}$ reached 1.5. The cells were then chilled on ice for 30 min, and ferric ammonium citrate (50 µM) and isopropyl β-D-1-thiogalactopyranoside (IPTG) (0.2 mM) were added for induction. The culture was then incubated at 16 ºC with shaking at 120 rpm overnight, and the cells were harvested by centrifugation (5,000 × g, 30 min, 4 ºC). The pellets were flash-frozen in liquid nitrogen and stored at -80 °C until further use.

Protein purification was performed under anaerobic conditions in a Coy vinyl anaerobic chamber (Coy Labs), and the samples were kept sealed during transfer outside the chamber. Cell pellets were resuspended in lysis buffer (50 mM HEPES, 300 mM NaCl, 10 mM imidazole, and 10% glycerol, pH 7.0) at 4 mL lysis buffer per gram of cells and supplemented with lysozyme (1 mg/mL). After incubating on ice for 30 min, the cells were lysed by sonication and clarified by centrifugation (17,000 × g for 50 min at 4 ºC). The supernatant was incubated with Ni-NTA resin (GoldBio) for 1 h at 4 ºC with a gentle rotation. The resin was washed with lysis buffer, followed by wash buffer (50 mM HEPES, 300 mM NaCl, 20 mM imidazole, and 10% glycerol, pH 7.0), and SinC* was eluted with elution buffer (50 mM HEPES, 300 mM NaCl, 250 mM imidazole, and 10% glycerol, pH 7.0). The eluted protein was concentrated using Amicon Ultra filters (3 kDa MWCO) (Merck) and desalted on a PD-10 column (GE healthcare) pre-equilibrated with storage buffer (50 mM HEPES and 10% glycerol, pH 7.0).

The purified protein was chemically reconstituted with [Fe–S] clusters under anaerobic conditions. Dithiothreitol (DTT) was added at a 100-fold molar excess relative to the protein, followed sequentially by $FeCl_3$ and $Na_2S$, each at a 10-fold molar excess. The reagents were rapidly combined to initiate cluster assembly. The protein solution was then added and mixed gently, and the reaction was incubated on ice for 3 h. Excess iron and sulfide were further removed using a PD-10 desalting column, and the protein solution was concentrated using Amicon Ultra filters.

**Overexpression and purification of $His_6$-SinG**

Plasmid pBAD-NHis6-*sinG* was transformed into *E. coli* LOBSTR BL21(DE3) cells and a single colony was grown overnight in 10 mL LB medium supplemented with ampicillin (100 µg/mL) at 37 ºC, 220 rpm overnight. The culture was inoculated into 1 L TB and grown at 37 ºC. Once the $OD_{600}$ reached 0.8, the culture was chilled on ice for 30 min and then induced with 0.05% L-arabinose. The induced cells were incubated at 16 ºC with shaking at 220 rpm for 18-20 h. Cells were harvested by centrifugation (5,000 × g, 30 min, 4 ºC), and the cell pellets were flash-frozen in liquid nitrogen and stored at -80 ºC until further use.

All purification steps for SinG were conducted under anaerobic conditions within a Coy vinyl anaerobic chamber. For any steps requiring transfer outside the chamber, samples were maintained in sealed containers. Cell pellets were resuspended in lysis buffer (50 mM HEPES, 300 mM NaCl, 10 mM imidazole, and 10% glycerol, pH 8.0) at a ratio of 4 mL per gram of wet cell weight, supplemented with 1 mg/mL lysozyme. After incubating on ice for 30 min, the cells were lysed by sonication and the lysate was clarified by centrifugation (17,000 × g, 50 min, 4 ºC). The supernatant was incubated with Ni-NTA resin for 1 h at 4 ºC with gentle rotation. The resin was washed sequentially with lysis buffer and wash buffer (50 mM HEPES, 300 mM NaCl, 20 mM imidazole, and 10% glycerol, pH8.0). The $His_6$-SinG was then eluted with elution buffer (50 mM HEPES, 300 mM NaCl, 250 mM imidazole, and 10% glycerol, pH8.0). The eluted protein was desalted on a PD-10 column pre-equilibrated with storage buffer (50 mM HEPES, 150 mM NaCl, 10% glycerol, pH 8.0) and concentrated using Amicon Ultra filters (10 kDa MWCO).

**Overexpression and purification of $His_6$-SAMS I317V**

Single colony of *E. coli* BL21(DE3) harboring pET28a-SAMS I317V was grown overnight in 10 mL LB medium supplemented with kanamycin (50 µg/mL) at 37 ºC, 220 rpm overnight. The culture was inoculated into 1 L TB and grown at 37 ºC until the $OD_{600}$ reached 1.5. Protein expression was induced by the addition of 0.1 mM IPTG, followed by incubation at 27 ºC and 220 rpm for 3 h. Cells were harvested by centrifugation (5,000 × g, 30 min, 4 ºC), and the cell pellets were flash-frozen in liquid nitrogen and stored at –80 ºC.

For protein purification, cell pellets were resuspended in lysis buffer (50 mM HEPES, 300 mM NaCl, 10 mM imidazole, 1 mM DTT and 10% glycerol, pH 8.0) at 4 mL lysis buffer per gram of cells and supplemented with lysozyme (1 mg/mL). After incubating on ice for 30 min, the cells were lysed by sonication (Qsonica) and the lysate was clarified by centrifugation (17,000 × g, 50 min, 4 ºC). The supernatant was collected and incubated with Ni-NTA resin for 1 h at 4 ºC with gentle rotation. The resin was washed sequentially with lysis buffer and wash buffer (50 mM HEPES, 300 mM NaCl, 20 mM imidazole, 1 mM DTT and 10% glycerol, pH8.0) and the protein was eluted with elution buffer (50 mM HEPES, 300 mM NaCl, 250 mM imidazole, 1 mM DTT and 10% glycerol, pH 8.0). The eluate was desalted via dialysis against storage buffer (50 mM HEPES, 150 mM NaCl, 1 mM DTT and 10% glycerol, pH 8.0) at 4 °C (one 2-h exchange followed by one overnight exchange), then passed through a pre-equilibrated PD-10 column. The protein solution was then concentrated using Amicon Ultra filters (10 kDa MWCO) and stored at -80 ºC.

**SinC* in-vitro assays**

Following [Fe–S] clusters reconstitution, the enzyme was used for in vitro activity assays. The reaction solution contained final concentrations of 2 mM SAM, 0.5 mM adenosylcobalamin (AdoCbl), 1 mM methyl viologen, 4 mM NADPH, 100 µM candidate substrate (including L-arg, D-arg and L-orn) and 20 µM SinC* in 50 mM HEPES (pH 7.0), with a total reaction volume of 100 µL. For adenosyl group incorporation tests, isotope labeled SAM and AdoCbl were prepared by the overnight in vitro reaction of SAMS I317V and SinG at the room temperature under anaerobic conditions, respectively. The reaction of SAMS I317V contains 10 mM $MgCl_2$, 2 mM ATP or $^{13}C_{10}$-ATP, 1 mM l-methionine and 100 µM SAMS I317V in the reaction buffer of 50 mM HEPES, pH8.0 with a total reaction volume of 50 µL. The reaction of SinG contains 10 mM $MgCl_2$, 1 mM ATP or $^{13}C_{10}$-ATP, 0.5 mM OHCbl, 10 mM DTT and 100 µM SinG in the reaction buffer of 50 mM HEPES (pH8.0) with a total reaction volume of 50 µL. After overnight incubation, the SinC* reaction components were added directly to the SAMS and SinG pre-reactions. The SinC* reaction mixtures were incubated at the room temperature under anaerobic conditions. After overnight incubation, reactions were quenched by the addition of MeOH to a final concentration of 80% (v/v). Precipitated protein was removed by centrifugation, and the supernatants were analyzed by LC–MS/MS.

**Overexpression and purification of $His_6$-SinF**

Plasmid pET28-N$His_6$-*sinF* was transformed into *E. coli* LOBSTR BL21(DE3) cells and a single colony was grown overnight in 10 mL LB medium supplemented with kanamycin (50 µg/mL) at 37 ºC, 220 rpm overnight. The culture was inoculated into 1 L TB and grown at 37 ºC until the $OD_{600}$ reached 1.5. The cells were chilled on ice for 30 min, induced with 0.2 mM IPTG, and incubated at 16 ºC, 220 rpm overnight. Cultures were harvested by centrifugation (5,000 × g, 30 min, 4 ºC), and the cell pellets were flash-frozen in liquid nitrogen.

For protein purification, cell pellets were resuspended in lysis buffer (50 mM HEPES, 300 mM NaCl, 10 mM imidazole, and 10% glycerol) at 4 mL lysis buffer per gram of cells and supplemented with lysozyme (1 mg/mL). After incubating on ice for 30 min, the cells were lysed by sonication and clarified by centrifugation (17,000 × g, 50 min, 4 ºC). The supernatant was incubated with Ni-NTA resin for 1 h at 4 ºC with gentle rotation. The resin was washed with lysis buffer followed by wash buffer (50 mM HEPES, 300 mM NaCl, 20 mM imidazole, and 10% glycerol), and $His_6$-SinF was eluted with elution buffer (50 mM HEPES, 300 mM NaCl, 250 mM imidazole, and 10% glycerol). The eluted protein was concentrated using Amicon Ultra filters (3 kDa MWCO), desalted on a PD-10 column pre-equilibrated with storage buffer (50 mM HEPES, 300 mM NaCl, 10% glycerol, pH 7.0), and dialyzed overnight at 4 ºC to remove residual imidazole. Purified SinF was stored at -80 ºC.

**SinF in-vitro assays**

Enzymatic assays were conducted in 50 mM HEPES (pH 7.0) containing 10 mM $MgCl_2$, 1 mM ATP, 100 µM of the candidate substrate (**1**, **2**, **4**, **5**, **6**, and **7**), and 20 µM SinF. Prior to the assays, SinF was desalted on a PD-10 column equilibrated with 50 mM HEPES to remove salts and glycerol. The reaction mixtures were incubated at 30 ºC overnight and quenched with an equal volume of 50% (v/v) MeOH, final concentration. Precipitated protein was removed by centrifugation, and the supernatants were analyzed using LC–MS/MS.

**Isolation of compounds 8 and 9 from a large-scale in-vitro assay**

Compounds **8** and **9** were purified from large-scale in-vitro assays of SinF using guanidinyl-sinefungin (**4**) and sinefungin (**1**) as substrates. Reaction conditions were the same as those described above. After overnight incubation, reactions were quenched by the addition of MeOH to a final concentration of 80% (v/v). Insoluble material was collected, re-dissolved in water, and residual precipitate was removed. The combined 80% MeOH-soluble and water-soluble fractions were applied to a Sephadex LH20 column (270 × 45 mm) (Cytiva) equilibrated with water. Fractions containing the target compounds were subsequently purified by HPLC on an Atlantis Silica HILIC column (5 µm, 250 × 10 mm) (Waters) (**Schemes S3 & S4**).

**Overexpression and purification of His6-SinR**

Plasmid pET28-NHis6-*sinR* was transformed into *E. coli* BL21(DE3) cells harboring pGro7 (Takara Bio), which encodes GroES/GroEL chaperones, by electroporation. A single colony was grown overnight in 10 mL LB containing kanamycin (50 µg/mL) and chloramphenicol (25 µg/mL) at 37 ºC with shaking at 220 rpm. The culture was used to inoculate 1 L TB. When the $OD_{600}$ reached 0.2, 50 µM 2,2′-dipyridyl was added to chelate iron. At $OD_{600} \approx 0.4$, L-arabinose (0.5 mg/mL) was added to induce GroES/GroEL expression. When cultures reached $OD_{600} \approx 1.5$, cells were chilled on ice for 30 min, and protein expression was induced with 0.2 mM IPTG along with supplementation of 1 mM $MnCl_2$. After overnight incubation at 16 ºC at 220 rpm, cultures were harvested by centrifugation (5,000 × g, 30 min, 4 ºC), and the cell pellets were flash-frozen in liquid nitrogen.

For protein purification, the pellets were resuspended in lysis buffer (50 mM Tris, pH 8.0, 300 mM NaCl, 10 mM imidazole, 10% glycerol) supplemented with 0.1 mM $MnCl_2$ at a ratio of 4 mL lysis buffer per gram of cells. Lysozyme (1 mg/mL) was added, and the samples were incubated on ice for 30 min before sonication. Lysates were clarified by centrifugation (17,000 × g, 50 min, 4 ºC), and the supernatant was incubated with Ni-NTA resin for 1 h at 4 ºC with rotation. The resin was washed with lysis buffer and subsequently incubated for 30 min at room temperature with wash buffer (50 mM Tris, 300 mM NaCl, 20 mM imidazole, 10% glycerol) supplemented with 5 mM ATP and 15 mM $MgCl_2$. After a final wash with wash buffer, $His_6$-SinR was eluted with elution buffer (50 mM Tris-HCl, 300 mM NaCl, 250 mM imidazole, and 10% glycerol). The eluted protein was dialyzed against 50 mM Tris (pH 9.0) twice for 3 h and once overnight. The following day, the protein was desalted using a PD-10 column equilibrated with 50 mM Tris (pH 9.0) to remove imidazole, salts, and glycerol prior to performing in-vitro assays.

**SinR in-vitro assays**

In-vitro reactions were performed in 50 mM Tris buffer (pH 9.0) containing 100 µM $MnCl_2$, 100 µM of the candidate substrate (including L-Arg, **4**, and **8**), and 20 µM freshly prepared SinR. The reactions were incubated at 30 ºC overnight and quenched with 80% MeOH. The MeOH-soluble fraction was dried, reconstituted in 50% ACN, and analyzed by LC–MS/MS.

**Overexpression and purification of $His_6$-SinD**

A single colony of *E. coli* LOBSTR BL21(DE3) carrying pET28-NHis$_6$-*sinD* was grown overnight in 10 mL LB containing kanamycin (50 µg/mL) at 37 ºC with shaking at 220 rpm. The culture was used to inoculate 1 L TB. When the $OD_{600}$ reached 0.2, 50 µM 2,2′-dipyridyl was added to chelate iron. Upon reaching $OD_{600} \approx 1.5$, cells were chilled on ice for 30 min and protein expression was induced with 0.2 mM IPTG together and supplemented with 1 mM $MnCl_2$. Cultures were incubated at 16 ºC at 220 rpm overnight, harvested by centrifugation (5,000 × g, 30 min, 4 ºC), and then flash-frozen in liquid nitrogen.

For purification, cell pellets were resuspended in lysis buffer (50 mM HEPES, 300 mM NaCl, 10 mM imidazole, 10% glycerol) at a ratio of 4 mL buffer per gram of cells, supplemented with lysozyme (1 mg/mL), and incubated on ice for 30 min before sonication. Lysates were clarified by centrifugation (17,000 × g, 50 min, 4 ºC), and the supernatant was incubated with Ni-NTA resin for 1 hour at 4 ºC with rotation. The resin was washed with lysis buffer followed by wash buffer (50 mM HEPES, 300 mM NaCl, 20 mM imidazole, and 10% glycerol), and $His_6$-SinH was eluted using elution buffer (50 mM HEPES, 300 mM NaCl, 250 mM imidazole, and 10% glycerol). The eluate was concentrated using Amicon Ultra filters (3 kDa MWCO), desalted on a PD-10 column equilibrated with storage buffer (50 mM HEPES, 300 mM NaCl, 10% glycerol, pH 7.0), and dialyzed overnight at 4 ºC to remove residual imidazole before storage at -80 ºC.

**SinD in-vitro assays**

SinD in-vitro reactions were carried out in 50 mM HEPES buffer (pH 7.0) containing 5 mM $MnCl_2$, 100 µM candidate substrate (including **8** and **9**), and 20 µM SinD. The reactions were incubated at 30 ºC overnight and quenched with 80% MeOH. The MeOH-soluble fraction was dried, reconstituted in 50% ACN, and analyzed by LC–MS/MS.

**Overexpression and purification of $SinB_1$ and $B_2$**

A single colony of *E. coli* BL21(DE3) carrying pACYCDuet-$His_6$-$SinB_1$ or pACYCDuet-$His_6$-$SinB_2$ was grown overnight in 10 mL LB containing chloramphenicol (25 mg/mL) at 37 ºC with shaking at 220 rpm. The culture was used to inoculate 1 L of LB which was grown at 37 ºC with shaking at 220 rpm until it reached $OD_{600} \approx 0.7$. The culture was chilled on ice in a cold room at 4 ºC for 20 min and protein expression was induced with 0.5 mM IPTG. Culture was incubated at 18 ºC, 220 rpm overnight, and harvested by centrifugation (4500 × g, 15 min, 4 ºC). Cell pellet from last step were resuspended in 15 mL of lysis buffer (20 mM Tris-Cl, 500 mM NaCl, 1 mM TCEP, 20 mM imidazole, 10% glycerol, pH 8.0) with protease inhibitor (ThermoFisher, #PIA32953) and gently rocked at 4 ºC for 20 min until resuspension reached homogeneous. Cells were sonicated on ice and lysate was clarified by centrifugation (4,5000 × g, 1 h, 4 ºC). The supernatant was incubated with Ni-NTA for 15 min with gentle rocking at 4 ºC and washed 3 times by a gradient of wash buffer (20 mM Tris-Cl, 500 mM NaCl, 1 mM TCEP, 25, 50, and 75 mM imidazole, 10% glycerol, pH 8.0). $His_6$-$SinB_1$ or $His_6$-$SinB_2$ was eluted with elution buffer (20 mM Tris-Cl, 500 mM NaCl, 1 mM TCEP, 300 mM imidazole, 10% glycerol, pH 8.0), buffer exchanged into storage buffer (20 mM Tris-Cl, 150 mM NaCl, 20% glycerol, pH 8.0) with a PD-10 column and concentrated with a Amicon Ultra filters (50 kDa MWCO). The concentrated $His_6$-$SinB_1$ or $His_6$-$SinB_2$ were further purified by size-exclusion chromatography using a Superdex 200 Increase 10/300 GL column (Cytiva) in SEC buffer (20 mM Tris-Cl, 500 mM NaCl, pH 8.0), concentrated, and stored at -80 ºC.

**Overexpression and purification of *S. incarnatus* alanyl-tRNA synthetase (siAlaRS)**

A single colony of *E. coli* BL21(DE3) carrying pET28a-siAlaRS-$His_6$ was grown overnight in 10 mL LB containing kanamycin (50 mg/mL) at 37 ºC with shaking at 220 rpm. The culture was used to inoculate 1 L of LB which was grown at 37 ºC with shaking at 220 rpm until it reached $OD_{600} \approx 0.7$. The culture was chilled on ice in a cold room at 4 ºC for 20 min and protein expression was induced with 0.25 mM IPTG. Culture was incubated at 16 ºC, 220 rpm overnight, and harvested by centrifugation (4500 × g, 15 min, 4 ºC). Cell pellet from last step were resuspended in 15 mL of lysis buffer (20 mM Tris-Cl, 500 mM NaCl, 1 mM TCEP, 20 mM imidazole, 10% glycerol, pH 8.0) with protease inhibitor (ThermoFisher, #PIA32953) and gently rocked at 4 ºC for 20 min until resuspension reached homogeneity. Cells were sonicated on ice and lysate was clarified by centrifugation (4,5000 × g, 1 h, 4 ºC). The supernatant was loaded onto HisPur™ Cobalt Resin (Thermofisher, # 89965) and washed 3 times by a gradient of wash buffer (20 mM Tris-Cl, 500 mM NaCl, 1 mM TCEP, 25, 50, and 75 mM imidazole, 10% glycerol, pH 8.0). SiAlaRS-$His_6$ was eluted with elution buffer (20 mM Tris-Cl, 500 mM NaCl, 1 mM TCEP, 300 mM imidazole, 10% glycerol, pH 8.0), buffer exchanged into storage buffer (20 mM Tris-Cl, 150 mM NaCl, 20% glycerol, pH 8.0) with a PD-10 column and concentrated with a Amicon Ultra filters (50 kDa MWCO).

**In-vitro transcription and aminoacylation of *S. incarnatus* $tRNA^{Ala}$s**

DNA template for the four $tRNA^{Ala}$s from *S. incarnatus* were prepared using Klenow fragment (NEB, #M0210S) and $tRNA^{Ala}$s were transcribed using HiScribe T7 High Yield RNA Synthesis Components (NEB, #E2040). Purified tRNAs were refolded by heating up to 92 ºC and slow cool-down to room temperature. Primers sequences used for $tRNA^{Ala}$s are listed in **Table S4**. Purified $tRNA^{Ala}$ (100 µM) were aminoacylated by 20 µM siAlaRS-$His_6$ in AARS buffer (50 mM Tris, 10 mM $MgCl_2$, 50 mM KCl, pH 7.5) containing 2 mM ATP, 1 mM DTT, and 1 mM L-Ala-

2,3-$^{13}C_2$ (MedChemExpress, #HY-N0229S9) at 37 ºC for 4 h. tRNAs were extracted using acid-phenol:chloroform, pH 4.5, and ethanol precipitated, and redissolved in 0.1 M sodium acetate, pH 5.2, and stored in -80 ºC for up to a week.

**Chemical synthesis of sinefungin A (6)**

To a dry 20 mL vial containing a magnetic stir bar was added sinefungin (**1**, 0.026 mmol, 1 equiv). 1M NaOH (64 µL) and water (536 µL) were added via syringe, followed by di-tert-butyl decarbonate (0.064 mmol, 2.5 equiv) in 1, 4-dioxane (600 µL). The reaction mixture was stirred at room temperature overnight. The crude mixture was then purified by HPLC to afford the Boc-protected sinefungin, (2S,5S)-6-((2R,3S,4R,5R)-5-(6-amino-9H-purin-9-yl)-3,4-dihydroxytetrahydrofuran-2-yl)-2,5-bis((tertbutoxycarbonyl)amino)hexanoic acid (**S1**). **S1** was then dissolved in DMF, followed by the addition of HATU (0.0273 mmol, 1.5 equiv) and DIEA (0.091 mmol, 5 equiv). After stirring for 10 minutes, L-Ala-OtBu (0.0273 mmol, 1.5 equiv) was added, and the reaction mixture was stirred overnight at room temperature. The reaction was quenched with 1 M HCl, extracted with ethyl acetate, and the organic phase was dried over $Na_2SO_4$, filtered, and concentrated. The resulting crude product was directly treated with trifluoroacetic acid (TFA, 1.0 mL) to remove the Boc and tert-butyl (tBu) protecting groups. After removal of TFA under reduced pressure, the crude residue was redissolved in water and purified by HPLC to obtain sinefungin A (**6**).

**In-vitro assays of SinB$_1$ and B$_2$**

20 µM of substrate was incubated with 5 µM of SinF, SinD, SinB$_1$, SinB$_2$, 50 µM of pre-aminoacylated tRNA$^{Ala}$s, 5 µM siAlaRS, 1mM L-Ala-2,3-$^{13}C_2$, 10 U Thermostable inorganic pyrophosphatase (NEB, #M0296), 10 U SUPERase•In™ RNase Inhibitor (ThermoFisher, #AM2694), in PEARL buffer (25 mM HEPES, 7 mM $MgCl_2$, 12 mM KCl, pH 7.5) containing 2 mM ATP at 30 ºC for overnight. The reactions were quenched with 2x volume cold acetonitrile. Proteins were removed by centrifugation and supernatant were lyophilized and redissolved in $H_2O$, 0.1% formic acid, and analyzed by LC-MS/MS. LC-MS/MS was performed on an Agilent 6545 LC/Q-TOF mass spectrometer coupled to an Agilent 1260 Infinity II HPLC system (Agilent) a Hypercarb™ Porous Graphitic Carbon HPLC Column (5 µm, 250 Å, 150 × 4.6 mm) (ThermoFisher, #35005-154630). Mobile phase were A ($H_2O$) and B (95% ACN), both with 0.1% formic acid. A linear gradient from 5% to 60% B was applied over 6 min at a flow rate of 1.0 mL/min. Mass spectrometric detection was performed in positive mode electrospray ionization, and targeted MS/MS analyses were performed using collision-induced dissociation (CID) at 10 V and 20 V.

**SinG adenosyltransferase complementation experiments in *S. enterica* JE23613 (*ΔpduO,ΔcobA,ΔeutT*)**

Plasmids pBAD-His$_6$–*sinG*, or pBAD-His$_6$–*sinG*_R137A were introduced into *S. enterica* JE23613 by electroporation. Transformants were plated on LB agar containing ampicillin (200 μg/mL). Single colonies were inoculated into 2 mL LB supplemented with ampicillin (200 μg/mL) and grown overnight at 37 ºC with shaking at 220 rpm.

The following day, 100 μL of overnight culture was inoculated into 10 mL LB supplemented with ampicillin (200 μg/mL) and incubated at 37 ºC at 220 rpm until the culture reached an $OD_{640}$ of 0.5–0.7. Protein expression was induced by adding 25% L-arabinose to a final concentration of 0.2%, followed by incubation for 3 h at 37 ºC and 220 rpm.

The cells were then washed twice with 2 mL of No-carbon E phosphate (NCE) medium supplemented with Wolfe's trace minerals, $MgSO_4$ (1mM), and ethanolamine hydrochloride (90mM) and resuspended in 1 mL of the same medium.[8] Cell suspensions were normalized to the same $OD_{640}$, and 4 μL of cells were added to 200 μL of the medium containing 0.2 μM OHCbl in wells of a 96-well plate. The plates were incubated at 37 ºC with shaking, and growth was monitored spectrophotometrically at 640 nm every hour for 48 h using a SpectraMax iD5 Multi-Mode Microplate Reader (Molecular Devices). Data acquisition and initial processing were performed using SoftMax Pro (v.7.1) software. The resulting growth data were further analyzed and plotted using GraphPad Prism (v.5.04).

#### Supplementary Figures

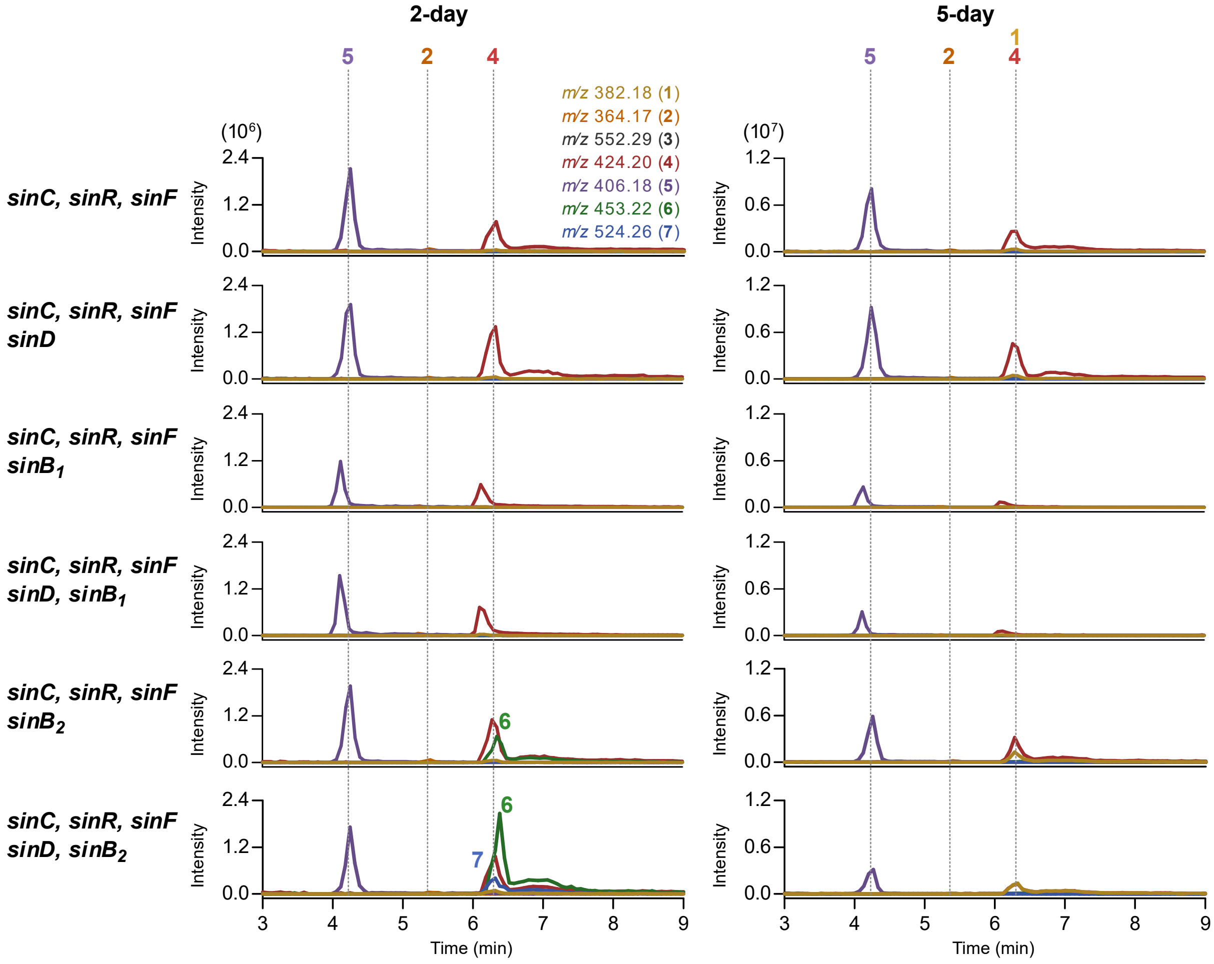

**Figure S1. Analysis of metabolites from bottom-up expression of *sin* BGC in *S. albus*. a, b,** Selected EIC(+) of extracts from S. albus harboring the indicated combinations of *sin* BGC genes and *sinE* after 2 days (a) and 5 days (b) of fermentation. Compound **6** is observed only when *sinB₂* was included, and **7** is produced when both *sinB₂* and *sinD* are present. Data are representative of three independent biological replicates.

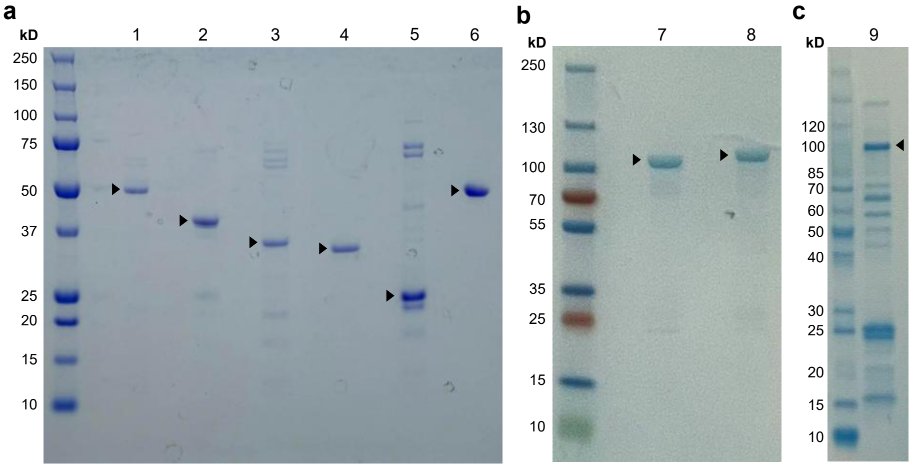

**Figure S2. SDS-PAGE of purified proteins for in vitro assays. a, b, c,** Purified recombinant proteins (2 µg/lane) were resolved on a 4–12% NuPAGE Bis-Tris gel (**a**) and a 4–12% Mini-PROTEAN® TGX™ protein gel (**b, c**) and visualized by Coomassie brilliant blue staining. Black arrowheads (▶) indicate the protein of interest in each lane. Lane 1: SinC*-CHis, 51.2 kDa; lane 2: NHis-SinR, 41.1 kDa; lane 3: NHis-SinD, 36.1 kDa; lane 4: NHis-SinF, 35.1 kDa; lane 5: NHis-SinG, 23.0 kDa; lane 6: NHis-SAMS I317V, 44.9 kDa; land 7: NHis-SinB1, 99.6 kDa; land 8: NHis-SinB2, 98.7 kDa; land 9: siAlaRS-CHis, 97.4 kDa.

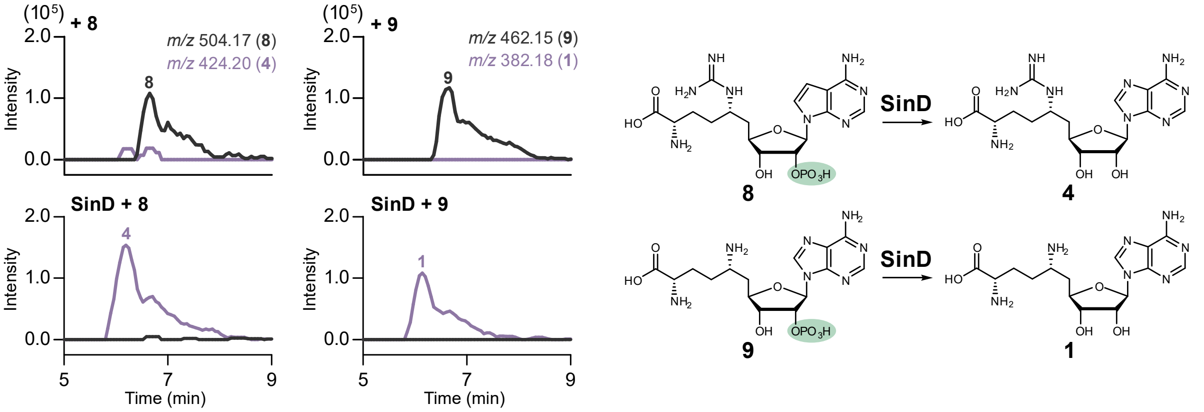

**Figure S3. Functional characterization of SinD.** LC-MS analysis of SinD in-vitro activity. SinD shows dephosphorylation activity toward **8** and **9**. Substrate and product traces are shown in black and purple, respectively.

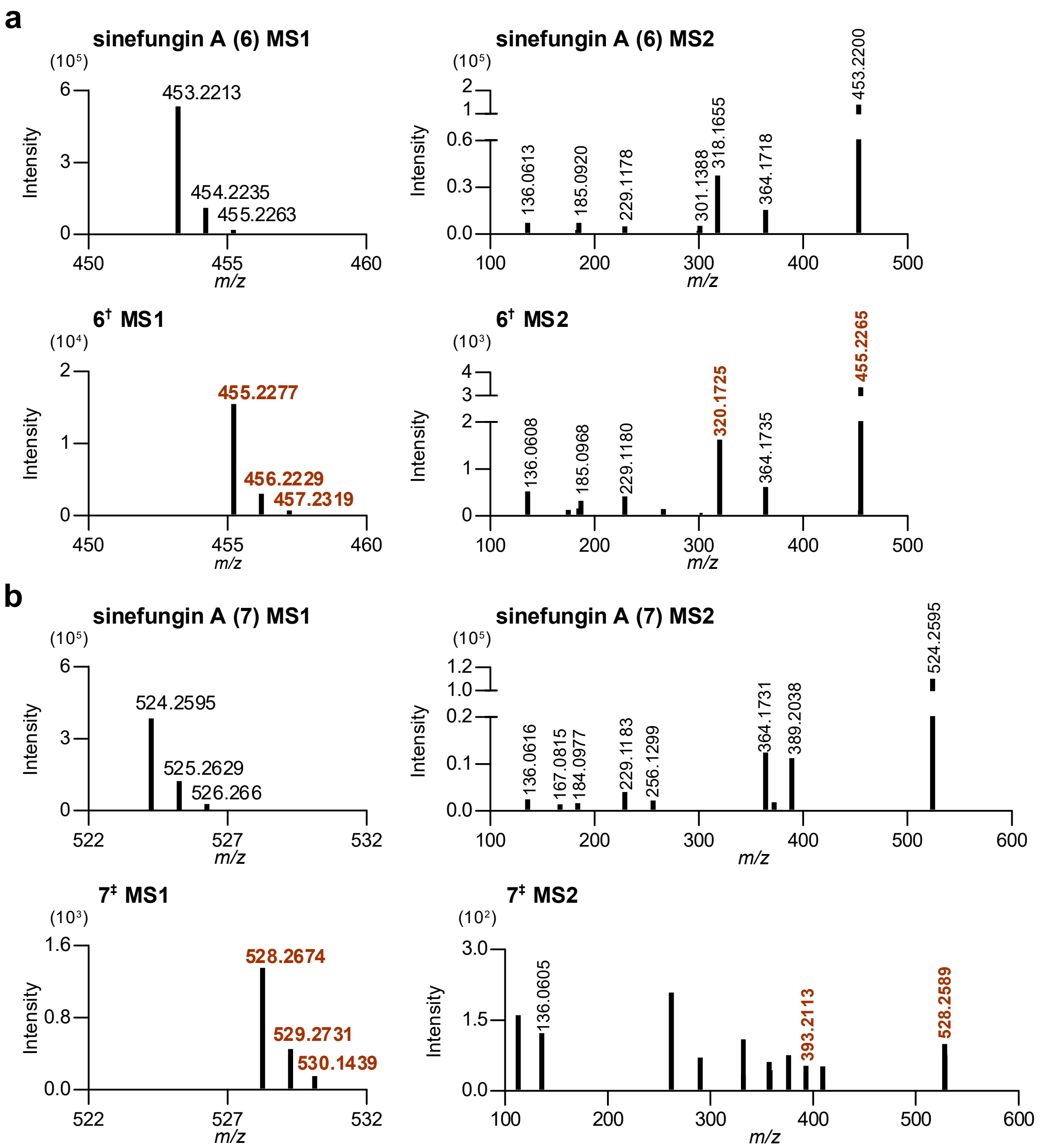

**Figure S4. High-resolution MS and MS/MS fragmentation spectra of products in SinB$_2$ reactions. a, b,** The top spectra correspond to the unlabeled reference standard of sinefungin A (**6**) and sinefungin AA (**7**). The bottom spectra illustrate the +2 or +4 Da mass shift in both the parent ion and characteristic structural fragments for **6**$^{\dagger}$ or **7**$^{\ddagger}$ (highlighted in red), confirming the selective incorporation of a single or two L-Ala-2,3-$^{13}C_2$ molecules.

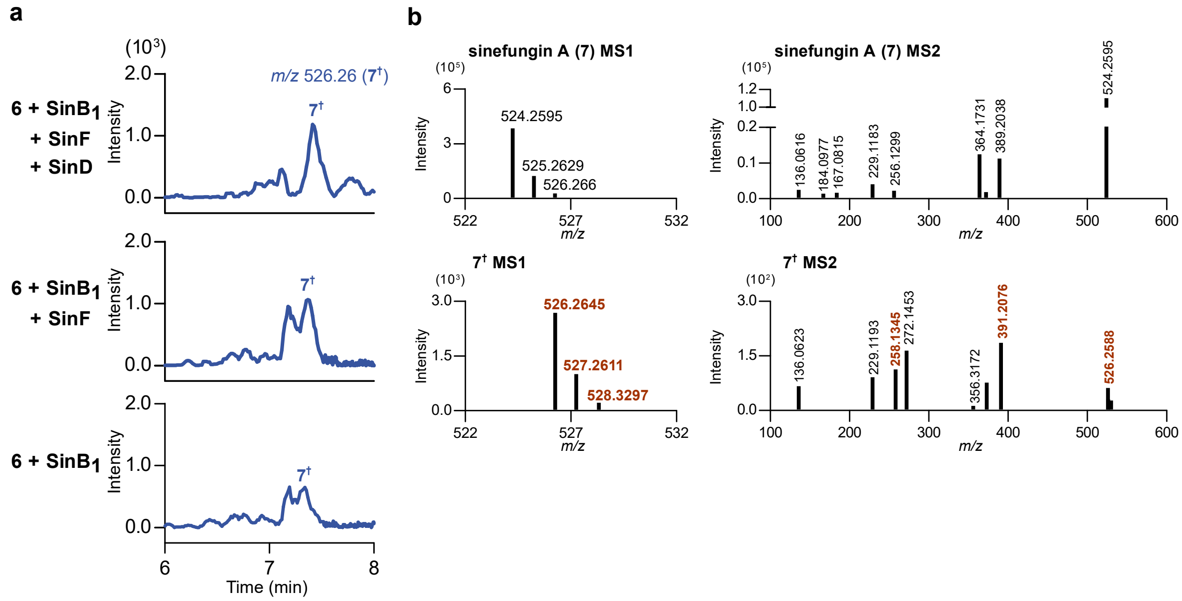

**Figure S5. In-vitro characterization of SinB1 activity with sinefungin A. a,** In-vitro reactions using **6** as the starting scaffold and $tRNA^{Ala}$s pre-charged with L-Ala-2,3-$^{13}C_2$ . Incubation of **6** with $SinB_1$ and with or without SinF and SinD yields formation of sinefungin AA with a single isotope-labeled Ala (**7**†). Addition of SinD and/or SinF does not significantly increase the yield of **7**† under these conditions. b, High-resolution MS and MS/MS fragmentation spectra confirm the identity of **7**†. The top spectra correspond to the unlabeled reference standard of sinefungin AA (**7**). The bottom spectra illustrate the +2 Da mass shift in both the parent ion and characteristic structural fragments (highlighted in red), confirming the selective incorporation of a single L-Ala-2,3-$^{13}C_2$ molecule.

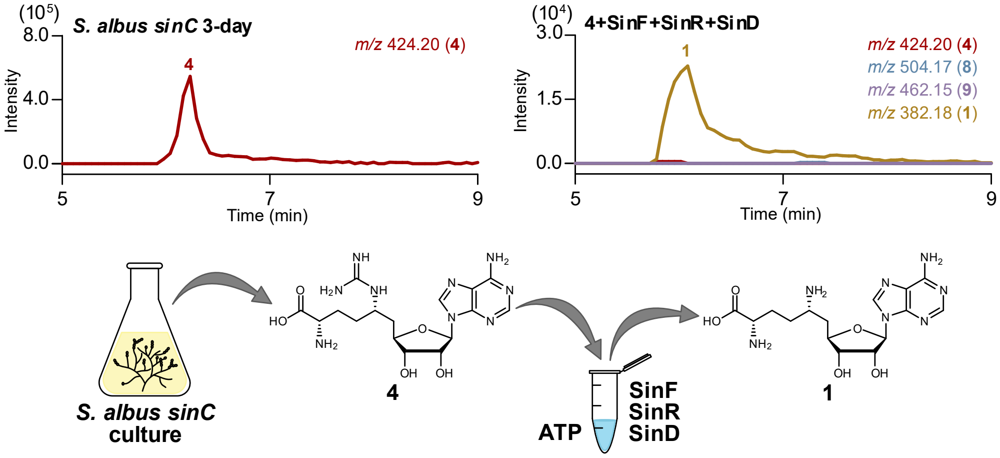

**Figure S6. A four-enzyme cascade to produce sinefungin.** The key intermediate **4** can be purified from the culture of *S. albus* expressing *sinC* alone. When **4** was used in an in-vitro coupled reaction with SinF, SinR and SinD, sinefungin (**1**) was successfully produced with near-complete conversion overnight.

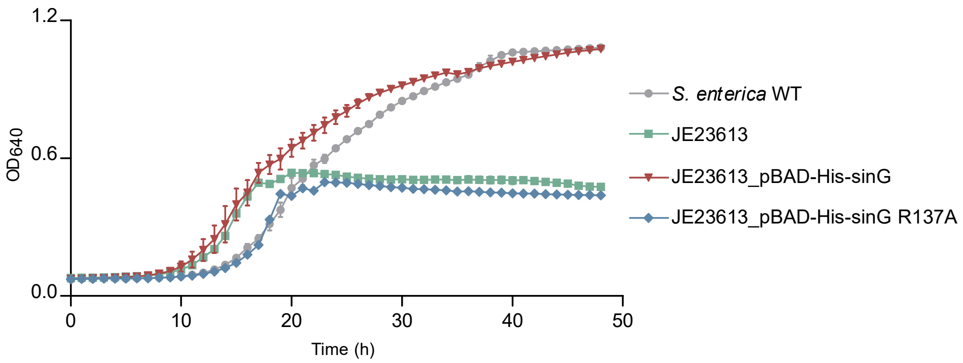

**Figure S7. Growth complementation of the *S. enterica* ΔACATs strain by *sinG*.** Growth curves of *S. enterica* wildtype (grey), the ΔACATs mutant JE23613 (green), and JE23613 strains expressing *sinG* (red) or the inactive *sinG* R137A variant (dark blue) in NCE minimal medium supplemented with Wolfe's trace minerals, $MgSO_4$ and ethanolamine hydrochloride. Expression of *sinG* and *sinG* R137A were pre-induced with 0.2% arabinose for three hours prior to inoculation into minimal medium. Data represent the mean ± standard deviation (s.d.) of three independent biological replicates.

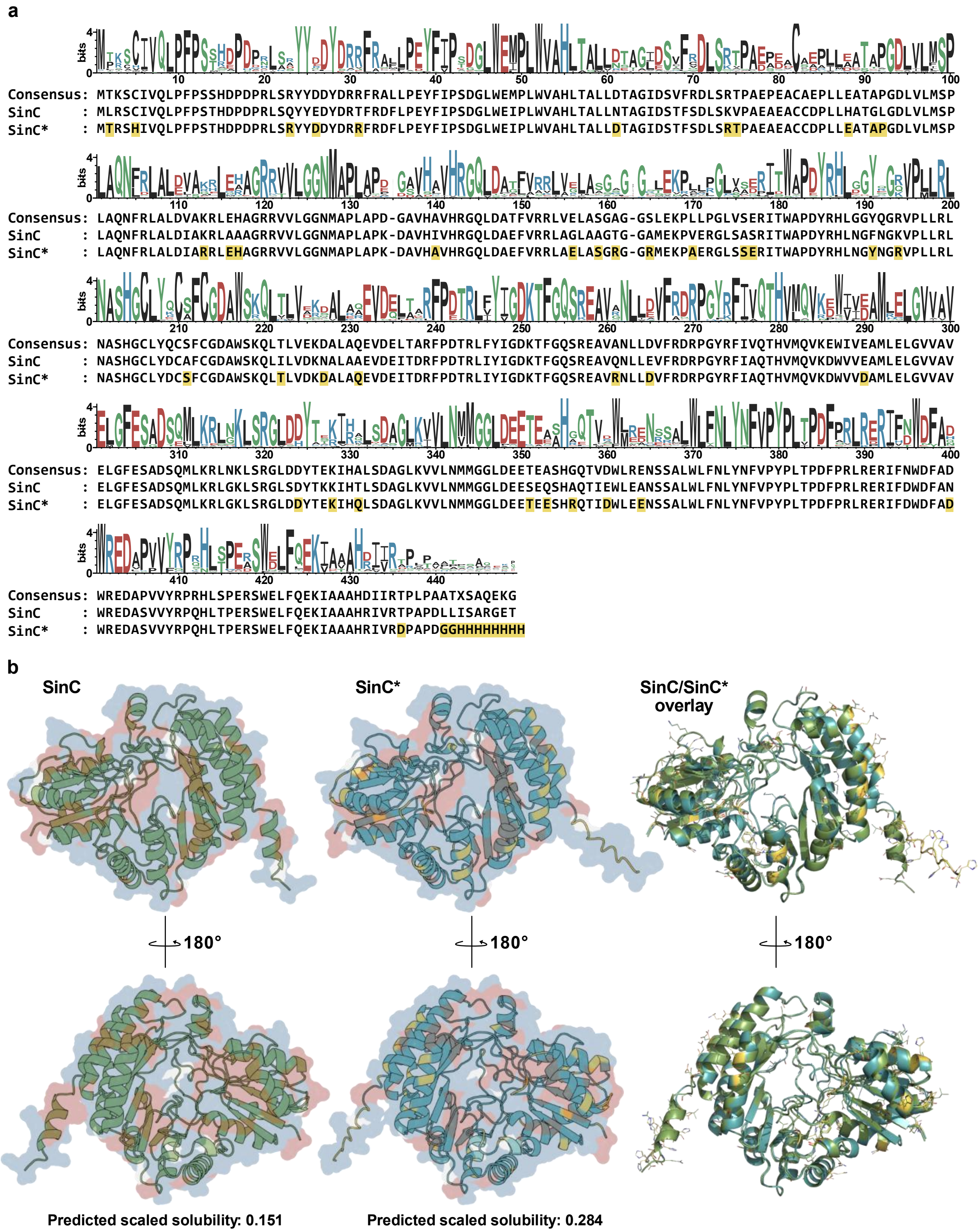

**Figure S8. Sequence and alphafold2-predicted structure alignment of SinC and designed SinC*. a,** Sequence logo generated from SinC and 98 putative homologs and sequence alignment of SinC, the SinC consensus, and the engineered SinC*. Amino acids highlighted in yellow in the sequence of SinC* are positions where residues were replaced with either the consensus amino acid or a highly represented polar or charged residues. b, AlphaFold2-predicted structures of SinC (left, green) and SinC* (middle, cyan), with an overlay shown on the right (RMSD= 0.260). The protein surfaces are colored according to amino acid hydrophobicity (Ala, Val, Leu, Ile, Met, Phe, Trp, Pro are in red; Arg, Lys, His, Asp, Glu, Asn, Gln, Ser, Thr, Tyr are in blue; Gly and Cys are in white). The bottom panels are rotated 180° horizontally from the top panels. Substituted residues in SinC* are labeled yellow. Predicted scaled solubility obtained by ProteinSol is indicated below each model, where higher values represent increased solubility.

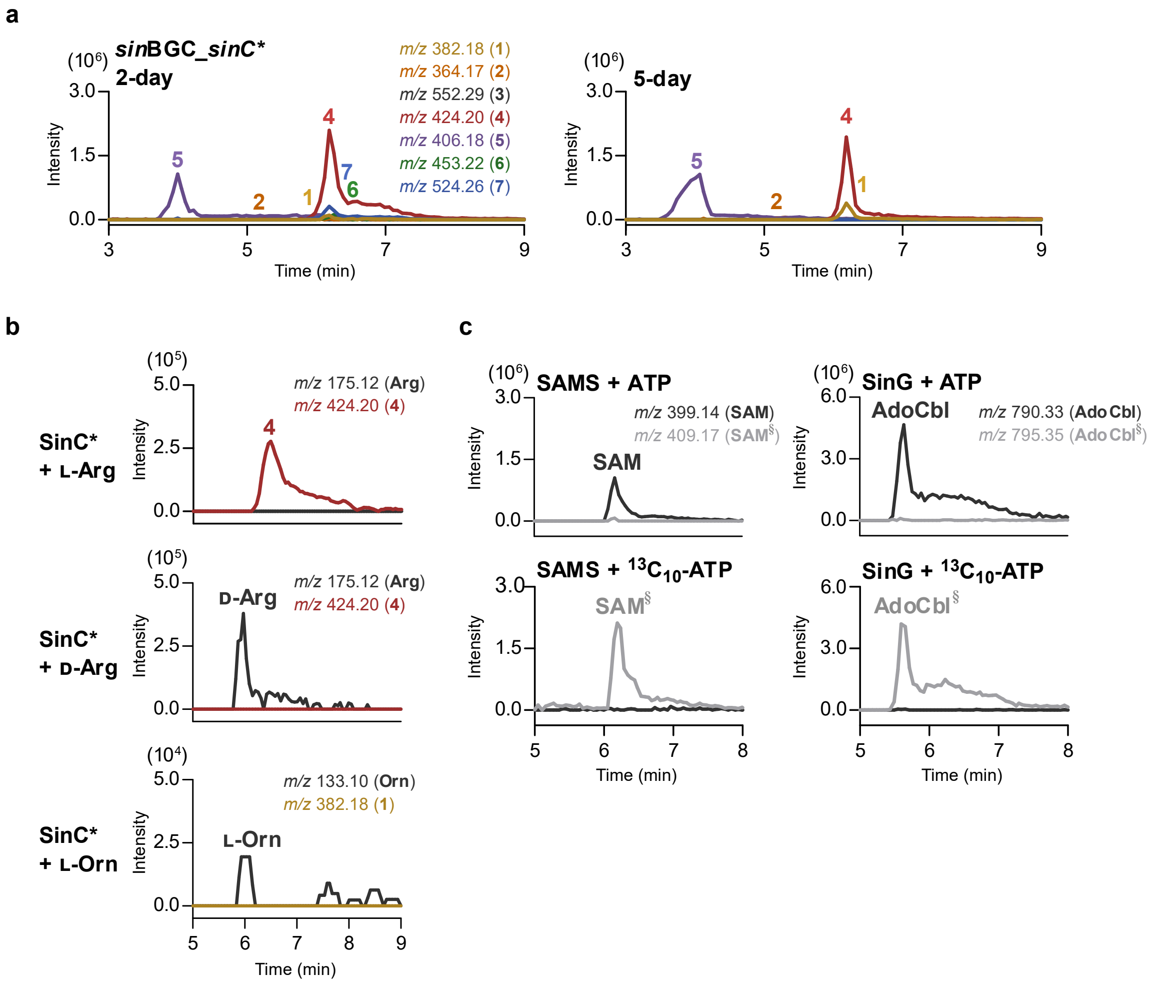

**Figure S9. Functional characterizations of SinC*. a**, LC-MS analyses of metabolite production in *S. albus* harboring the *sin* BGC with *sinC* replaced by the engineered *sinC**. Heterologous expression of the *sin* BGC with the *sinC** variant yields a metabolite profile identical to wildtype *sin* BGC. **b**, Substrate specificity of SinC*. SinC* converts L-Arg to 4 but displays no detectable activity toward D-Arg or L-Orn. c, Control reactions for the SinC* coupled-enzyme assays. SAMS and SinG utilize ATP and the isotope-labeled ATP to generate SAM and AdoCbl, respectively. The signifier (§) denotes the incorporation of the $^{13}C_{10}$ isotopic label.

#### Supplementary Tables

##### Table S1. Accession IDs for proteins encoded in the *sin* BGC from *S. incarnatus* NRRL 8089.

| **Gene name** | **Protein name** | **Accession (NCBI reference sequence)** | **Putative function** |
|---|---|---|---|
| *sinA* | SinA | AKJ15727.1 | Precursor peptide |
| *sinB$_1$* | SinB$_1$ | WP_208903155.1 | Peptide aminoacyl-tRNA ligases |
| *sinC* | SinC | WP_208903154.1 | $B_{12}$-dependent rSAM enzyme |
| *sinR* | SinR | WP_208903153.1 | Arginase |
| *sinD* | SinD | WP_208903152.1 | PHP domain-containing protein |
| *sinB$_2$* | SinB$_2$ | WP_208903151.1 | Peptide aminoacyl-tRNA ligases |
| *sinE* | SinE | WP_208903150.1 | MFS transporter |
| *sinF* | SinF | WP_208903149.1 | Aminoglycoside phosphotransferase |
| *sinG* | SinG | WP_208903148.1 | Cob(I)yrinic acid a,c-diamide adenosyltransferase |

**Table S2. DNA and protein sequences used in this study.**

| | | |
|---|---|---|
| ***sinA*** | **Description** | Potential precursor peptide. Codon-optimized for *E. coli.* |
| | **DNA Sequence** | ATGGCGCAGGCTTTTGGATTCTTTGGAGCCGCGTCAGAATCCGTGG<br>GGCAAGGCAAAGATCGGCCTGGGTATACTCCACCTGAGTTTGGCGA<br>ACTTCACGGCCCAGCGTTCGGCCGTGCATTACATGAACTGACTGCG<br>GAAATTCGACTGCAATTACTGGAAGGTCAGGTGGATCGGTCCTCAG<br>CAGAACCAGTTTTCGTAGCTGGAGGAGGTGAGCACCGTGTCGCAG<br>GTGATGCATGA |
| | **Protein Sequence** | MAQAFGFFGAASESVGQGKDRPGYTPPEFGELHGPAFGRALHELTAEI<br>RLQLLEGQVDRSSAEPVFVAGGGEHRVAGDA |
| ***sinB$_1$*** | **Description** | Lantibiotic dehydratase. Codon-optimized for *E. coli.* |
| | **DNA Sequence** | ATGGGAACCTCGGGTGCATCTTCGTCCGATAACCGCCTGCGCGACG<br>AAGATATGCATGAAGGCACGTCCGATGTCGGTGGCGGTTGGTCCTT<br>CGTACCAGTAGCCTTGTTAAGACACGCAGGATTTCCAGTCGCGCTG<br>CTTGATCGCTTTGCTGATGTGAGCGCTGCTGAAGAGGCGGATCTGC<br>TGTTGACTCGCGCCGATGCCGCCCGTCATTTGGCACAGGAAGTAAA<br>AGCCGCCCTGCGAGAGGCGCGCGTTGGTAGCCAGGGAGATATCTC<br>GTCAGGCGTGGGAATGCTGCGAGCGTTTGGAGATCAAGATCTGGC<br>ACGGCTGCGGGCTGATCTGGCGCCTTCGGCTTTAGAAATAGTGGAA<br>CGTTATCAGGACGCTGCACTCGGACTCGATCGCATGTGGATCCAGT<br>TTGAGCGCGATATGACGGAAGGTTTGGCACGTGCCCGTCATAATGT<br>GCAGGAACTGTTTCGGGATCCCGCGCTGAAACAAGTATTGTTGCTG<br>TCTAACGACGCCCGTTTTCCAGAATTTGCTGATTGGATTGATCGCGG<br>AAGTGGTGGCAGTCCGGGGCGTGCCCGACGTATGACTGATCTTCTC<br>GCCATGTACCTTCAACGGGTCACCACAAAGAACGAAACGCACTCC<br>CACTTTGGTCCTTTGGGTGTGGCTAGAGTTTCACCAGGTACATCTG<br>GGATCTCGTGGGCTGCAGGACCTTTACGTCGGACTGCCTTCTTGAC<br>GCATTGGGCAGGTGAACAGTTGGCCGAAACGTTTTCGCGTCGCCC<br>GGAAGCCTTTGAAGAAGTGCGTCCGCGCCGACGGCCACTTGCGTT<br>CGCCGAGGGAAGTCGGCTTGCCCTTTACGCGTTTGAATCAGTTACC<br>GGTATGCCTGATGGATGGAGATTTGTGCCTGTGGCAACCGCAACAT<br>TAGATGATGATCAGCTTTGGCTGTGGCAGCGCTGCGACGGTGAATG<br>CACCGTTGCACGCTTACGGAGAGCTTGGCATCTCCGGGAAGATGAA<br>GGGGATGCGCCTCTGCGAGGGTTCGACGATGTGCTCAATGAACTGA<br>TCGAAAAGGATTGGCTTGTGGGACGCTGGGAAATCCCCATTGGCTG<br>TCCTTATCCGCTGCGCGTGCTGACGGAACATCTTCAGAGTCCAGAT<br>GTACCCCGTACAGCAGATGGGGCTCGTGATCTGGCTACTGCCGCCT<br>GGTTTGAAGCTATGTTACAGGAATTCGCTACAGCGCCGATGGCCAA<br>TCGGGCCGCGTTGCTTACAAGCATGAAGGAGCACTTTCACCAGGTC<br>ACGGGAGCAGCAGCCAATCGAGCGGGCGGACGCCACTACGCTGAT<br>CGTTCCATCCTTTATGAGGAAGCGCATGGCCCGGTGCGCGATCTGC<br>GCATAGGCCAAGGAATAGCAGACCTGATAAGCCGCGAGTTAACTAT<br>CGTCTATGATCTCGCCTTGTTGGCCCCACGGTTACGTGTCCGACGTG<br>AACTGGAAGTGTTAGGACGCTGGGTTGAGAGACGTTTCGGAGCAG<br>AGGTAGACGTCCCACTTGAGCATTTCTACAATGGGTTTTACGCCGAT<br>CGTCCAGAACTGGCGCGTGAGTGTGAGGATATTGATCGCGAGATCA<br>CGGCGGTGGATCGTGAGATTACGGCACTGCTGCTGGATGGGTACGA<br>AGAGGGAACCGATGAAGCGGTAGTTGACCGCGAGCGCCTTGAGGA<br>GTTCCTTGCCCGTTTCCCTTCTTCGCCGCCAGCATTGTGTAACCCTG<br>ACGTTATGATAGCCTCGGAGAACGCTGGTGCGCTTGCACGTGGGGA<br>CCTGTTAGCAGTCGTTGGAGACTGTCACGGATTACGCGAGCTTCTG<br>ACACATTCCAGTTTCGCGCCGCTGATACAGACGGAAGCGCCGGAG<br>CTGCTGGATGAGGTATACAGCCGTTACCAGGACATGCTGGACTCAG<br>ACGAGTTGTTACTGGATCTTGCACGGGCACACCCAGACAAGACAG<br>GTGCCCAATTAGTGTACCCCTGCCCGGACTTGGAAGTTTATGGACTT<br>TCCGCGAAGACACGCGACCAGGTACTGCAGCCTCAGCAACTGTAC<br>GTAACTGTGCGCGGTGGACGTGCGCAACTCCGTGCTCAAGGCACG<br>GAGATGAGACTGAAGTTGTTAGCCCCCTTAGCTGGTGGTCCGTCCA<br>TACGACAAGATCCATTGAGCCCTTTCGCGTTCCCAAGACATTTCGG<br>CGGTATAGGTCTGGGCGCAGCTGATCATGACCGGTTGCCCCGTATCC<br>GATGTGGTCGGGTGGTACTTCAAAGAGCTCGTCGTCGCATACCTGC |

| | | |
|---|---|---|
| | | ACAAGCGTTCAGAGGTTGGAGTCCGGGAGTGCATCGTGGCAATGG<br>CGACGCGGCGGAATTCGCCGCAGGCCGTATGTTGCGCCGTGCGTTT<br>GACCTGCCTGAACAGGGCTTCGTGAAGATCCCAGGAGAACCGAAA<br>CCTGTCTACGTTGATTGGGAGAGTCCATTGCTGGTTAGACAGATTTT<br>CCGGTTAGTTCGCAAGACCGACTTGCCGGTCGACTTCTCGGAGATG<br>CTTCCCAGTAGAGAGCAATTATGGCTTGAGATCGGCGGTCAGCGCT<br>TTACCTCAGAACTTAGATGTGCGGTATTCTCTAGACGAGTCGGTTAA |
| | **Protein Sequence** | MGTSGASSSDNRLRDEDMHEGTSDVGGGWSFVPVALLRHAGFPVAL<br>LDRFADVSAAEEADLLLTRADAARHLAQEVKAALREARVGSQGDISS<br>GVGMLRAFGDQDLARLRADLAPSALEIVERYQDAALGLDRMWIQFE<br>RDMTEGLARARHNVQELFRDPALKQVLLLSNDARFPEFADWIDRGSG<br>GSPGRARRMTDLLAMYLQRVTTKNETHSHFGPLGVARVSPGTSGISW<br>AAGPLRRTAFLTHWAGEQLAETFSRRPEAFEEVRPRRRPLAFAEGSRL<br>ALYAFESVTGMPDGWRFVPVATATLDDDQLWLWQRCDGECTVARLR<br>RAWHLREDEGDAPLRGFDDVLNELIEKDWLVGRWEIPIGCPYPLRVLT<br>EHLQSPDVPRTADGARDLATAAWFEAMLQEFATAPMANRAALLTSMK<br>EHFHQVTGAAANRAGGRHYADRSILYEEAHGPVRDLRIGQGIADLISR<br>ELTIVYDLALLAPRLRVRRELEVLGRWVERRFGAEVDVPLEHFYNGF<br>YADRPELARECEDIDREITAVDREITALLLDGYEEGTDEAVVDRERLEE<br>FLARFPSSPPALCNPDVMIASENAGALARGDLLAVVGDCHGLRELLTH<br>SSFAPLIQTEAPELLDEVYSRYQDMLDSDELLLDLARAHPDKTGAQLV<br>YPCPDLEVYGLSAKTRDQVLQPQQLYVTVRGGRAQLRAQGTEMRLK<br>LLAPLAGGPSIRQDPLSPFAFPRHFGGIGLGAADHDRLPRIRCGRVVLQ<br>RARRRIPAQAFRGWSPGVHRGNGDAAEFAAGRMLRRAFDLPEQGFV<br>KIPGEPKPVYYVDWESPLLVRQIFRLVRKTDLPVDFSEMLPSREQLWLEI<br>GGQRFTSELRCAVFSRRVG |
| ***sinC**** | **Description** | Engineered SinC (rSAM enzyme) with C terminal his tag. Codon-optimized for *E. coli.* |
| | **DNA Sequence** | ATGACCCGTTCTCATATTGTTCAGCTGCCTTTTCCAAGTACTCATGA<br>CCCCGATCCTCGTCTGAGCCGCTATTACGATGATTATGACAGACGCT<br>TTCGTGACTTTTTGCCGGAGTACTTCATACCTTCCGACGGCCTCTGG<br>GAAATCCCACTGTGGGTCGCCCACCTTACGGCTCTGCTGGATACCG<br>CTGGTATAGATTCAACATTTTCCGATTTGTCACGCACGCCGGCTGAA<br>GCCGAGGCGTGTTGTGATCCCTTGCTGGAAGCAACAGCGCCAGGG<br>GACCTGGTGCTTATGTCGCCGCTCGCGCAAAACTTTCGTCTGGCAC<br>TTGACATCGCCCGTCGTCTCGAACATGCCGGCCGGAGAGTCGTCCT<br>GGGTGGTAACATGGCACCCCTTGCCCCTAAAGATGCAGTACACGCT<br>GTTCACCGAGGTCAGCTGGACGCCGAATTTGTGAGAAGATTGGCC<br>GAGCTTGCATCGGGTCGCGGAGGACGCATGGAAAAGCCAGCGGAA<br>CGGGGGCTTTCATCTGAACGTATTACATGGGCTCCAGACTATCGGCA<br>TCTGAATGGGTATAATGGTCGTGTTCCGTTATTGCGACTGAATGCCT<br>CACACGGATGCTTATACGACTGTTCGTTCTGTGGTGACGCCTGGTC<br>CAAACAGTTAACATTGGTCGACAAAGATGCGCTTGCTCAGGAAGTT<br>GACGAAATTACAGATCGTTTTCCGGATACCAGACTCATTTATATTGG<br>CGATAAAACATTCGGCCAAAGTCGGGAGGCAGTTCGAAATCTGTTA<br>GATGTTTTTCGGGACCGTCCCGGCTATCGGTTTATTGCTCAGACCCA<br>CGTTATGCAGGTTAAAGACTGGGTGGTCGATGCTATGCTGGAATTG<br>GGAGTTGTTGCCGTGGAACTGGGCTTCGAGAGCGCTGACTCGCAG<br>ATGCTGAAACGTCTGGGCAAACTCTCCCGCGGGCTTGACGATTATA<br>CGGAAAAAATTCACCAACTTTCAGATGCCGGTTTAAAAGTGGTGCT<br>GAACATGATGGGGGGTCTGGACGAGGAGACCGAGGAATCCCATCG<br>CCAAACCATCGATTGGCTGGAGGAAAATTCTTCGGCCCTCTGGCTG<br>TTTAATCTTTACAACTTTGTCCCGTATCCCTTGACTCCAGATTTTCCT<br>CGGCTCCGAGAACGCATTTTCGACTGGGATTTTGCAGATTGGCGAG<br>AAGATGCTAGCGTTGTATATCGTCCGCAGCATCTGACTCCGGAACGT<br>TCGTGGGAGCTGTTTCAAGAAAAAATCGCAGCGGCCCATAGAATAG<br>TTCGTGATCCTGCCCCAGATGGTGGCCACCACCATCACCATCATCAC<br>CATTAA |
| | **Protein Sequence** | MTRSHIVQLPFPSTHDPDPRLSRYYDDYDRRFRDFLPEYFIPSDGLWEI<br>PLWVAHLTALLDTAGIDSTFSDLSRTPAEAEACCDPLLEATAPGDLVLM<br>SPLAQNFRLALDIARRLEHAGRRVVLGGNMAPLAPKDAVHAVHRGQ |

| | | |
|---|---|---|
| | | LDAEFVRRLAELASGRGGRMEKPAERGLSSERITWAPDYRHLNGYNG<br>RVPLLRLNASHGCLYDCSFCGDAWSKQLTLVDKDALAQEVDEITDRF<br>PDTRLIYIGDKTFGQSREAVRNLLDVFRDRPGYRFIAQTHVMQVKDW<br>VVDAMLELGVVAVELGFESADSQMLKRLGKLSRGLDDYTEKIHQLSD<br>AGLKVVLNMMGGLDEETEESHRQTIDWLEENSSALWLFNLYNFVPYP<br>LTPDFPRLRERIFDWDFADWREDASVVYRPQHLTPERSWELFQEKIAA<br>AHRIVRDPAPDGGHHHHHHHH |
| ***sinR*** | **Description** | Arginase. Codon-optimized for *E. coli* |
| | **DNA Sequence** | ATGACCCAGCAGCTCCCGGACTTCGAGGCCTTACAACACTCGGTGA<br>AGCCAGAGGACTTCGAGACCGACCCATTGCCGGGCTGGGCTGGTG<br>TTGCAACGCTGTTCGGCGCGCCTGCCACGTCCATGTCTCAAGTGGA<br>CGTCGGAACCAATGGCTGGGTAGTTAGCGGAGTGCCGTTCGACGC<br>GACTGCGTCAAGTCGCCCTGGTGCCGCGGAAGGGCCCAGAGCAAT<br>CCGGCAGGCTTCACTGGTTTTCAGCTCGTATCTGACTTCTCTGGGC<br>GAATGGAGAATGCTCGACACACGCTCAGGTCACACCTTCCGTTACC<br>GGGCCCCCAAGGTAGCCGACGCGGGCGACCTGCACGTCTACCCCA<br>CTAACACTGTTCGTACATTCCAGGCGGTAGCGGCCGAGAGTCGGAG<br>ATTAAGTCAGGCGGCCCCGCATTTAATTTTCCTCAACGGAGACCAC<br>AGTGCCACCTTCCCAACATTCGCCGGATTCCGTGCGGGTCGCTTGG<br>ACGGTGGGGCCGAGCGTGTTGGATTCATCAACATTGATCACCACTT<br>CGACTTCGGGAACTGGTCGACGCTGCACGGCCCACTTTACCACGGT<br>TCGAACTCACGCCGGATCAGCGAGCTGCCGGGCGTGCGTGCAGAA<br>GATATCGCCTTCGTCGGCGTTGGCGATGTGACGAAGTACGACCAGT<br>TCCGGGGCCTTCTGGAACAAGGTTTCCACGTTGTCCCGGGTGTACA<br>GATCTTAAAAGAGGGAGCAGCAACGGCGCTGAGACCGGTCATTGA<br>GCACCTGGGCGAGCGTTGCGACACAGTATACGTGTCTCTGGACATA<br>GACGTGCTGGACTCGGCCGTCTCCACCGGTACGGGCCATGTAACAA<br>TGGGCGGGATCGACACCGGAGCACTTTTAGACATATACGAGGAGTT<br>ACGTGCCTTACCGGTTGGAGCAGTTGACATCGCTGAGGTCGCACCT<br>CGCTATGACCCCAGCGGCAGCACAGCACAGATAGCCGCTCACTTAT<br>TATTCAATTTCTTGTTCAGAACGGAGTCGGCCGCTAGTGGACTGGC<br>GCCGTGGTCGGACGGTGGGGCGGTGGCCCTCGTGGTGA |
| | **Protein Sequence** | MTQQLPDFEALQHSVKPEDFETDPLPGWAGVATLFGAPATSMSQVDV<br>GTNGWVVSGVPFDATASSRPGAAEGPRAIRQASLVFSSYLTSLGEWR<br>MLDTRSGHTFRYRAPKVADAGDLHVYPTNTVRTFQAVAAESRRLSQA<br>APHLIFLNGDHSATFPTFAGFRAGRLDGGAERVGFINIDHHFDFGNWS<br>TLHGPLYHGSNSRRISELPGVRAEDIAFVGVGDVTKYDQFRGLLEQGF<br>HVVPGVQILKEGAATALRPVIEHLGERCDTVYVSLDIDVLDSAVSTGT<br>GHVTMGGIDTGALLDIYEELRALPVGAVDIAEVAPRYDPSGSTAQIAA<br>HLLFNFLFRTESAASGLAPWSDGGAGGPRGDR |
| ***sinD*** | **Description** | PHP domain-containing protein. Codon-optimized for *E. coli* |
| | **DNA Sequence** | ATGCCTGGAGTCCCAGCCATTTTAGCCGGCCCTGTTGATTTACACTT<br>ACATACACGTCGTAGCGATGGGGATGAATCCCCGGCTGAAGTTGCC<br>CGCCGTTGCCTGGCAGCAGGTTTGCGTGTGGCATCCGTGACAGATC<br>ATAATACGATGGCTGGTATTGCTGAATTTACTGGTACCGTTGGGCGC<br>AGCATGACCGTCGTACCTGGTTCAGAAGTGACCGCACGCTGGCGA<br>GGTGAAGAGGTCCATTGTCTGGCTTATTTTATTGACCCTGCAGATCG<br>CCGATTTCAACGGCAATTACAACGCATTCATGATGCGGATCTTGATT<br>GGTGGCGAGCTTGGGCCGAACGTGTTAGACATATTGGCGTCCCTCT<br>GACATGGGAAGATATAGCTCAACGGATTGGCGGGGATCGTATAGCA<br>TATCCAGGTGATTATTTAGCACTCTTAGCTGAAGCGGCCCGAGATGA<br>TCCACGCTTTGCGGAATATGGACCTACCGAGCATGAACGCCTGGCG<br>ACCGATTGGTGTAGACCTGGTAGACCTCTTCACGTACCTGAACCCT<br>GGATGCCGGATATAATGGATGTTCTGAGCTGGATAGCTGATGCTGGC<br>GGTGTAGCAGTATTGGCACATCCAGCTCGGGTGTTAGATGTTTCTGA<br>TTCCGCACAATGTCGTGCCTTATTGGAGCCTTTAGTTGCTGCTGGCT<br>TAGCGGGCTTAGAAGTATGGACCTCATGGCATACGCCTGCTGAATC<br>CGCAGGTCTTGCGCGTATATGCGCAAGTTTGGGGATGGCTGCAACT<br>GCTGGGAGCGATTACCACGGCGTGCGCGTGAAAAGTTGGGTAGCA<br>GCGCCAGGACGTCTTCCATCTCTGCCGCCCGAGCCTATGGCGGTTT<br>TAGATGCCTTGTGTGATCGGCGTCCGACTCCTTTAAAACCAGAAGC<br>AGAGCTGTAA |

| | | |
|---|---|---|
| | **Protein Sequence** | MPGVPAILAGPVDLHLHTRRSDGDESPAEVARRCLAAGLRVASVTDH<br>NTMAGIAEFTGTVGRSMTVVPGSEVTARWRGEEVHCLAYFIDPADRR<br>FQRQLQRIHDADLDWWRAWAERVRHIGVPLTWEDIAQRIGGDRIAYP<br>GDYLALLAEAARDDPRFAEYGPTEHERLATDWCRPGRPLHVPEPWM<br>PDIMDVLSWIADAGGVAVLAHPARVLDVSDSAQCRALLEPLVAAGLA<br>GLEVWTSWHTPAESAGLARICASLGMAATAGSDYHGVRVKSWVAAP<br>GRLPSLPPEPMAVLDALCDRRPTPLKPEAEL |
| **$sinB_2$** | **Description** | Lantibiotic dehydratase. Codon-optimized for *E. coli.* |
| | **DNA Sequence** | ATGCGACCTACTGCTCACCCAACAGAAGCACGCGGCGGTACGATG<br>ACGGCCGTCATGACTCCTAGTGACCCACACAGTCAGGCTGCTGCAG<br>CAAGCAGACCACCCGAGGTAGGGAGAGCATGGTCCTTAGTTCCTG<br>TTGTGGTTTTACGTCAAGCTGGTTACCCGATGGAGTTACTGGACCCT<br>TTACTGGCACCTGACGCGGCAGAGGAAGCAATGAGTCTTCTTGATA<br>GCAGAGACCGACTCGCTGGTTTGGCAGATGAGGTTAAGAAGCTTC<br>TTCGCAAGCACCAAATCACTGGTGGCCAGCAGTTGAGCAGTCGCG<br>CGGGGCAACTCCGTCCGGTCCCAGAGGCAGAGTTGAACGGGGCTC<br>TTGCCGGACTGCCTAATGACGCCAGCGCAACTTTAAGAAATTACCA<br>AGAGGCGGCGATCCGTCTGGGAGACCGGTGGGCAGCATTTGCTGT<br>TCGTCACAGAGAACGGCTGGCAGCAGCTCGTCTCGCAGTAACGGA<br>GACCTTCTCCGACCCATCATTGAGACAAGTACTCCTGCTGTCAAAC<br>GACGCGGCGTACCCGGAGTTTTCTTCGTGGTTAGACTCATTCCACG<br>GCGAGGCAGGGCCACGGACCCGCAAGATGACCGACCTGCTGGCAA<br>TGTACTTACAACGCGTGACGACAAAGAACGAAACCCATTCGCACTT<br>TGGTCCTATAGCTGTAGGGCGCGTCGTTCGGGACACACCGGGTATA<br>GCTTGGGTACCTCGCCAACCTTTGGAGCGCCGATCCTATTTCGCGC<br>ATTGGGCAGCTGAGAGACTTGCCGCGGCCGCCGGACAAACGCCTG<br>GATTGGGCGACCACGTCCGACCGCGTCGTCGCCCGTTGGCTTTCCT<br>TCGTGATGGACGCATCGATCTGTACGCCTTCACGACTCGCGACGGT<br>TTGGACATTGACTGGGATTTTCAGCACCTGGGTGGAGCGGAAGTGT<br>CCCCCAGTGAAGAGTGGTTATGGCAACGCTGCAATGGAGATCGGA<br>CGGTCTTAGAGTTGCGGGCTGAGTGGTCAGAACGTCACGGGCCCC<br>GCCATCAATCCCTCTTCGACGAGGTGTTGCGCCGTTTGACAGAGCG<br>CGAGTGGGTGATTGCCGAGTTCGAGATTCCCGTAGGCGGATCACAG<br>CCTTTAGCAGCACTCCGCGATATGCTTCCTGCCGCTGTGGAGCCCG<br>CCCGCCAAATCTTGGACGTTATCACACGTTTCGAGCAAGACCTTGT<br>CCGGTTCTCCGCTCTGCCGCTTGCTCAGCGCCCTGCACTTCTGTCT<br>GACGCCAAGCACCGTTTCGAGTCGGTTACGCGATCGCCTGCCAACA<br>GAAATTCGGGGTTACACTACGCCGACAGAAGCATCTTATTCGAAGA<br>GGCCCACGGTGAATTGCGCGACCTGACCATTGGTCCTGACATAGCT<br>CGCTTCATTACTGACGAGTTGAGCATAGTCTACGACACAGTTCTGG<br>CTGGCCCACGTCTGCGTATGCGCCGCGAAACGGCTATCCTCACACG<br>TTGGGTTACAGATCGGTTCGGAACAGGTACCGAGGTGGACTTGGAT<br>CGATTGTACGCTGACTTCTTCGACGATCGCGATCGTCTTGCCGCTGA<br>ATGTGCCGTTGTTGATGCCGAGTTAGCCGACCTGGACGAGGCGATG<br>ACCGGTGCAGTCCTGGGTGACGATACCGAGCGGCGTGAAGTTGTA<br>ATTCCGCGAGAGCGTGTTGAAGAGGTTCTTTCGGCGTATCCGGACA<br>GTCCCGCAGCCGTCTGCAACCCTGATATCCTCTTCGCGGCGACATC<br>GCAGGAAGCCTTACGGCGGGGAGACTTTACGGCCGTAATAGGGGA<br>CTGCCACGCTGTTCGTGAAGTCATCACGCATACGAGTTTCGGCCCA<br>TTAGTCCAAGAGACCGCCCCTCAGCTTTTACCTGAGGTATACCGGG<br>GTTACCTCTCACTGTTAGATGACGACGAAGTGCTTGTAAATTTGAG<br>CAGAGGCCATCCGGACAAAACATCAACGCAACTGTGTTACCCATGC<br>TATGACCTTGAAGTTTATGGTCGTTCCGCTCAGACGCGCGATAAAGT<br>GCTCCAACCATCTCAGCTCTACGTCGTGGTTCGTGATGGCCGCTTA<br>GAGCTGCGTGCCAGAGGAGTCGAGGGTCGGTTACGTCTTATGGCG<br>CCCCCGGCAGGTGGCCCGTCCATCGTACAAGACCCATTGAGTCCAT<br>TTAGTTTCCCACGTCATTTCGGTGGAGTAGGCCTTCGCGCTAGTGCA<br>CTTGACCATGTGCCGCGCATTCGTTGTGGTCGCGTAGTTCTTCACCG<br>AGAGACTTGGCGCATCCCTACCGCTCGCCTGCGCGGGTTGGCCCTC<br>TCTGGGGATAGAGTAACGGCCGACGATGCTGCGCAGTTCTTAACTA<br>TATGCCGTTTACGCGCCGAGCATGGTCTCCCACAGCAAGTCTTCGT<br>TAAGGTGCCTGGTGAGCCTAAGCCTTTGTACGTTGATTGGGACGCA |

| | | |
|---|---|---|
| | | CCCTTATTGGTACGTCAGCTCTGTCGCCTGGCCCGGAAAACCGACG<br>GCACATTAGAGATATCAGAGATGCTCCCGGGCCGCGATCAAAGATG<br>GCTTGAAACCGGGGCTCGGCGTTATACTGCCGAACTGCGGTGCGCG<br>GTTTTCAGTAGTGGCAGACGCCGGTAA |
| | **Protein Sequence** | MRPTAHPTEARGGTMTAVMTPSDPHSQAAAASRPPEVGRAWSLVPVV<br>VLRQAGYPMELLDPLLAPDAAEEAMSLLDSRDRLAGLADEVKKLLR<br>KHQITGGQQLSSRAGQLRPVPEAELNGALAGLPNDASATLRNYQEAA<br>IRLGDRWAAFAVRHRERLAAARLAVTETFSDPSLRQVLLLSNDAAYPE<br>FSSWLDSFHGEAGPRTRKMTDLLAMYLQRVTTKNETHSHFGPIAVGR<br>VVRDTPGIAWVPRQPLERRSYFAHWAAERLAAAAGQTPGLGDHVRP<br>RRRPLAFLRDGRIDLYAFTTRDGLDIDWDFQHLGGAEVSPSEEWLWQ<br>RCNGDRTVLELRAEWSERHGPRHQSLFDEVLRRLTEREWVIAEFEIPV<br>GGSQPLAALRDMLPAAVEPARQILDVITRFEQDLVRFSALPLAQRPALL<br>SDAKHRFESVTRSPANRNSGLHYADRSILFEEAHGELRDLTIGPDIARFI<br>TDELSIVYDTVLAGPRLRMRRETAILTRWVTDRFGTGTEVDLDRLYAD<br>FFDDRDRLAAECAVVDAELADLDEAMTGAVLGDDTERREVVIPRERV<br>EEVLSAYPDSPAAVCNPDILFAATSQEALRRGDFTAVIGDCHAVREVIT<br>HTSFGPLVQETAPQLLPEVYRGYLSLLDDDEVLVNLSRGHPDKTSTQL<br>CYPCYDLEVYGRSAQTRDKVLQPSQLYVVVRDGRLELRARGVEGRL<br>RLMAPPAGGPSIVQDPLSPFSFPRHFGGVGLRASALDHVPRIRCGRVV<br>LHRETWRIPTARLRGLALSGDRVTADDAAQFLTICRLRAEHGLPQQVF<br>VKVPGEPKPLYVDWDAPLLVRQLCRLARKTDGTLEISEMLPGRDQRW<br>LETGARRYTAELRCAVFSSGRRR |
| ***sinF*** | **Description** | Phosphotransferase. Codon-optimized for *E. coli.* |
| | **DNA Sequence** | ATGTCGGAAAGATTAGATACCTTGACCGAACTGGCAGAACGGTGG<br>GGCATTGATGCACCCACACTGGCTGGGAGCGGTCTCGAATTTAGTG<br>TTTATCGTGCACGCAGTCGTGATGGCGATCCGGTTGCCTTGCGCGTA<br>GCTCATAGAAGATTTGACAGTAATGCGAACGATCCGGCGGTCGATA<br>CGCGCGCTCTCCTTGTTCAAGAGTATCGTATTACACGTCACCTCGCT<br>GCGCATGGTTTCCCAGTAGCTGAACCGGTCAAGTTATTCTTAAGTG<br>ATGACTCTTCCTCTCCCGATGTTTTGTTGAGTCGGTATGTTCCGGAC<br>GATGGTGCTCCTTTAGATTGTTTTGCACTGGGCCAGCTTCTGGCAA<br>GACTGCACTTATTACCGCCCCCTCGTCTCGACTTAGTGGCTAGCGA<br>AGGAATGTCAACTCCTAAAGTTATTGCAACACGTATTCAAAGACGG<br>TGGGCAGAGGTCGGTTGCCACGTGATGGATTGGCCTAAACCCCCTG<br>GTCATGCACATTTAGCTCGCTTATTAGCGGGCGTATCAGGAAATTCC<br>TTACTTCACCTGGATGTAAGATCCGATAATATACGACGTGAAGATGG<br>TGAGATTACGGCTTTATTAGACTGGTCCAATGCCCTGCCAGGTAATT<br>CAGCAATGGAGTTTGGTCGGTTAGTTGAATATGCACTCTACGCAGA<br>AAATGAGTTAGATATGGCTACCTTACGCGCCGGATATGATACAGTGT<br>GTCAAGCACCGCCCGAGGGTGGTAAAGAGCTTCTGGTGTGTAGAC<br>TTGATGCGGCTGTCATGCTGGCTCTGGTGTTCTTGTCAGAGGCACC<br>GGACCCCAAACGGGGTGCTCTCGCGGCCGATCATGTACGTGAATTA<br>GGGGCTATGGTAGCCGCTTTAGAGGCATAG |
| | **Protein Sequence** | MSERLDTLTELAERWGIDAPTLAGSGLEFSVYRARSRDGDPVALRVAH<br>RRFDSNANDPAVDTRALLVQEYRITRHLAAHGFPVAEPVKLFLSDDSS<br>SPDVLLSRYVPDDGAPLDCFALGQLLARLHLLPPPRLDLVASEGMSTP<br>KVIATRIQRRWAEVGCHVMDWPKPPGHAHLARLLAGVSGNSLLHLD<br>VRSDNIRREDGEITALLDWSNALPGNSAMEFGRLVEYALYAENELDM<br>ATLRAGYDTVCQAPPEGGKELLVCRLDAAVMLALVFLSEAPDPKRGA<br>LAADHVRELGAMVAALEA |
| ***sinG*** | **Description** | ATP:corrinoid adenosyltransferase. Codon-optimized for *E. coli.* |
| | **DNA Sequence** | ATGGGCCATATGGTTAACTTGACCCGCATCTACACAAAGACAGGCG<br>ATGATGGGACAACCGCGTTGGGAGATATGTCTCGGGTTTCTAAGAC<br>CGACTCGCGCATCGCCGCGTATGCAGATGTTGACGAGGCTAACAGT<br>GCGATTGGGCTGGCGATTGCAGTTGGTGAATTAGACGCTGAGACGA<br>GCGCCCTTCTTCTTCGTATTCAGAACGACTTATTTGATGTCGGCGCC<br>GATCTTTGCACTCCACTGGAAGGCGCTGACCCGAACGCACCTGCTC<br>TCCGGGTGGAGCCCGCTTACATCAGATTCTTGGAAGAACAGTGCGA<br>TCGTCGGAACGAGGGCTTGACAACGTTGAGAAGTTTCATCCTTCCC<br>GGCGGATCTCGGGGAGCCTCTTACCTGCATCTTGCTCGCACCGTAG<br>TAAGACGTGCTGAACGCTCGACATGGATCGCTATAGAGCAGCATGG |

| | | |
|---|---|---|
| | | CGACACCATGAACAAGACCACGGCTCACTATTTAAACCGGCTTAGC<br>GATCTGCTGTTCATTCTGGCCAGAGCTGCTAATTGCGCTAGAGGTG<br>ACATCCCTTGGCAACCCGGGGCGAATCGTTAA |
| | **Protein Sequence** | MGHMVNLTRIYTKTGDDGTTALGDMSRVSKTDSRIAAYADVDEANS<br>AIGLAIAVGELDAETSALLLRIQNDLFDVGADLCTPLEGADPNAPALR<br>VEPAYIRFLEEQCDRRNEGLTTLRSFILPGGSRGASYLHLARTVVRRAE<br>RSTWIAIEQHGDTMNKTTAHYLNRLSDLLFILARAANCARGDIPWQP<br>GANR |
| **siAlaRS** | **Description** | *S. incarnatus* alanyl-tRNA synthetase. Codon-optimized for *E. coli.* |
| | **DNA Sequence** | ATGGAAAGTGCGGAAATTCGCCGCCGTTGGCTTAGCTTCTTCGAAG<br>AACGTGGCCACACCGTGGTCCCCAGCGCGTCGCTTATAGCTGATGA<br>CCCAACCCTCTTGCTTGTTCCAGCAGGAATGGTCCCCTTCAAGCCG<br>TACTTCTTAGGAGAGGTCAAACCACCTTTCCCGCGAGCTACAAGTG<br>TGCAGAAATGTGTACGAACCCCCGACATTGAAGAAGTGGGCAAAA<br>CCACAAGACACGGTACCTTCTTTCAAATGTGTGGAAACTTTTCCTT<br>TGGAGACTATTTTAAGGAAGGAGCAATCAAGCTTGCTTGGGAGTTA<br>CTCACTACGCCTCAGGACAAGGGTGGCTACGGCCTTGACCCCGAG<br>CGTTTATGGATAACGGTGTACAAAGATGACGACGAAGCCGAAAGA<br>ATTTGGCACGAAGTGGTGGGTGTACCTAAAGAGCGGATCCAACGTC<br>TCGGCATGAAAGACAATTACTGGTCCATGGGTGTGCCTGGTCCTTG<br>TGGGCCTTGTAGTGAGATCAATTACGACCGAGGACCTGAGTTCGGG<br>GTGGAAGGCGGTCCCGCCGTCAACGATGAGCGTTATGTCGAGATCT<br>GGAATTTAGTTTTCATGCAATATGAGAGAGGTGAAGGTACAGGGAA<br>GGATAATTTCGAGATTCTGGGAGAGTTGCCGAGTAAGAATATCGATA<br>CTGGTTTGGGCCTGGAGCGCCTGGCCATGATCCTTCAGGGCGTGCA<br>GAACATGTACGAGATCGACACCTCGATGGCTGTTATAGACAAAGCC<br>ACGGAACTTACGGGCGTCAGATACGGCGACGCGCATGACAGCGAT<br>GTTAGTTTACGCGTCGTGACTGACCATATGCGGACGGCCACCATGC<br>TGATAGGTGATGGGGTCACCCCTGGGAACGAAGGACGCGGCTATGT<br>TTTACGTCGAATCATGCGTCGCGCTATCCGCAACATGCGATTACTTG<br>GGGCAACAGGACCGGTAGTGAAGGACTTACTTGATACAGTGATACA<br>AATGATGGGGCGCCAATACCCGGAGCTCGTCACAGACCGCGAACG<br>TATTGAAAAGGTCGCAATCGCAGAGGAAAATGCGTTCTTGAAGAC<br>ATTGAAGGCTGGGACTAACATTCTGGACACAGCGGTAACTGAGAC<br>AAAACAAGCGGGCGGTACAGTCCTGTCCGGGGACAAGGCTTTCCT<br>GCTGCACGACACGTGGGGCTTTCCCATAGACCTGACTTTGGAGATG<br>GCTGCGGAACAGGGGCTGTCGGTAGATGAGGAAGGATTCCGGAGA<br>CTCATGAAAGAGCAACGCGAGAGAGCAAAGGCGGACGCACAGGC<br>AAAGAAGACCGGTCACGCGGACATGGGCGCCTATCGTGAAATCGC<br>AGACACTGCCGGGGAGACTGACTTTATTGGATACTCCGACACCGAG<br>GGAGAATCTACTATAGTTGGTCTTTTGGTCGATGGGATCTCTTCCCC<br>TGCAGCCACCGAAGGTGACGAGGTGGAAGTCGTTTTGGACCGGAC<br>ACCATTTTACGCAGAAGGGGGGGGTCAAATAGGTGATACCGGTCGA<br>ATCAAGGTCGACTCTGGTGCAGTTATTGAGATTCGAGACTGCCAAA<br>AGCCGGTTCCCGGTGTGTATGTCCATAAGGGTGTTGTGCAGGTAGG<br>TGAGGTAACCGTAGGCGCGAAAGCTCACGCCAGTATCGACAGCAG<br>ACGCCGGAAGGCTATAGCGCGAGCCCACTCAGCTACTCACCTTACG<br>CACCAAGCCTTGCGTGACGCGTTAGGCCCTACAGCGGCACAGGCA<br>GGGTCAGAGAACCAACCAGGAAGATTCCGCTTCGACTTCGGTTCC<br>CCTTCTGCGGTACCTACCGCGGTTATGACTGATGTTGAACAAAAGA<br>TTAATGAAGTACTTGCACGTGATTTGGACGTTCACGCCGAGATTATG<br>GGCATCGATGAAGCGAAGAAGCAAGGTGCGATCGCAGAGTTTGGA<br>GAGAAGTATGGAGAACGGGTACGCGTTGTGACAATTGGAGACTTTA<br>GTAAGGAACTTTGCGGTGGAACGCATGTGCATAACACTGCACAACT<br>GGGCCTGGTCAAACTTCTGGGCGAATCTTCAATCGGAAGTGGGGTA<br>CGACGCATTGAGGCACTGGTCGGGGTGGACGCATACACATTCCTTG<br>CCCGCGAGCATACAGTGGTAAACCAACTTACAGAACTGTTAAAGG<br>GACGCCCAGAGGAGCTTCCCGAAAAGGTATCATCAATGTTAGGCAA<br>GCTGAAGGACGCGGAAAAGGAGATTGAAAAGTTTCGGGCTGAGA<br>AAGTACTGCAAGCTGCCGCTGGATTAGCTGAATCGGCCAAGGACGT<br>GCGTGGCGTTGCGGTCGTCACTGGACAAGTACCGGATGGTACTACA<br>CCTGACGATCTGCGTAAACTGGTGCTGGACGTTCGTGGACGCATCC |

| | | |
|---|---|---|
| | | AAGGCGGACGCGCGGCCGTAGTAGCCTTATTCACTGTGAACAACG GTAAGCCTCTTACGGTAATAGCGACTAACGAAGCAGCCCGTGAAAG AGGACTGAAAGCGGGTGACTTAGTTCGCACCGCAGCGAAAACCCT GGGCGGTGGCGGTGGTGGAAAGCCGGACGTCGCACAAGGCGGTG GACAAAACCCTGCTGCTGTGGGTGAAGCGGTAGAAGCAGTTGAGC GTTTGGTCGCGGACACCGCTAAAGGTGGTGGTGGTCACCACCACC ACCACCACTGA |
| | **Protein Sequence** | MESAEIRRRWLSFFEERGHTVVPSASLIADDPTLLLVPAGMVPFKPYFL GEVKPPFPRATSVQKCVRTPDIEEVGKTTRHGTFFQMCGNFSFGDYFK EGAIKLAWELLTTPQDKGGYGLDPERLWITVYKDDDEAERIWHEVVG VPKERIQRLGMKDNYWSMGVPGPCGPCSEINYDRGPEFGVEGGPAVN DERYVEIWNLVFMQYERGEGTGKDNFEILGELPSKNIDTGLGLERLA MILQGVQNMYEIDTSMAVIDKATELTGVRYGDAHDSDVSLRVVTDH MRTATMLIGDGVTPGNEGRGYVLRRIMRRAIRNMRLLGATGPVVKDL LDTVIQMMGRQYPELVTDRERIEKVAIAEENAFLKTLKAGTNILDTAV TETKQAGGTVLSGDKAFLLHDTWGFPIDLTLEMAAEQGLSVDEEGFR RLMKEQRERAKADAQAKKTGHADMGAYREIADTAGETDFIGYSDTE GESTIVGLLVDGISSPAATEGDEVEVVLDRTPFYAEGGGQIGDTGRIKV DSGAVIEIRDCQKPVPGVYVHKGVVQVGEVTVGAKAHASIDSRRRKA IARAHSATHLTHQALRDALGPTAAQAGSENQPGRFRFDFGSPSAVPTA VMTDVEQKINEVLARDLDVHAEIMGIDEAKKQGAIAEFGEKYGERVR VVTIGDFSKELCGGTHVHNTAQLGLVKLLGESSIGSGVRRIEALVGVD AYTFLAREHTVVNQLTELLKGRPEELPEKVSSMLGKLKDAEKEIEKFR AEKVLQAAAGLAESAKDVRGVAVVTGQVPDGTTPDDLRKLVLDVRG RIQGGRAAVVALFTVNNGKPLTVIATNEAARERGLKAGDLVRTAAKTL GGGGGGKPDVAQGGGQNPAAVGEAVEAVERLVADTAKGGGGHHHH HH |

#### Table S3. Primers used in this study.

| ID# | Name | Sequence (5'-3') | Description |
|---|---|---|---|
| prCL070 | Sin BGC-1_LD1-F | GTGGGTACCAGCGGGGCGAG | For construction of *sin* BGC expression, gene deletion and bottom-up expression plasmid in *S. albidoflavus* |
| prCL071 | Sin BGC-2_LD1-R | GATGCAGCTCCTGAGCATGGGTGTCCCTCACGTCGCTTGGG | |
| prCL072 | Sin BGC-3_SinB-F | ATGCTCAGGAGCTGCATCGTTCAAT | |
| prCL073 | Sin BGC-4_SinC-R | ATCGCGTCGCCTCACCTGTC | |
| prCL074 | Sin BGC-5_Sin-PHP-F | GACAGGTGAGGCGACGCGATCCTTCC | |
| prCL085 | Sin BGC-6-2_Sin-PHP-R | GCACATGGTCGCCGAGACCCGG | |
| prCL086 | Sin BGC-14_Sin-LD2-F | CAGCGGGACAGACACCGGGTCTCG | |
| prCL087 | Sin BGC-15_Sin-LD2-R | TATCCGGAGGCGGAACGGAAGGCC | |
| prCL076 | Sin BGC-7_Sin-MFS-F | TTGACGACAAGCCCGGCCTTCCG | |
| prCL077 | Sin BGC-8_SinD-R | ATTCTGGGAAATTCCCGGCGCGTAATGTTAC | |
| prCL078 | Sin BGC-9_His-SinA-F | CCGGGAATTTCCCAGAATATGCATCATCATCATCATCACGCACAGGCG | |
| prCL079 | Sin BGC-10_SinA-R | CTACGCGTCGCCGGCTACCCG | |
| prCL082 | pSYH-SinBGC_Fwd | TAGCCGGCGACGCGTAGATGATCTCGGTACCAAACCAATTATTAAAGACGC | |
| prCL083 | pSYH-SinBGC_Rev | GCTCGCCCCGCTGGTACCCATTAGATGTCTCCTTACTTAGATACACC | |
| prCL110 | SinBGC_xsinB-F | CAGCCACCACCTGCTCATCTCAGCCCGTG | |
| prCL111 | SinBGC_xsinB-R | GAGATGAGCAGGTGGTGGCTGATGTGACGATGAGTC | |
| prCL112 | SinBGC_xsinC-R | ATCGCGTCGCCTCACCTGTCTGGGTCATTTCGTCTCCCCACG | |
| prCL131 | SinBGC_xsinA-F | ACATTACGCGATGATCTCGGTACCAAACCAATTATTAAAGACGC | |
| prCL132 | SinBGC_xsinA-R | CGAGATCATCGCGTAATGTTACCGATTGGCAC | |
| prCL133 | SinBGC_xLD1-R | CGATGCAGCTCCTGAGCATTAGATGTCTCCTTACTTAGATACACC | |
| prCL134 | SinBGC_xLD2-F | TATGACCGCGCTACACCGCCGAACTCCGGTG | |
| prCL135 | SinBGC_xLD2-R | GGTGTAGCGCGGTCATAGTTCCGCCTCG | |
| prCL136 | SinBGC_xPHP-F | CTTCCCTCAAGTCCGGGTCAAGTCGTGGGTG | |
| prCL137 | SinBGC_xPHP-R | TGACCCGGACTTGAGGGAAGGATCGCGTCGCC | |
| prCL138 | SinBGC_xAPH-F | TGACTGAGCCCATGGTCGCAGCCCTTGAAGC | |
| prCL139 | SinBGC_xAPH-R | CGACCATGGGCTCAGTCAGTGTGTCCAGTCGC | |
| prCL144 | SinBGC_xsinD-F | AAGCCTGACATTACGCGCCGGGAATTTCCCA | |
| prCL145 | SinBGC_xsinD-R | CGCGTAATGTCAGGCTTCAAGGGCTGCGACC | |
| prCL155 | SinBGC_xMFS-R | CCGTTCCGCCTCCGGATACAGACGAAGCCACCGGCAAAAAC | |
| prCL176 | SinMFS-F | GAATCCGTGACACCGCCGAACTCCGGTGC | |

| | | | |
|---|---|---|---|
| prCL177 | SinC-R | CGGCGGTGTCACGGATTCCCGGGCGGAGC | |
| prCL178 | pSYH-synP_APH_vec-F | CTACATCAGTGATCTCGGTACCAAACCAATTATTAAAGACGC | |
| prCL179 | pSYH-synP_SinAPH_R | ACCGAGATCACTGATGTAGGTCAGGCTTCAAGGG | |
| prCL183 | sinBC_APH insert-F | AGGTGAGGCGACGCGATTACAGACGAAGCCACCGGCAAAAAC | |
| prCL184 | sinB_MFS-R | GGCGGTGTCACGGATTCCTGGGTCATTTCGTCTCCCCACG | |
| prCL185 | sinB-APH-F | GGAGACGAAATGACCCAGTACAGACGAAGCCACCGGCAA | |
| prCL186 | sinB-R | CTGGGTCATTTCGTCTCCCCACG | |
| prCL028 | pET28b-Sin-arg-F2 | TGCCGCGCGGCAGCCATATGACCCAGCAGCTCCCGGA | For construction of protein expression plasmid in *E. coli* |
| prCL029 | pET28b-Sin-arg-R2 | GTGGTGGTGGTGGTGCTCGAGGAATTCAGCGGTCACCAC | |
| prCL146 | Sin-aph-F | CCGCGCGGCAGCCATATGTCGGAAAGATTAGATACCTTGAC | |
| prCL019 | pET28_Sin_PHP+APH_insert-rev | GGTGGTGGTGCTCGACTATGCCTCTAAAGCGGCTAC | |
| prCL149 | Sin-php-F | CCGCGCGGCAGCCATATGCCTGGAGTCCCAGCCATTTTAG | |
| prCL150 | Sin-php-R | GGTGGTGGTGCTCGATTACAGCTCTGCTTCTGGTTTTAAAG | |
| prHZ001 | pACYC vector-F | GCATAATGCTTAAGTCGAACAGAAAGTAATCGTATTGTACAC | |
| prHZ002 | pACYC vector-R | CTGGCTGTGGTGATGATGGTGATG | |
| prHZ003 | pACYC-SinPEARL1-F | CATCACCATCATCACCACAGCCAGATGGGAACCTCGGGTG | |
| prHZ004 | pACYC- SinPEARL1-R | CGACTTAAGCATTATGCTTAACCGACTCGTCTAGAG | |
| prHZ005 | pACYC- SinPEARL2-F | CATCACCACAGCCAGATGCGACCTACTGCTCACC | |
| prHZ006 | pACYC- SinPEARL2-R | GACTTAAGCATTATGCTTACCGGCGTCTGCCAC | |
| prHZ007 | PET28a vector-F | GGTGGTGGTGGTCACCACCACCACCACCACTGAGATCCG | |
| prHZ008 | PET28a vector-R | GGTATATCTCCTTCTTAAAGTTAAACAAAATTATTTCTAGAGGGGAATTGTTATCCGC | |
| prHZ009 | pET28a-siAlaRS-F | CTAGAAATAATTTTGTTTAACTTTAAGAAGGAGATATACCATGGAAAGTGCGGAAATTC | |
| prHZ010 | pET28a-siAlaRS-R | CCGGATCTCAGTGGTGGTGGTGGTGGTGACCACCACCACCTTTAGCGGTGTCCG | |
| prCL032 | pBAD vector-F | TATGGGAATTCGAAGCTTGGG | For complementation experiment in *S. enterica* |
| prCL033 | pBAD vector with N-His-R | CATATGGCTGCCGCGCG | |
| prCL034 | pBAD-Sin-adetran-F | CGCGCGGCAGCCATATGGTTAACTTGACCCGCATCTAC | |
| prCL035 | pBAD-Sin-adetran-R | CCAAGCTTCGAATTCCCATATTAACGATTCGCCCCGGG | |
| prCL189 | Ecoli_sinD-R137A-F | AGCACGTGCTGAACGCTCGACATG | |
| prCL190 | Ecoli_sinD-R137A-R | AGCGTTCAGCACGTGCTACTACGGTGCGAGCAAGATGC | |

#### Table S4. DNA primer sequences for synthesizing tRNA$^{Ala}$s.

| Name | Sequence (5'-3') |
|---|---|
| SiAla1-F | AATTCCTGCAGTAATACGACTCACTATAGGGGTCGTAGCACAGAGGTAGTGCGCCGC |
| SiAla1-R | mUmGGGGGCGGGGCGGTCGGATTTGAACCGACGTCCTCCGAGATGCTGTCGCGGCGCACT |
| SiAla2-F | AATTCCTGCAGTAATACGACTCACTATAGGGGCCTTAGCTCAGTTGGTAGAGCGCCG |
| SiAla2-R | mUmGGTGGAGCCTAGGGGAGTCGAACCCCTGACATCCGCCATGCAAAGACGGCGCTCTA |
| SiAla3-F | AATTCCTGCAGTAATACGACTCACTATAGGGCCTGTGGCGCAGTCCGGTAGCGCACC |
| SiAla3-R | mUmGGTGGACCTGAGGGGATTTGAACCCCTGGCCCCCTCGATGCGAACGAGGTGCGCTAC |
| SiAla4-F | AATTCCTGCAGTAATACGACTCACTATAGGGGCTATAGCTCAGTTGGTAGAGCGCCT |
| SiAla4-R | mUmGGTGGAGCTAAGGAGAATTGAACTCCTGACCTCCTGCATGCCATGCAGGCGCTCTA |

#### Table S5. List of SinC homolog accession IDs used to generate the consensus sequence.

| | | | | | | | |
|---|---|---|---|---|---|---|---|
| **1** | WP_259622325.1 | **26** | WP_403071258.1 | **51** | WP_403942556.1 | **76** | SinC |
| **2** | WP_204029241.1 | **27** | WP_401055025.1 | **52** | WP_362780823.1 | **77** | AKJ15732.1 |
| **3** | WP_271216415.1 | **28** | WP_359909551.1 | **53** | WP_317886067.1 | **78** | WP_405606862.1 |
| **4** | WP_363943350.1 | **29** | WP_380548035.1 | **54** | WP_205360540.1 | **79** | WP_391827195.1 |
| **5** | HZM76203.1 | **30** | WP_387693625.1 | **55** | WP_360561391.1 | **80** | WP_382036519.1 |
| **6** | WP_179084206.1 | **31** | WP_355254138.1 | **56** | WP_358974552.1 | **81** | WP_190021319.1 |
| **7** | WP_150242401.1 | **32** | WP_387758603.1 | **57** | WP_365657760.1 | **82** | WP_168093386.1 |
| **8** | KAA6223796.1 | **33** | BFD94196.1 | **58** | WP_159050442.1 | **83** | WP_394617058.1 |
| **9** | WP_347667940.1 | **34** | WP_401117360.1 | **59** | KUM92580.1 | **84** | WP_072625732.1 |
| **10** | WP_393424040.1 | **35** | WP_380661352.1 | **60** | WP_384258535.1 | **85** | AVT38634.1 |
| **11** | WP_229402282.1 | **36** | WP_380743071.1 | **61** | WP_301490458.1 | **86** | WP_159104707.1 |
| **12** | WP_281554895.1 | **37** | WP_388044806.1 | **62** | WP_159702671.1 | **87** | WP_107270352.1 |
| **13** | WP_406586939.1 | **38** | WP_049649751.1 | **63** | WP_400473697.1 | **88** | WP_334437929.1 |
| **14** | WP_398741471.1 | **39** | WP_074004065.1 | **64** | WP_380486114.1 | **89** | WP_091609756.1 |
| **15** | WP_125498608.1 | **40** | WP_212008091.1 | **65** | WP_063346604.1 | **90** | WP_088970654.1 |
| **16** | WP_168131207.1 | **41** | WP_212238895.1 | **66** | WP_380339762.1 | **91** | HEY0639865.1 |
| **17** | RBM19018.1 | **42** | WP_194922991.1 | **67** | WP_380424451.1 | **92** | HSZ31459.1 |
| **18** | WP_147255521.1 | **43** | WP_194891496.1 | **68** | WP_380492437.1 | **93** | HEX2051111.1 |
| **19** | WP_402400803.1 | **44** | WP_402001799.1 | **69** | WP_380559703.1 | **94** | WP_095984442.1 |
| **20** | WP_360354533.1 | **45** | WP_406727214.1 | **70** | WP_363618373.1 | **95** | WP_081713813.1 |
| **21** | WP_380455311.1 | **46** | WP_386875550.1 | **71** | WP_387982065.1 | **96** | EPX60360.1 |
| **22** | WP_396299053.1 | **47** | GHD86847.1 | **72** | WP_380370652.1 | **97** | HZJ65492.1 |
| **23** | WP_380765853.1 | **48** | WP_190177065.1 | **73** | WP_402437597.1 | **98** | WP_428268523.1 |
| **24** | WP_224275093.1 | **49** | WP_202235228.1 | **74** | WP_363441976.1 | **99** | HWO26310.1 |
| **25** | WP_363386324.1 | **50** | WP_403663021.1 | **75** | WP_037862919.1 | | |

**Table S6. $^{1}H$ NMR (500 MHz) and $^{13}C$ NMR (125 MHz) spectroscopic data of sinefungin (1) in $D_2O$.**

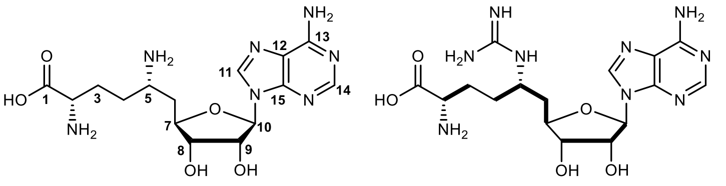

sinefungin (**1**)

— COSY

| | | From reference;[9] in $D_2O$ | | From this work; in $D_2O$ | | |
|---|---|---|---|---|---|---|
| position | | δH (mult, *J* in Hz) | δC | δH (mult, *J* in Hz) | δC | COSY |
| | 1 | | 173.8 | | 173.8 | |
| | 2 | 3.62 (1H, t, 7.0) | 53.8 | 3.79 (1H, m) | 54.1 | 1.95 |
| | 3 | 1.80 (2H, m) | 22.8 | 1.95 (2H, m) | 26.4 | 3.79 |
| | 4 | 1.70 (1H, m) | 26.5 | 1.81 (1H, m) | 27.6 | 3.58 |
| | | 1.90 (1H, m) | | 1.92 (1H, m) | | |
| | 5 | 3.41 (1H, m) | 48.8 | 3.58 (1H, m) | 48.8 | 1.81, 1.92, 2.22 |
| | 6 | 2.06 (2H, m) | 34 | 2.22 (2H, t, 6.1) | 34.3 | 3.58, 4.28 |
| | 7 | 4.15 (1H, dt, 6.2, 6.0) | 72.5 | 4.28 (1H, q, 6.2) | 73.0 (overlapped with C9) | 2.22 |
| adenosine | 8 | 4.20 (1H, dd, 6.2, 6.2) | 79.5 | 4.34 (1H, t, 5.7) | 79.5 | 4.76 |
| | 9 | 4.60 (1H, dd, 7.0, 6.2) | 72.3 | 4.76 (1H, m, overlapped with solvent peak) | 73.0 (overlapped with C7) | 4.34, 6.04 |
| | 10 | 5.88 (1H, d, 7.0) | 88.5 | 6.04 (1H, d, 4.4) | 88.7 | 4.76 |
| | 11 | 8.10 (1H, s) | 140.5 | 8.28 (1H, s) | 140.8 | |
| | 12 | | 119.2 | | 119.0 | |
| | 13 | | 155.8 | | 154.6 | |
| | 14 | 8.08 (1H, s) | 153 | 8.25 (1H, s) | 151.4 | |
| | 15 | | 148.5 | | 148.5 | |

**Table S7. $^{1}H$ NMR (600 MHz) and $^{13}C$ NMR (150 MHz) spectroscopic data of sinefungin lactam (2) in methanol-*d4*.**

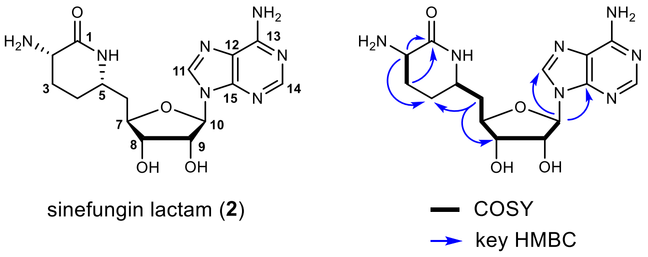

| position | | δH (mult, *J* in Hz) | δC (type) | COSY | HMBC | NOESY |
|---|---|---|---|---|---|---|
| | **1** | | 169.3 (C) | | | |
| | **2** | 3.82 (1H, m) | 50.5 (CH) | 1.93, 2.12 | 22.9, 26.5, 169.3 | 2.12 |
| | **3** | 1.93 (1H, overlapped) | 22.9 (CH2) | 3.82 | 50.5, 169.3 (w) | 2.12 |
| | | 2.12 (1H, overlapped) | | | | 1.94 |
| | **4** | 2.05 (1H, overlapped) | 26.5 (CH2) | | | 1.93, 3.70 |
| | | 1.94 (1H, overlapped) | | | | 3.70, 4.13 (w) |
| | **5** | 3.70 (1H, m) | 50.0 (CH) | 2.06 | | 1.94, 2.05, 4.14 |
| | **6** | 2.06 (2H, overlapped) | 40.4 (CH2) | 3.70, 4.13 | 26.5, 50.0, 75.2, 82.4 | 4.13, 4.25 |
| | **7** | 4.13 (1H, m) | 82.4 (CH) | 2.06 | 50.0 | 1.94, 2.06, 3.70, 5.94 (w) |
| **adenosine** | **8** | 4.25 (1H, t, 5.5) | 75.2 (CH) | 4.13, 4.77 | 40.4, 82.4 (w), 90.9 | 2.06, 3.70 (w), 4.77, 8.24 (w) |
| | **9** | 4.77 (1H, t, 5.0) | 74.6 (CH) | 4.25, 5.94 | 82.4, 90.9 | 2.06, 4.25, 5.94 |
| | **10** | 5.94 (1H, d, 4.5) | 90.9 (CH) | 4.77 | 74.6, 141.7, 150.5 | 4.13, 4.77, 8.24 |
| | **11** | 8.24 (1H, s) | 141.7 (CH) | | 120.8, 150.5 | 4.25 (w), 4.77, 5.94 |
| | **12** | | 120.8 (C) | | | |
| | **13** | | 157.5 (C) | | | |
| | **14** | 8.21 (1H, s) | 153.9 (CH) | | 120.8 (w), 150.5, 157.5 | |
| | **15** | | 150.5 (C) | | | |

(w) = weak signal.

**Table S8. $^{1}H$ NMR (600 MHz) and $^{13}C$ NMR (150 MHz) spectroscopic data of guanidinyl-sinefungin (4) in methanol-*d4*.**

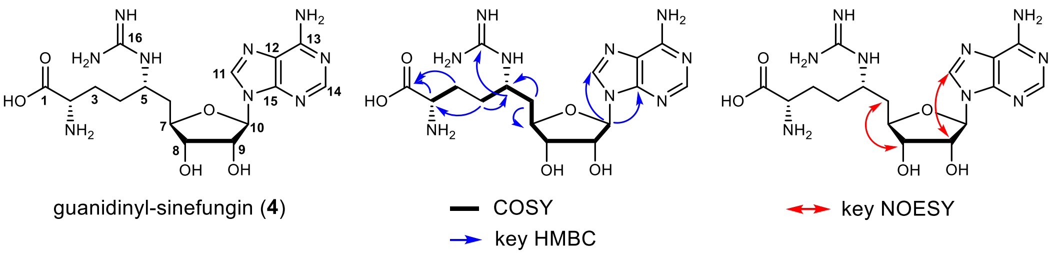

| position | | δH (mult, *J* in Hz) | δC (type) | COSY | HMBC | NOESY |
|---|---|---|---|---|---|---|
| | 1 | | 174.3 (C) | | | |
| | 2 | 3.57 (1H, m) | 55.6 (CH) | 1.88 | 28.5, 174.3 | 1.88 |
| | 3 | 1.88 (2H, overlapped) | 28.5 ($CH_2$) | | 32.3, 55.6, 174.3 | |
| | 4 | 1.64 (1H, m) | 32.3 ($CH_2$) | 1.76 | 28.5, 50.2, 55.6 | |
| | | 1.76 (1H, m) | | 1.64 | 50.2, 55.6 | 3.70 |
| | 5 | 3.70 (1H, m) | 50.2 (CH) | 1.76, 1.90 | 158.7 (w) | 1.76, 1.88 (w) |
| | 6 | 1.90 (2H, overlapped) | 39.3 ($CH_2$) | 2.17, 3.68 | 50.2 | 2.17 |
| | | 2.17 (H, ddd, 14.4, 11.1, 3.2) | | 1.90, 4.10 | 82.0 (w) | 1.90, 3.70, 4.10, 4.24 |
| | 7 | 4.10 (1H, ddd, 11.2, 4.7, 2.5) | 82.0 (CH) | 2.17, 4.24 | 50.2 (w) | 1.90, 2.17, 4.24 |
| adenosine | 8 | 4.24 (1H, t, 5.0) | 75.2 (CH) | 4.10, 4.87 | 39.3, 90.6 | 2.17, 4.10, 4.87 |
| | 9 | 4.87 (1H, t, 5.1) | 74.5 (CH) | 4.24, 5.97 | 82.0, 90.6 | 4.24, 5.97, 8.26 (w) |
| | 10 | 5.97 (1H, d, 5.0) | 90.6 (CH) | 4.87 | 74.5, 82.0 (w), 141.9, 150.6 | 4.87, 8.26 (w) |
| | 11 | 8.26 (1H, s) | 141.9 (CH) | | 120.8, 150.6, 157.4 | 4.87, 5.97 |
| | 12 | | 120.8 (C) | | | |
| | 13 | | 157.4 (C) | | | |
| | 14 | 8.22 (1H, s) | 154.0 (CH) | | 120.8, 150.6, 157.4 | |
| | 15 | | 150.6 (C) | | | |
| guanidine | 16 | | 158.7 (C) | | | |

(w) = weak signal.

**Table S9. $^{1}H$ NMR (600 MHz) and $^{13}C$ NMR (150 MHz) spectroscopic data of guanidinyl-sinefungin lactam (5) in methanol-$d_4$.**

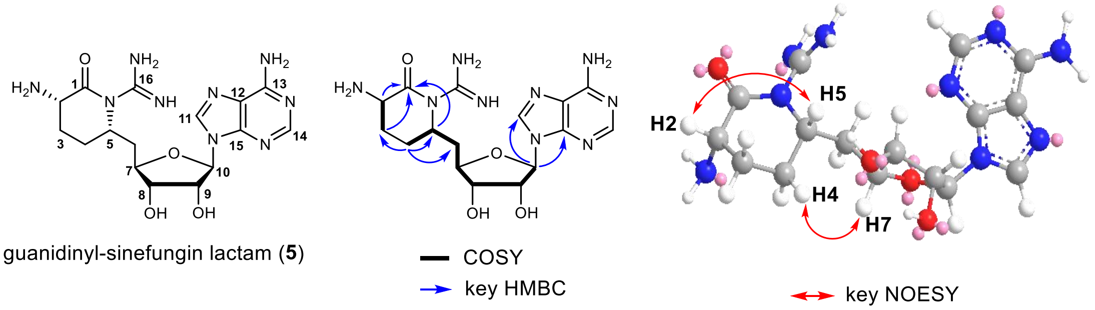

| position | | δH (mult, *J* in Hz) | δC (type) | COSY | HMBC | NOESY |
|---|---|---|---|---|---|---|
| | 1 | | 172.9 (C) | | | |
| | 2 | 4.25 (1H, overlapped) | 50.6 (CH) | 1.91 | 25.4, 172.9 | |
| | 3 | 1.91 (2H, m) | 25.4 (CH2) | 4.25 | 27.0, 50.3, 172.9 | 3.65, 4.25 |
| | 4 | 1.80 (1H, m) | 27.0 (CH2) | 3.65 | 25.4, 40.4, 50.3 | 2.00, 3.65, 4.14 |
| | | 2.00 (1H, m) | | 3.65 | 25.4, 40.4, 50.3 | 1.80, 3.65, 4.25 |
| | 5 | 3.65 (1H, m) | 50.3 (CH) | 1.80, 2.11 | 40.4, 82.7, 172.9 | 1.80, 2.05, 2.11, 4.14, 4.25 |
| | 6 | 2.05 (1H, m) | 40.4 (CH2) | 3.65, 4.14 | 27.0, 50.3, 75.2, 82.7 | 4.24 |
| | | 2.11 (1H, m) | | 3.65, 4.14 | 27.0, 50.3, 75.2, 82.7 | 1.80, 1.91, 3.65, 4.14, 4.24 |
| | 7 | 4.14 (1H, m) | 82.7 (CH) | 2.05, 4.24 | 75.2 | 1.80, 2.05, 2.11, 3.65 |
| adenosine | 8 | 4.24 (1H, overlapped) | 75.2 (CH) | 4.14, 4.76 | 40.4, 90.7 | 2.00, 2.05, 4.76 |
| | 9 | 4.76 (1H, t, 5.0) | 74.7 (CH) | 4.24, 5.95 | 82.7, 90.7 | 2.05, 4.24, 5.95, 8.24 |
| | 10 | 5.95 (1H, d, 4.5) | 90.7 (CH) | 4.76 | 74.7, 141.6, 150.5 | 4.14, 4.76, 8.24 |
| | 11 | 8.24 (1H, s) | 141.6 (CH) | | 120.9, 150.5 | 4.24, 4.76, 5.95 |
| | 12 | | 120.9 (C) | | | |
| | 13 | | 157.3 (C) | | | |
| | 14 | 8.20 (1H, s) | 153.9 (CH) | | 150.5, 157.3 | |
| | 15 | | 150.5 (C) | | | |
| guanidine | 16 | | nd | | | |

nd = not detected.

**Table S10. $^{1}$H NMR (600 MHz) and $^{13}$C NMR (150 MHz) spectroscopic data of sinefungin AA (7) in $D_2O$.**

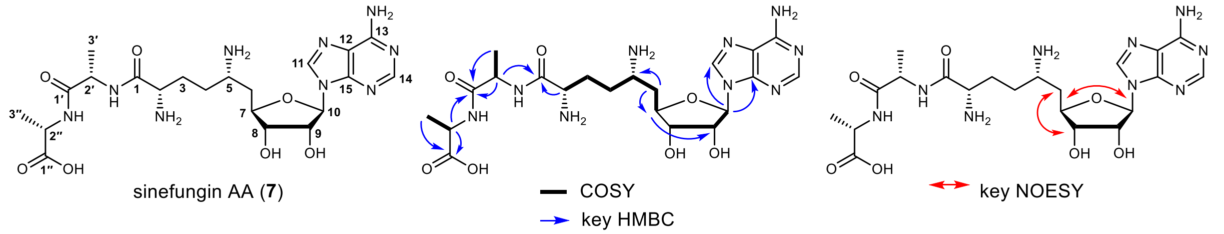

| position | | δH (mult, *J* in Hz) | δC (type) | COSY | HMBC | NOESY |
|---|---|---|---|---|---|---|
| | **1** | | 168.7 (C) | | | |
| | **2** | 4.00 (1H, overlapped) | 51.6 (CH) | 1.94 | 26.6, 168.7 | 1.94 |
| | **3** | 1.94 (2H, m) | 26.6 ($CH_2$, overlapped with C4) | 1.75, 4.00 | | 4.00 |
| | **4** | 1.75 (1H, m) | 26.8 ($CH_2$, overlapped with C3) | 1.83, 1.94, 3.51 | | |
| | | 1.83 (1H, m) | | 1.75, 1.94, 3.51 | | |
| | **5** | 3.51 (1H, m) | 48.6 (CH) | 1.83, 2.15 | | 2.15 |
| | **6** | 2.15 (2H, m) | 34.1 ($CH_2$) | 3.51, 4.20 | 48.6, 79.4 | 3.51, 4.20, 4.28 |
| | **7** | 4.20 (1H, q, 5.7) | 79.4 (CH) | 2.15 | 72.7 | 2.15, 5.98 |
| **adenosine** | **8** | 4.28 (1H, overlapped) | 72.9 (CH) | | 34.1, 88.6 | 2.15, 5.98 |
| | **9** | 4.73 (1H, overlapped with solvent peak) | 72.7 (CH) | 4.28, 5.98 | 79.4 | |
| | **10** | 5.98 (1H, d, 4.5) | 88.6 (CH) | 4.73 | 72.7, 140.5, 148.7 | 4.20, 8.21 |
| | **11** | 8.21 (1H, s) | 140.5 (CH) | | 119.1, 148.7 | 5.98 |
| | **12** | | 119.1 (C) | | | |
| | **13** | | 155.5 (C) | | | |
| | **14** | 8.20 (1H, s) | 152.6 (CH) | | 148.7, 155.5 | |
| | **15** | | 148.7 (C) | | | |
| **alanine** | **1′** | | 173.3 (C) | | | |
| | **2′** | 4.29 (1H, overlapped) | 49.8 (CH) | 1.36 | 16.4, 168.7, 173.3 | 1.36 |
| | **3′** | 1.36 (3H, d, 7.3) | 16.4 ($CH_3$) | 4.29 | 49.8, 173.3 | 4.29 |
| **alanine** | **1″** | | 179.7 (C) | | | |
| | **2″** | 4.03 (1H, overlapped) | 51.0 (CH) | 1.26 | 17.1, 173.3 (w), 179.7 | 1.26 |
| | **3″** | 1.26 (3H, d, 7.3) | 17.1 ($CH_3$) | 4.03 | 51.0, 179.7 | 4.03 |

(w) = weak signal.

**Table S11. $^{1}$H NMR (600 MHz) and $^{13}$C NMR (150 MHz) spectroscopic data of 2’-phosphoguanidinyl-sinefungin (8) in $D_2O$.**

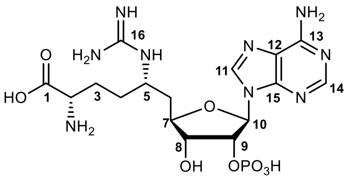

2’-phosphoguanidinyl-sinefungin (**8**)

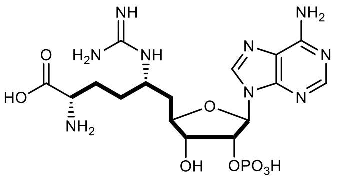

— COSY

| position | | δH (mult, *J* in Hz) | δC (type) | COSY |
|---|---|---|---|---|
| | **1** | | 174.1 (C) | |
| | **2** | 3.64 (1H, t, 5.9) | 54.2 (CH) | 1.79 |
| | **3** | 1.79 (2H, m) | 26.6 (CH2) | 1.48, 1.67, 3.64 |
| | **4** | 1.48 (1H, m) | 29.9 (CH2) | 1.67, 1.79, 3.56 |
| | | 1.67 (1H, m) | | 1.67, 1.79, 3.56 |
| | **5** | 3.56 (1H, m) | 48.8 (CH) | 1.48, 1.67, 1.90 |
| | **6** | 1.90 (1H, m) | 36.9 (CH2) | 2.06, 3.56 |
| | | 2.06 (1H, m) | | 1.90, 4.13 |
| | **7** | 4.13 (1H, m) | 80.5 (CH) | 1.90, 2.06, 4.33 |
| **adenosine** | **8** | 4.33 (1H, t, 5.0) | 73.0 (CH) | 4.13, 5.14 |
| | **9** | 5.14 (1H, m) | 75.4 (CH) | 4.33, 6.05 |
| | **10** | 6.05 (1H, d, 5.1) | 87.2 (CH) | 5.14 |
| | **11** | 8.21 (1H, s) | 140.8 (CH) | |
| | **12** | | 118.9 (C) | |
| | **13** | | 156.5 (C) | |
| | **14** | 8.13 (1H, s) | 152.2 (CH) | |
| | **15** | | 148.8 (C) | |
| **guanidine** | **16** | | 155.2 (C) | |

**Table S12. $^1$H NMR (600 MHz) and $^{13}$C NMR (150 MHz) spectroscopic data of 2'-phosphosinefungin (9) in $D_2O$.**

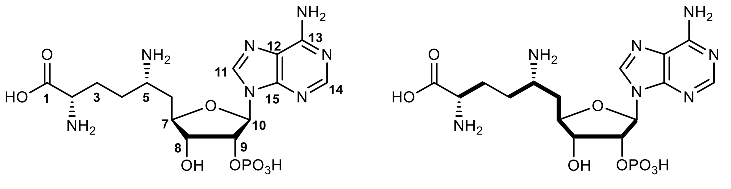

2'-phosphosinefungin (**9**)

| position | | δH (mult, *J* in Hz) | δC (type) | COSY |
|---|---|---|---|---|
| | **1** | | 173.8 (C) | |
| | **2** | 3.70 (1H, m) | 53.9 (CH) | 1.88 |
| | **3** | 1.88 (2H, m) | 26.3 (CH2) | 3.70 |
| | **4** | 1.69 (1H, m) | 27.5 (CH2) | 1.82 |
| | | 1.82 (1H, m) | | 1.69 |
| | **5** | 3.50 (1H, m) | 48.5 (CH) | 1.69 (w), 1.82 (w), 2.15 |
| | **6** | 2.15 (2H, m) | 33.8 (CH2) | 3.50, 4.22 |
| | **7** | 4.22 (1H, m) | 79.3 (CH) | 2.15, 4.40 |
| **adenosine** | **8** | 4.40 (1H, t, 5.9) | 72.4 (CH) | 4.22, 5.03 |
| | **9** | 5.03 (1H m) | 75.3 (CH) | 6.09 |
| | **10** | 6.09 (1H, d, 4.3) | 87.9 (CH) | 5.03 |
| | **11** | 8.20 (1H, s) | 141.1 (CH) | |
| | **12** | | 118.7 (C) | |
| | **13** | | 155.1 (C) | |
| | **14** | 8.17 (1H, s) | 152.0 (CH) | |
| | **15** | | 148.4 (C) | |

(w) = weak signal.

#### Supplementary Schemes

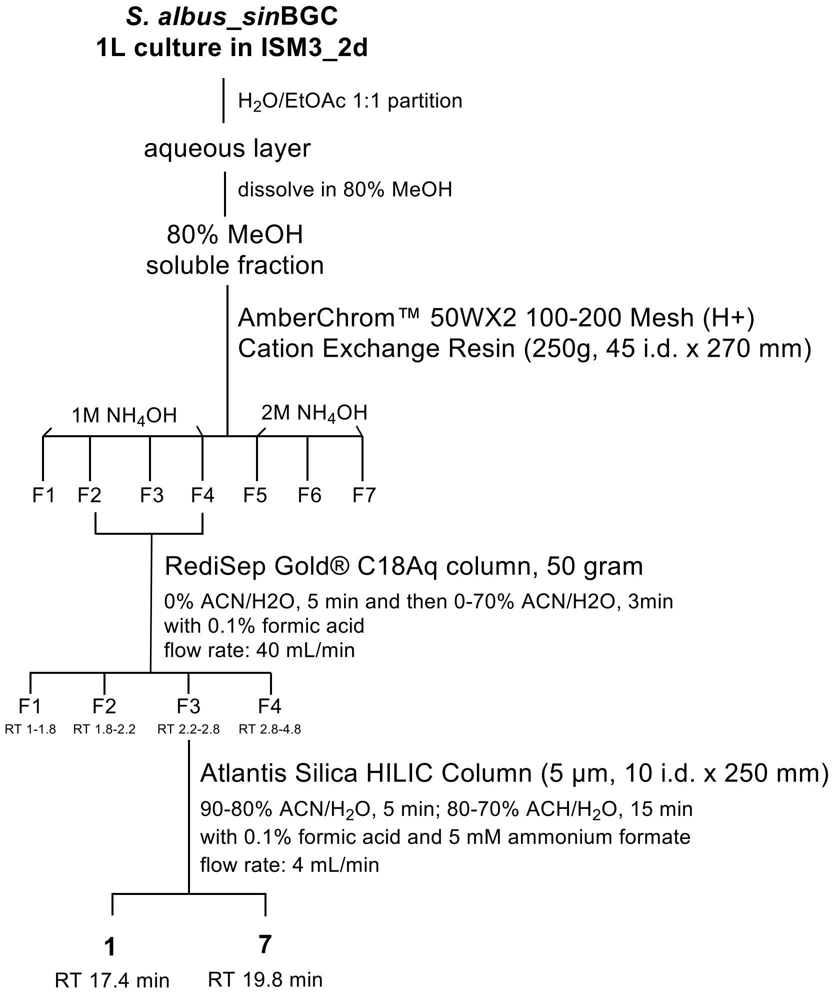

**Scheme S1. Purification procedure for sinefungin (1) and sinefungin AA (7).**

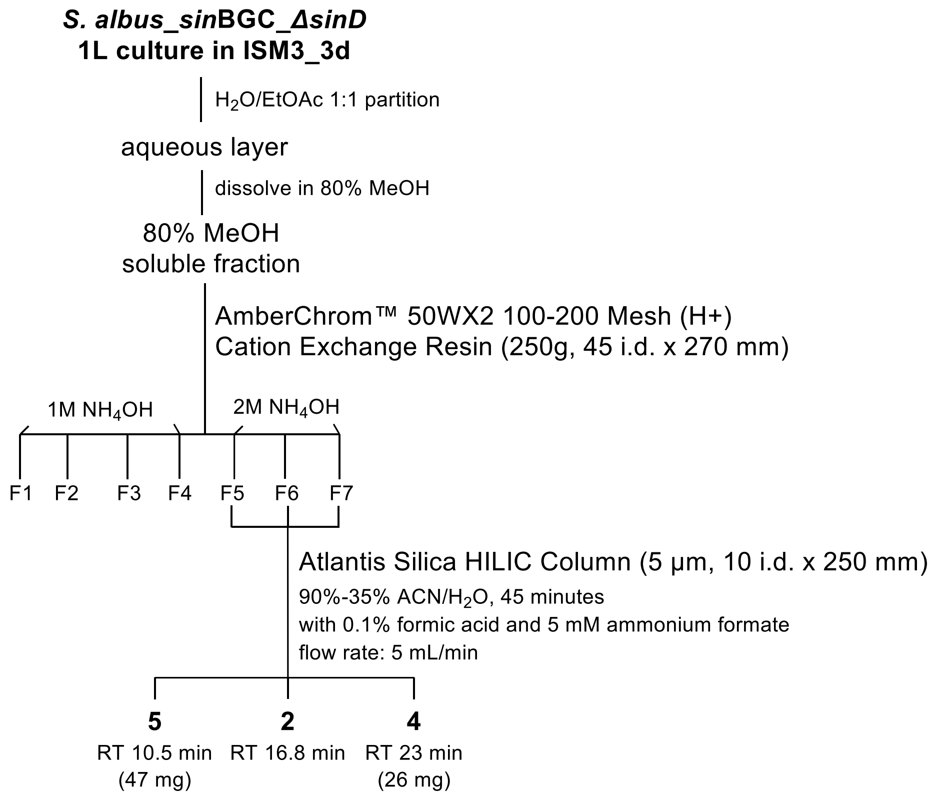

**Scheme S2. Purification procedure for sinefungin lactam (2), guanidinyl-sinefungin (4) and guanidinyl-sinefungin lactam (5).**

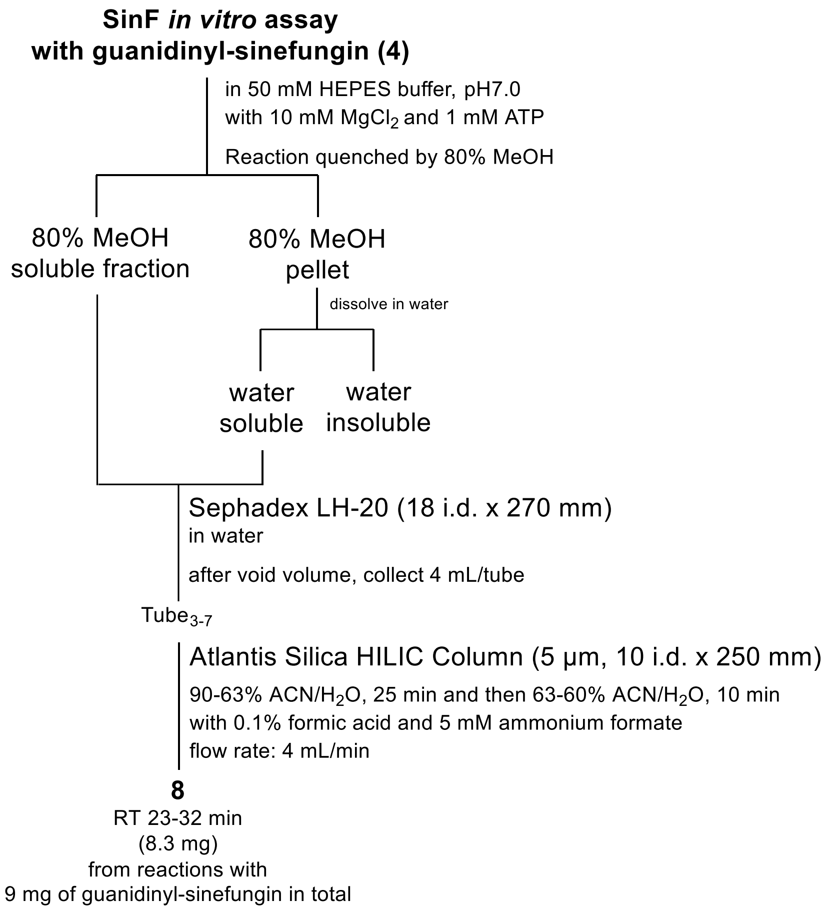

**Scheme S3. Purification procedure for 2'-phosphoguanidinyl-sinefungin (8).**

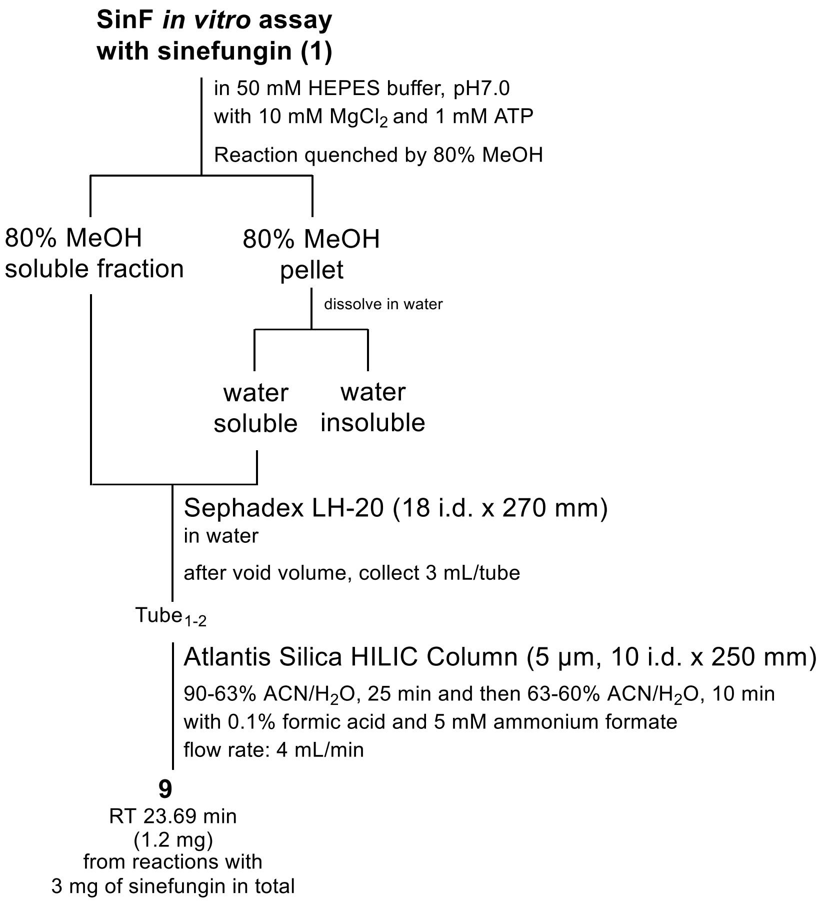

**Scheme S4. Purification procedure for 2'-phosphosinefungin (9).**

### Characterization Data for Purified Compounds

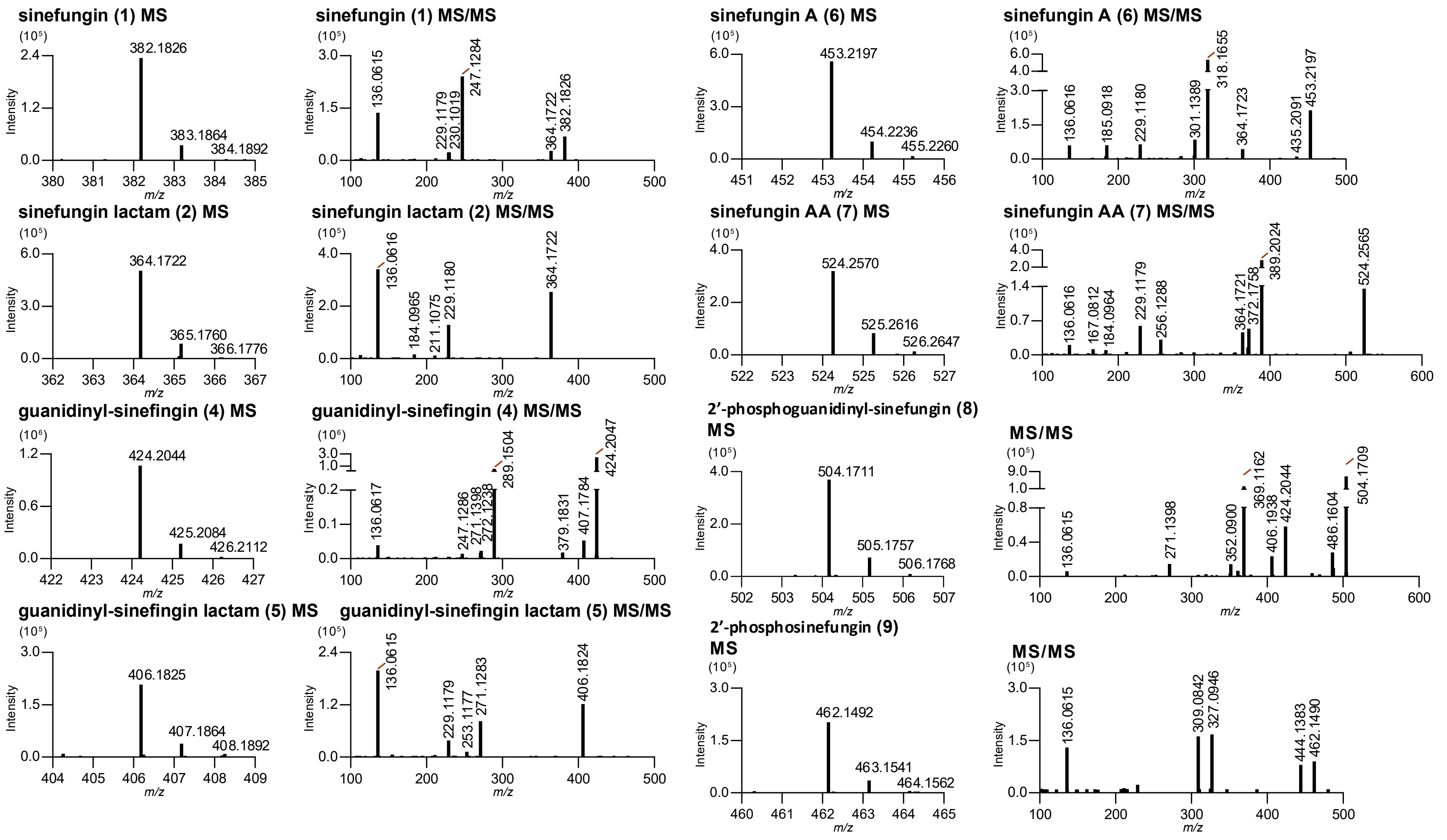

**Figure S10. High-resolution mass spectrometry (HRMS) and MS/MS fragmentation spectra for purified compounds.**

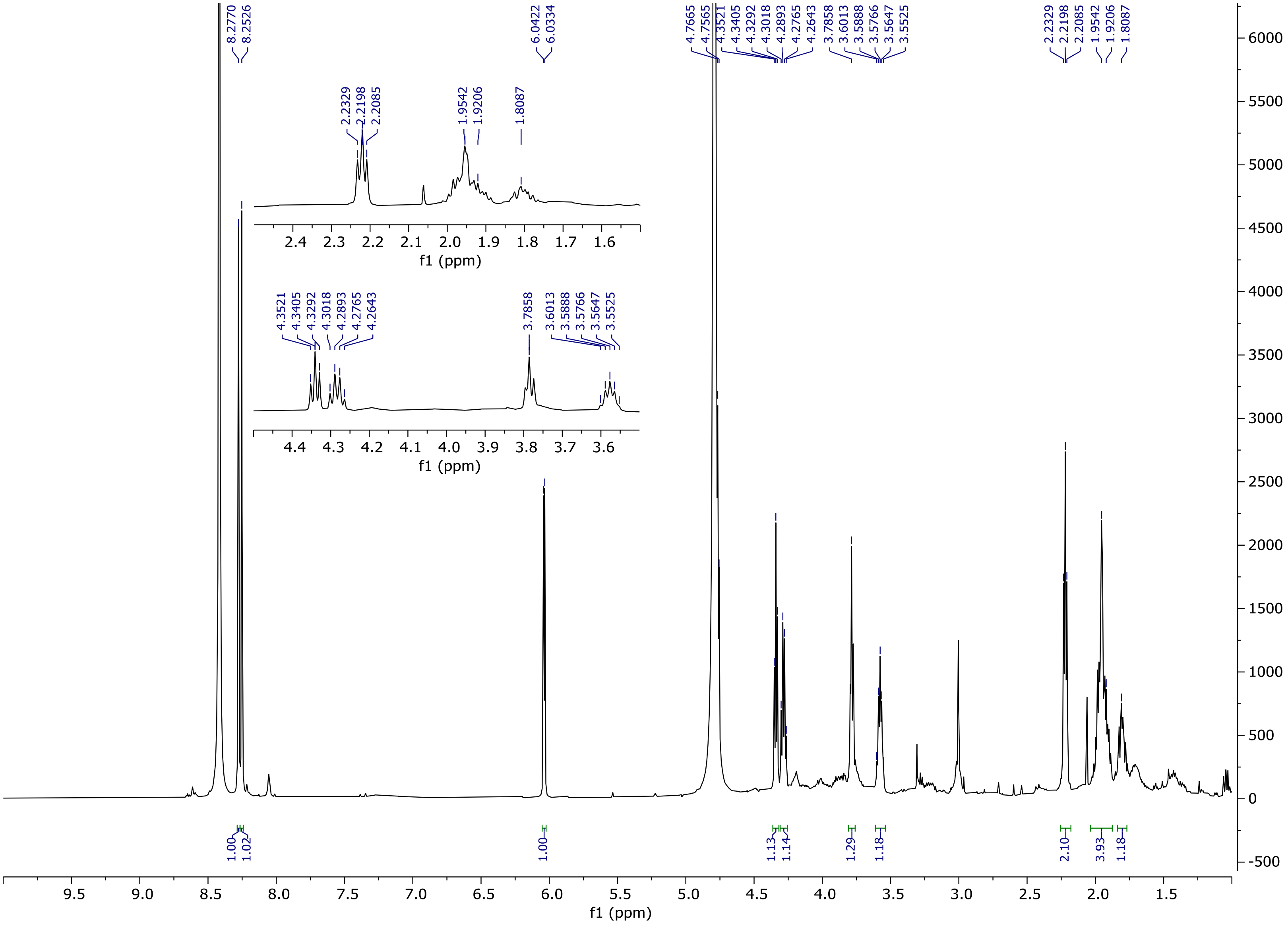

**Figure S11. $^{1}$H NMR spectrum of sinefungin (1) ($D_2O$, 500 MHz).**

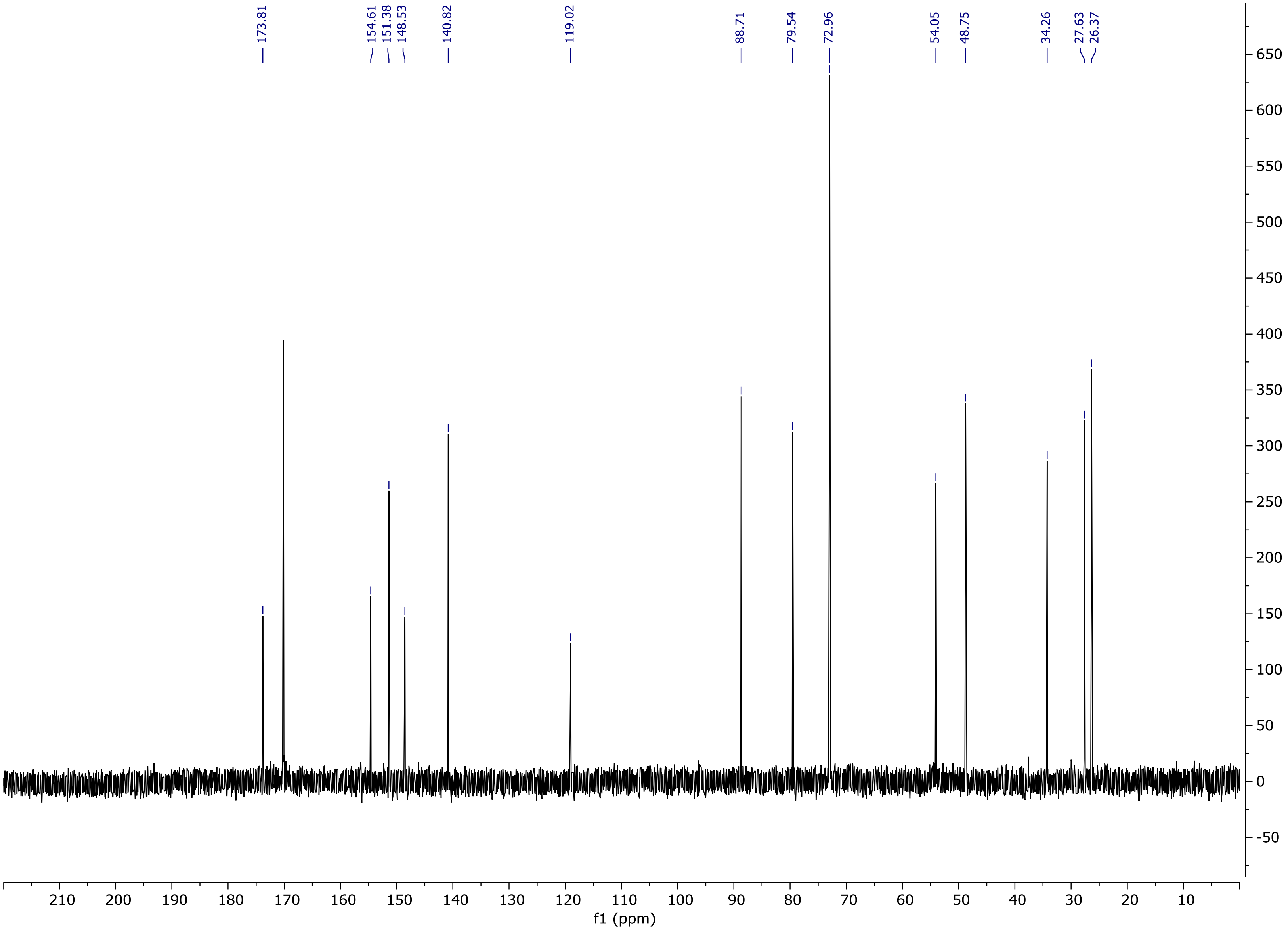

**Figure S12.** $^{13}$**C NMR spectrum of sinefungin (1) (**$D_2O$**, 125 MHz).**

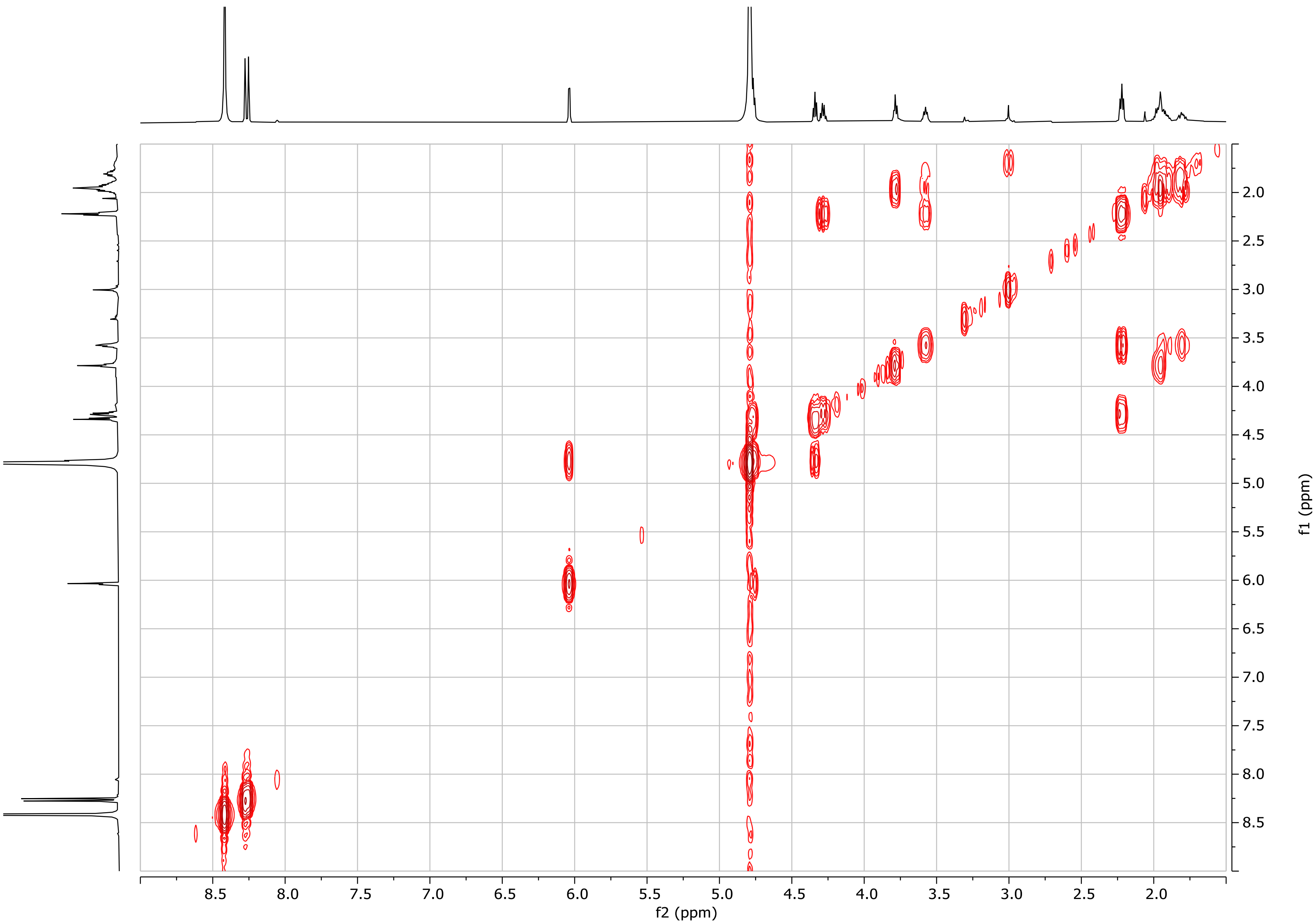

**Figure S13. COSY spectrum of sinefungin (1) ($D_2O$).**

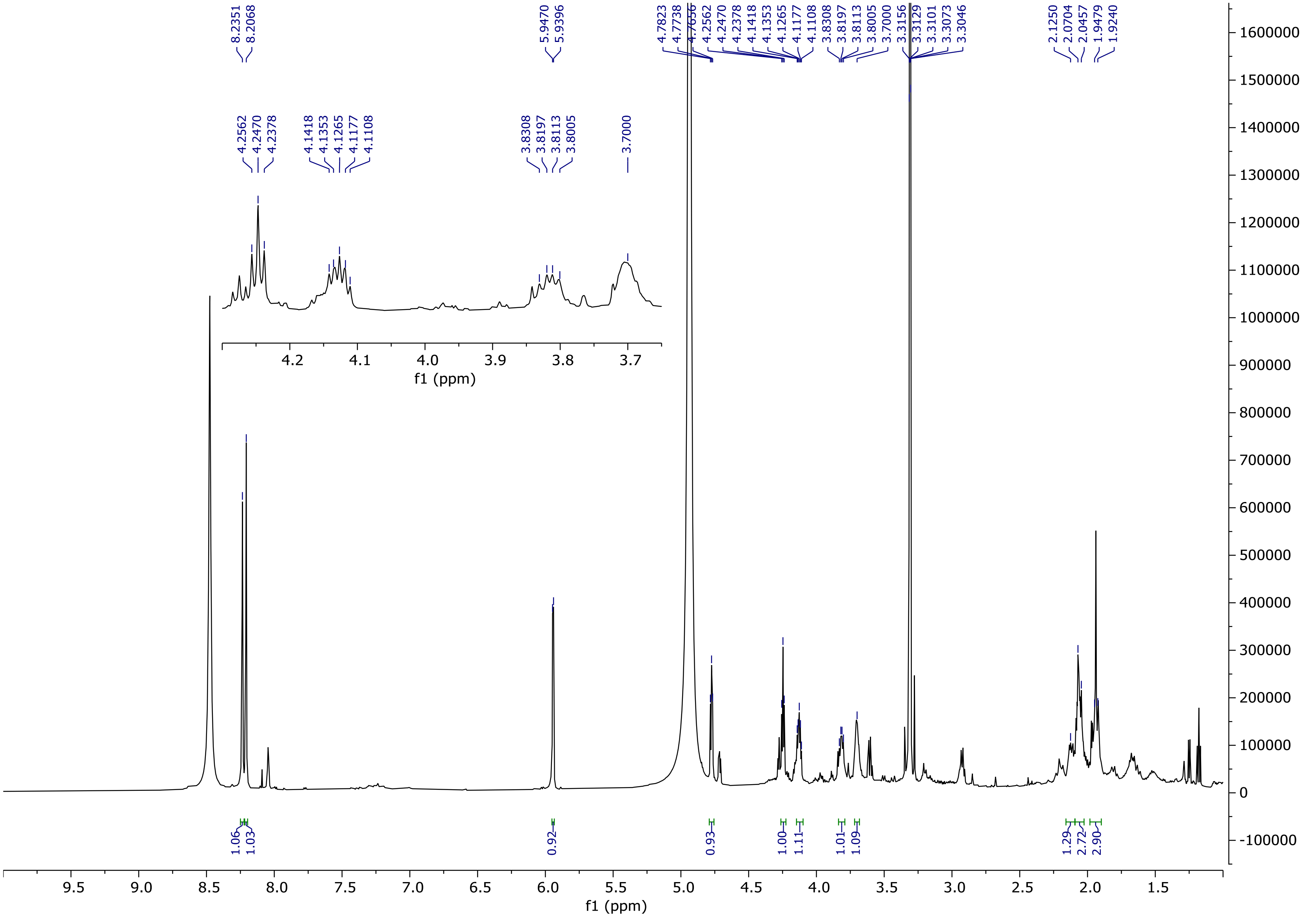

**Figure S14. $^1$H NMR spectrum of sinefungin lactam (2) (methanol-$d_4$, 600 MHz).**

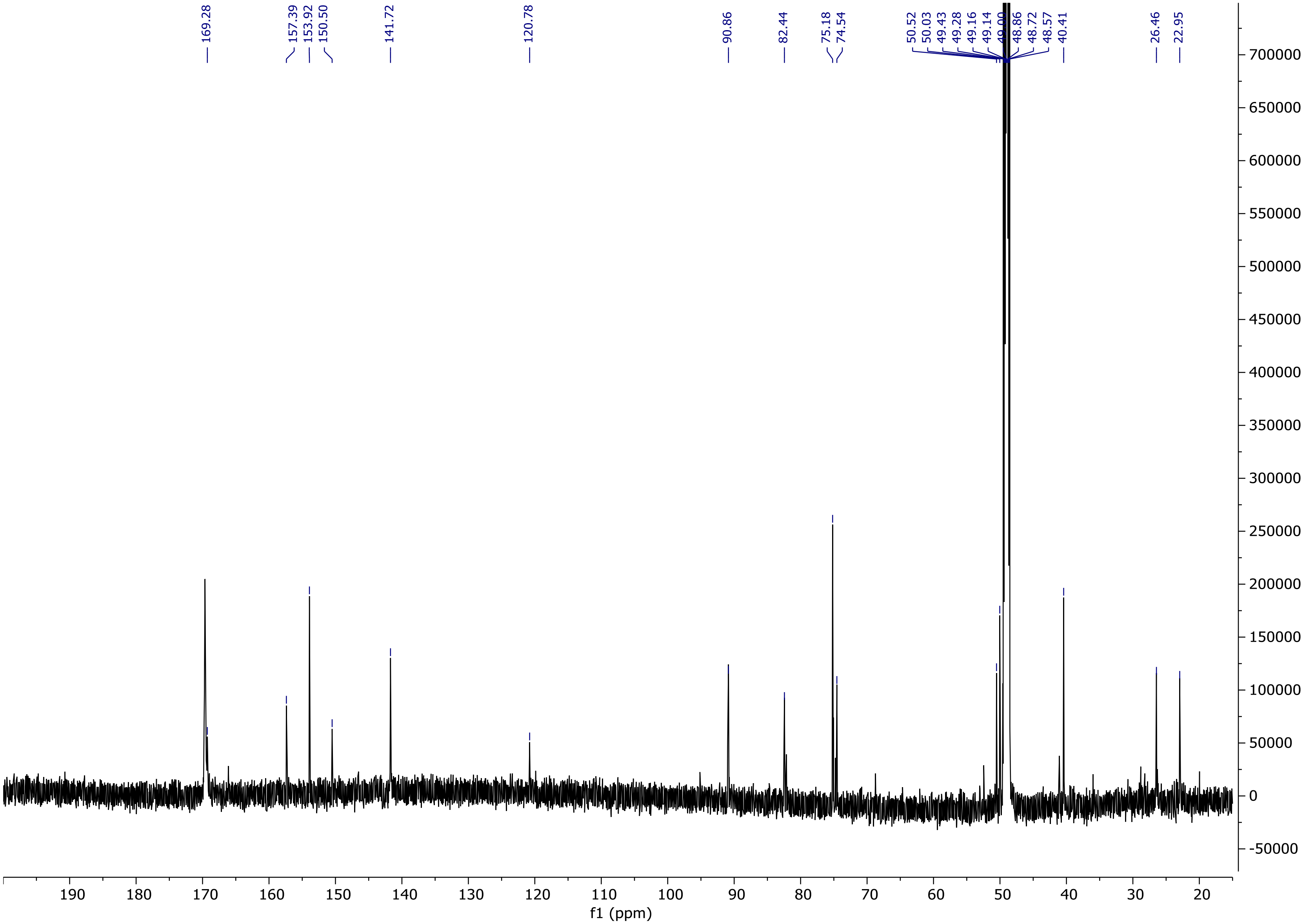

**Figure S15. $^{13}$C NMR spectrum of sinefungin lactam (2) (methanol-*d₄*, 150 MHz).**

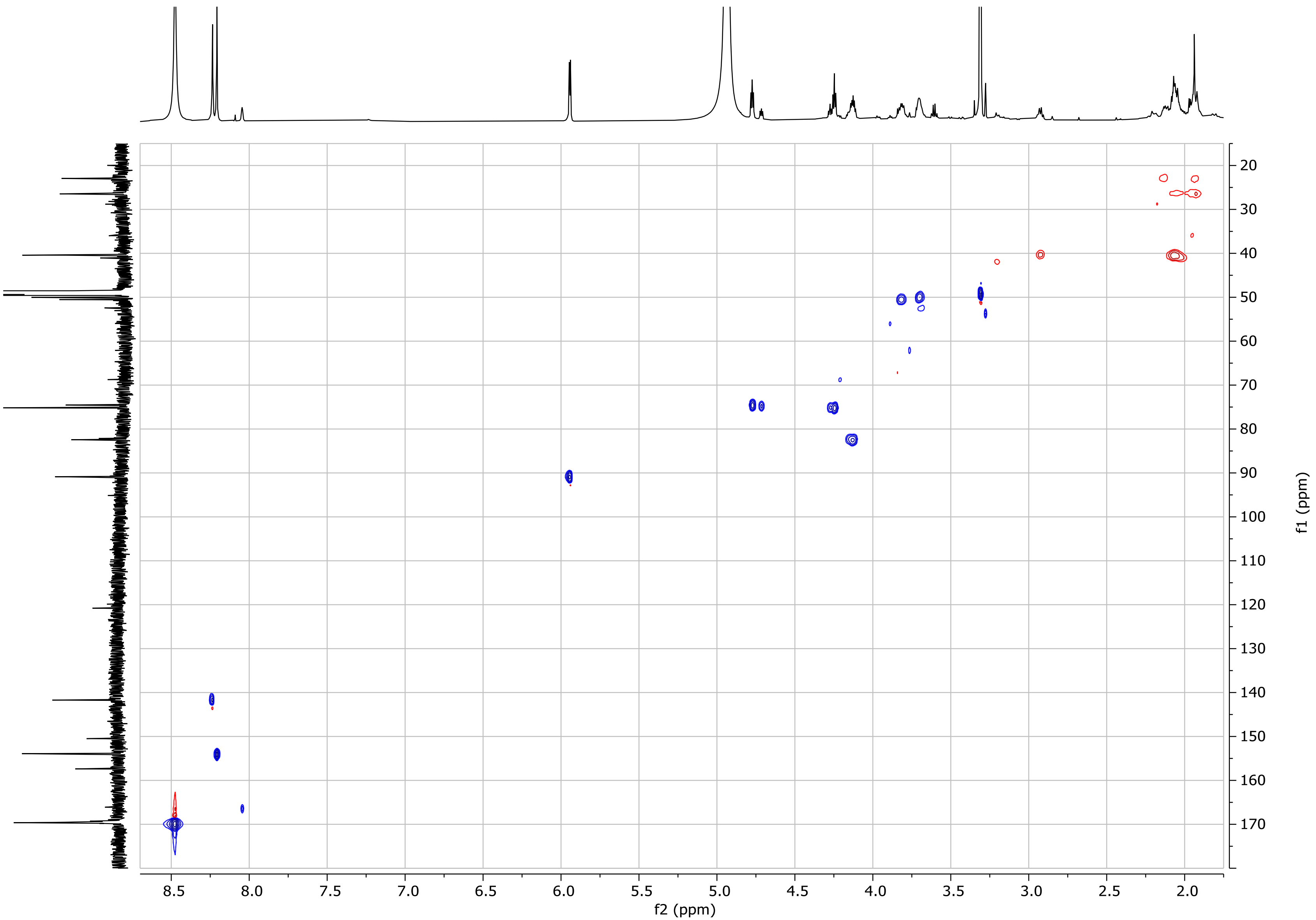

**Figure S16. HSQC spectrum of sinefungin lactam (2) (methanol-*d₄*).**

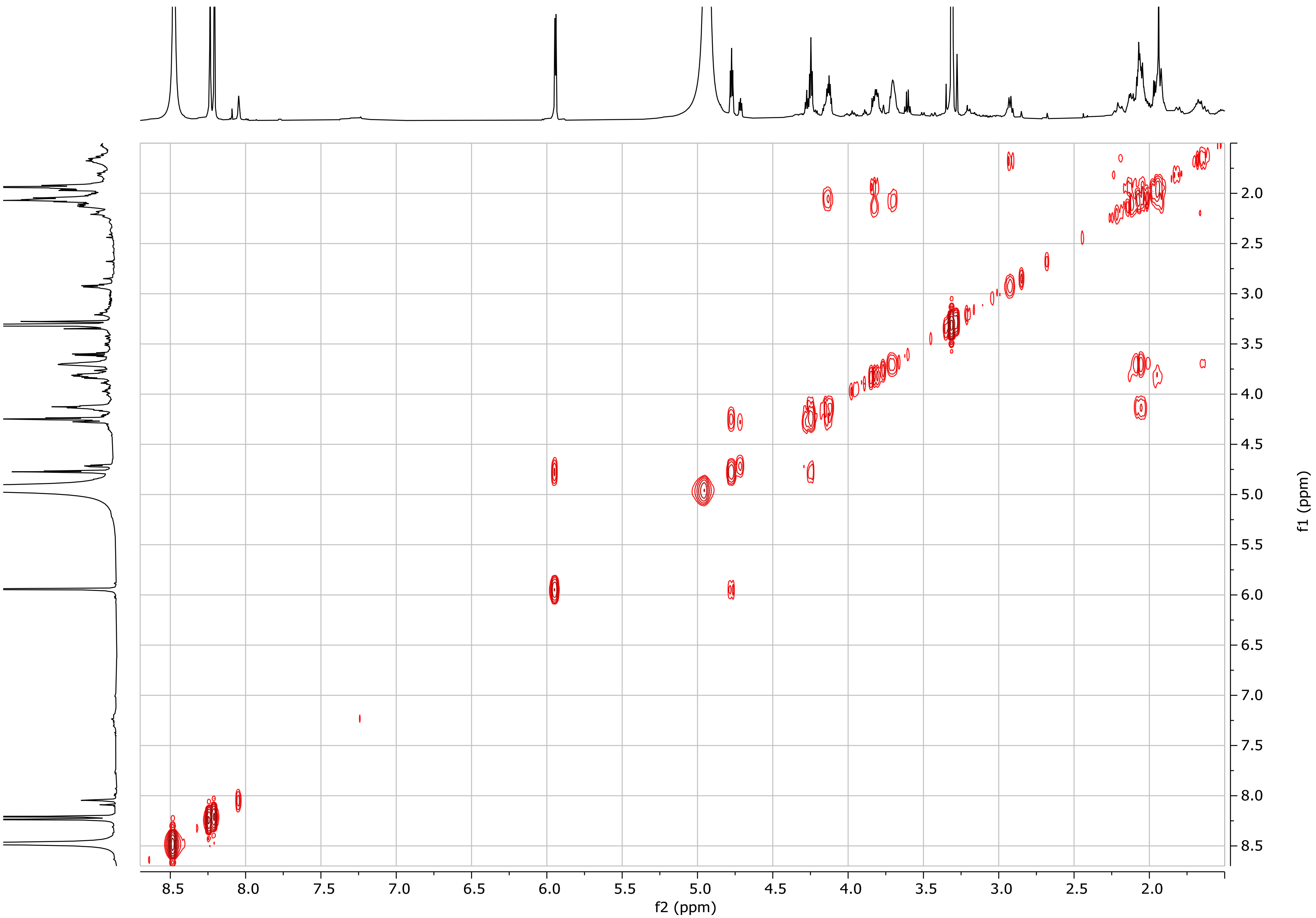

**Figure S17. COSY spectrum of sinefungin lactam (2) (methanol-*d*$_4$).**

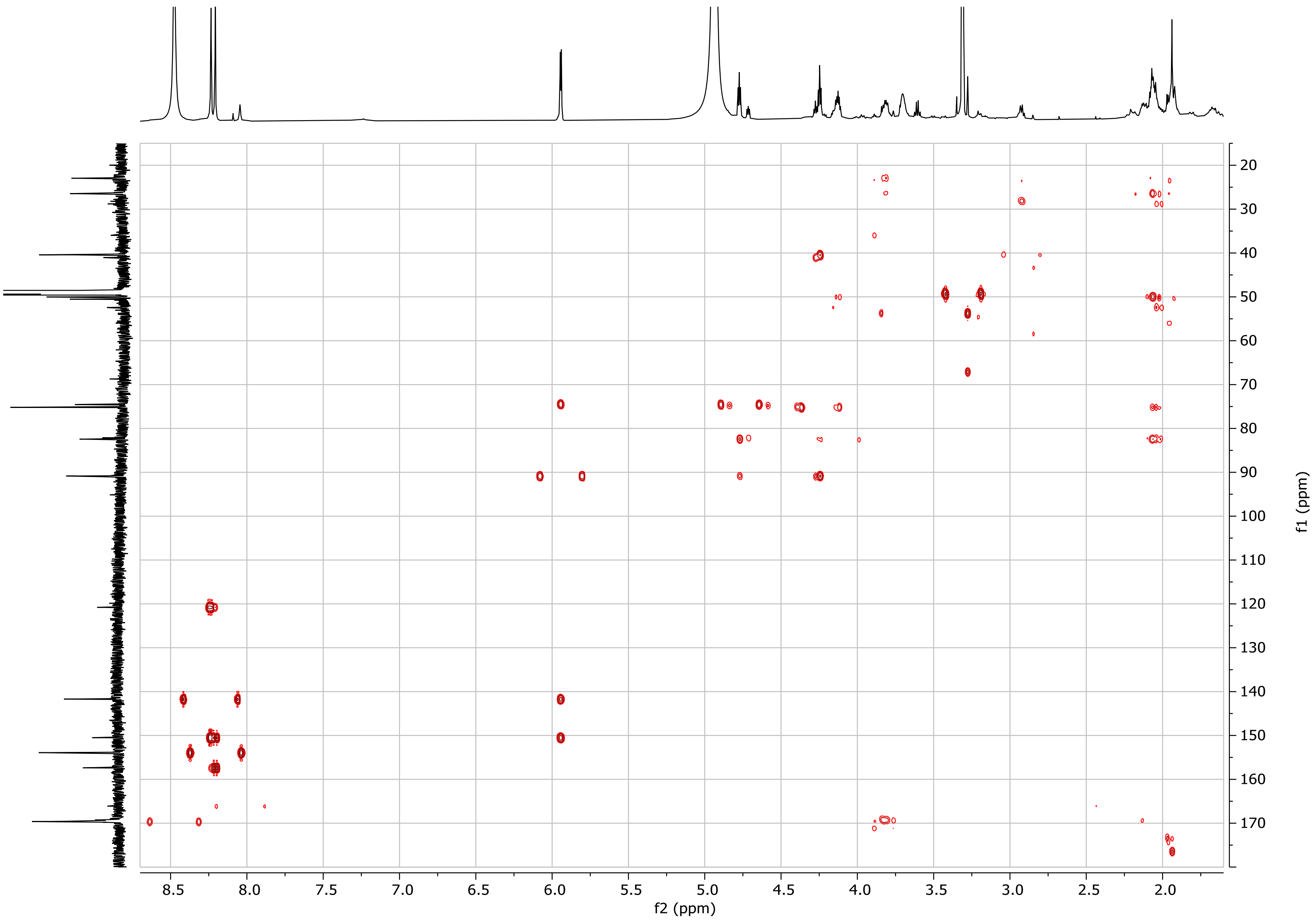

**Figure S18. HMBC spectrum of sinefungin lactam (2) (methanol-$d_4$).**

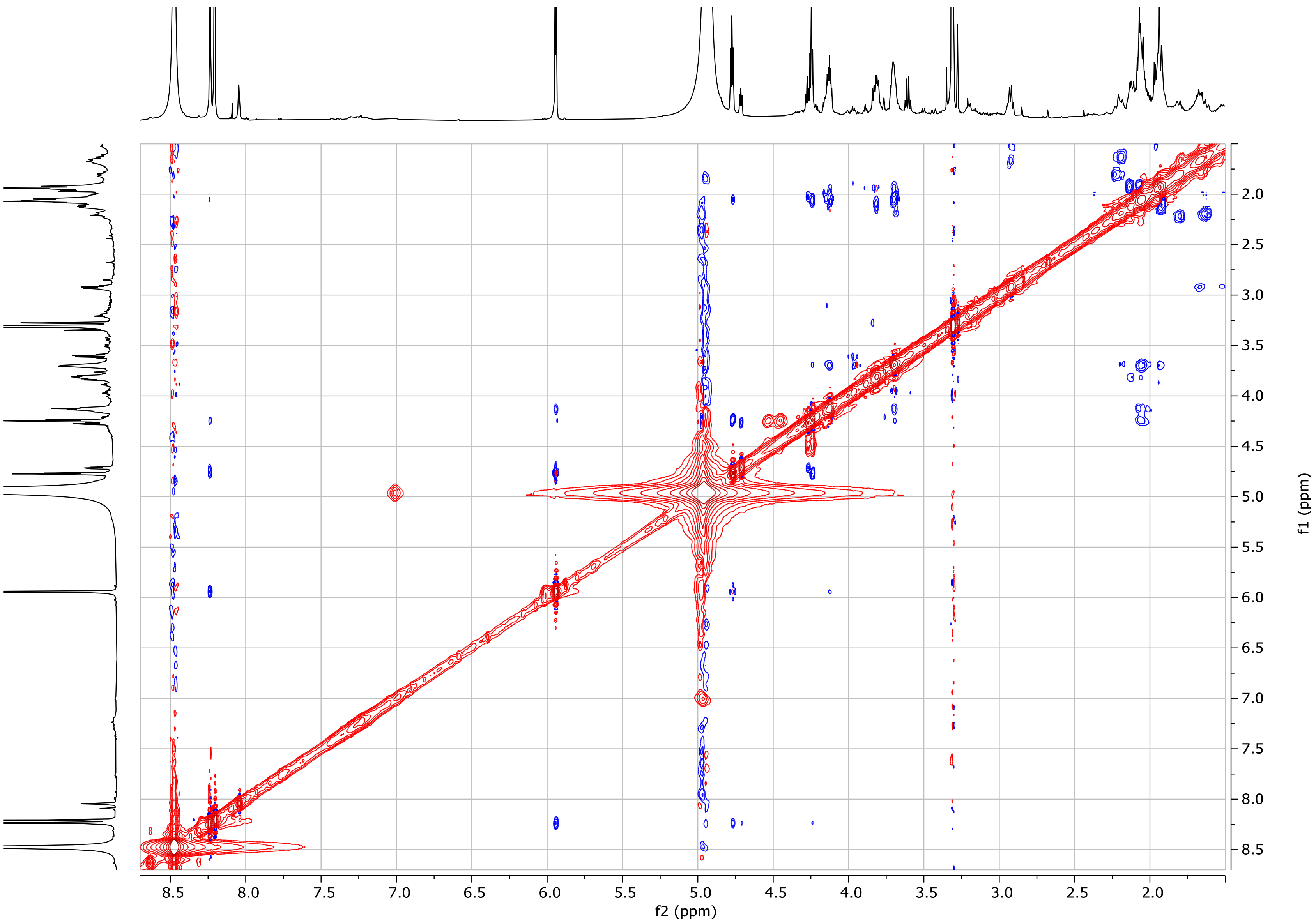

**Figure S19. NOESY spectrum of sinefungin lactam (2) (methanol-$d_4$).**

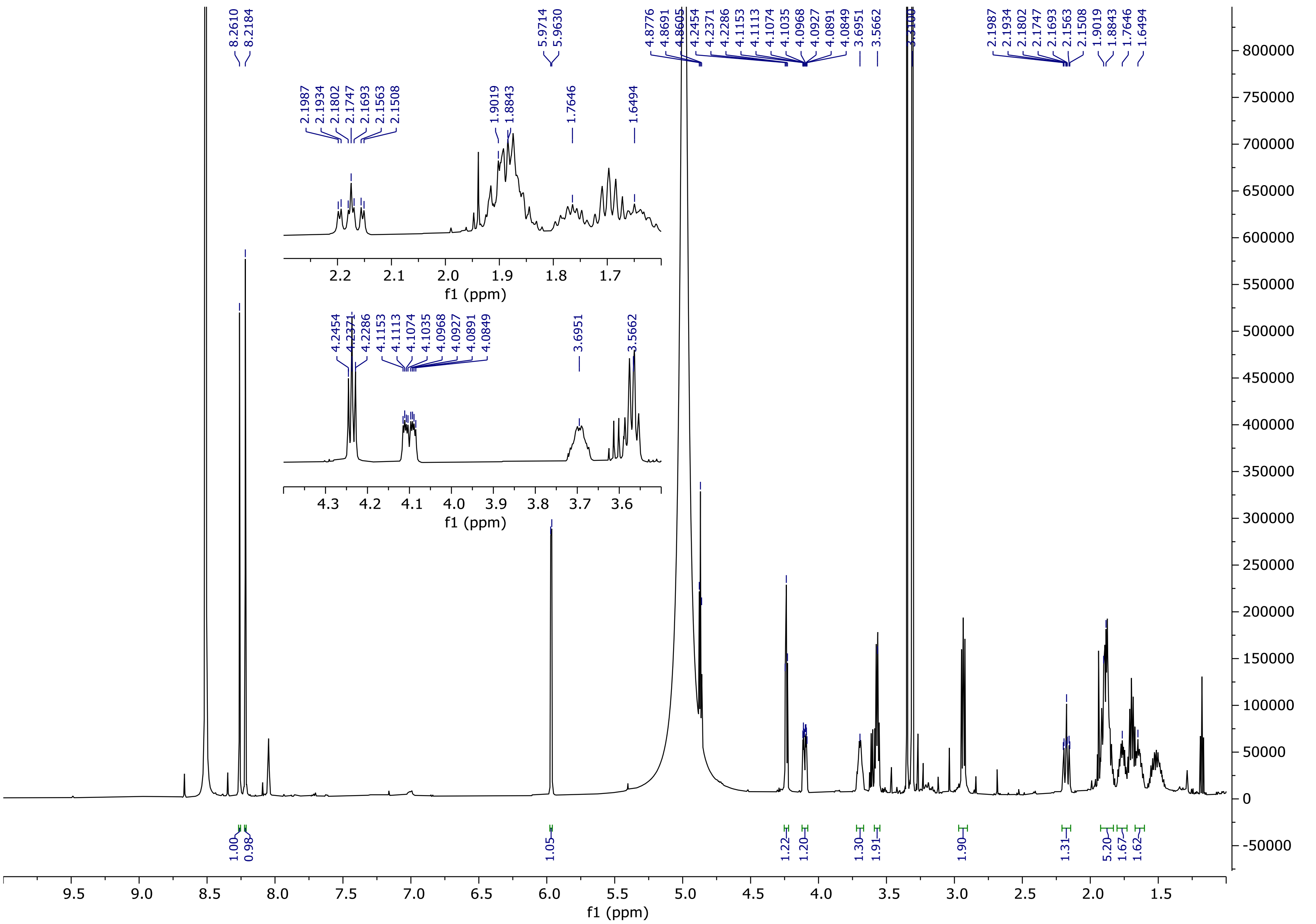

**Figure S20.** $^{1}H$ **NMR spectrum of guanidinyl-sinefungin (4) (methanol-*d*$_4$, 600 MHz).**

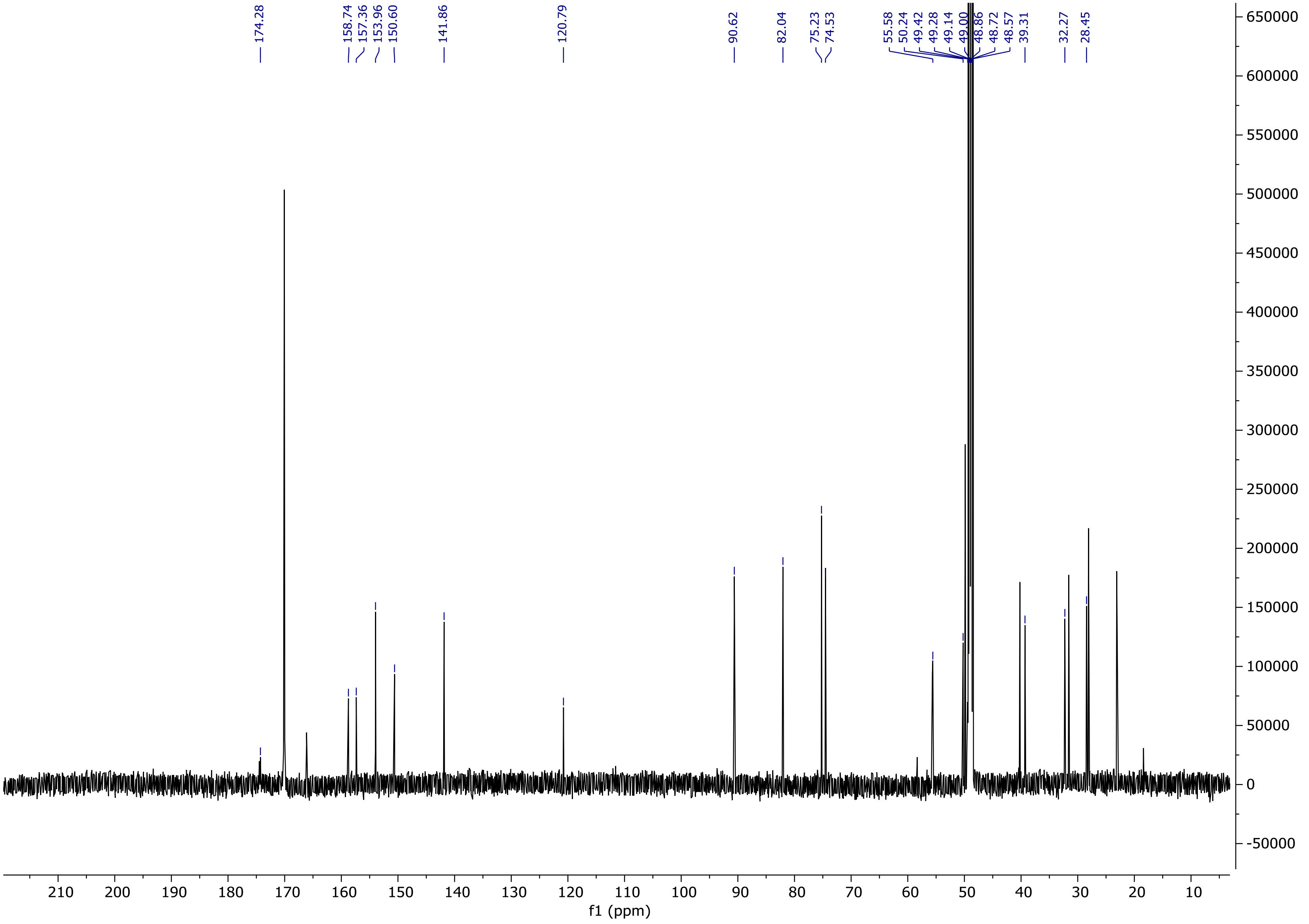

**Figure S21.** $^{13}$**C NMR spectrum of guanidinyl-sinefungin (4) (methanol-*d*$_4$, 150 MHz).**

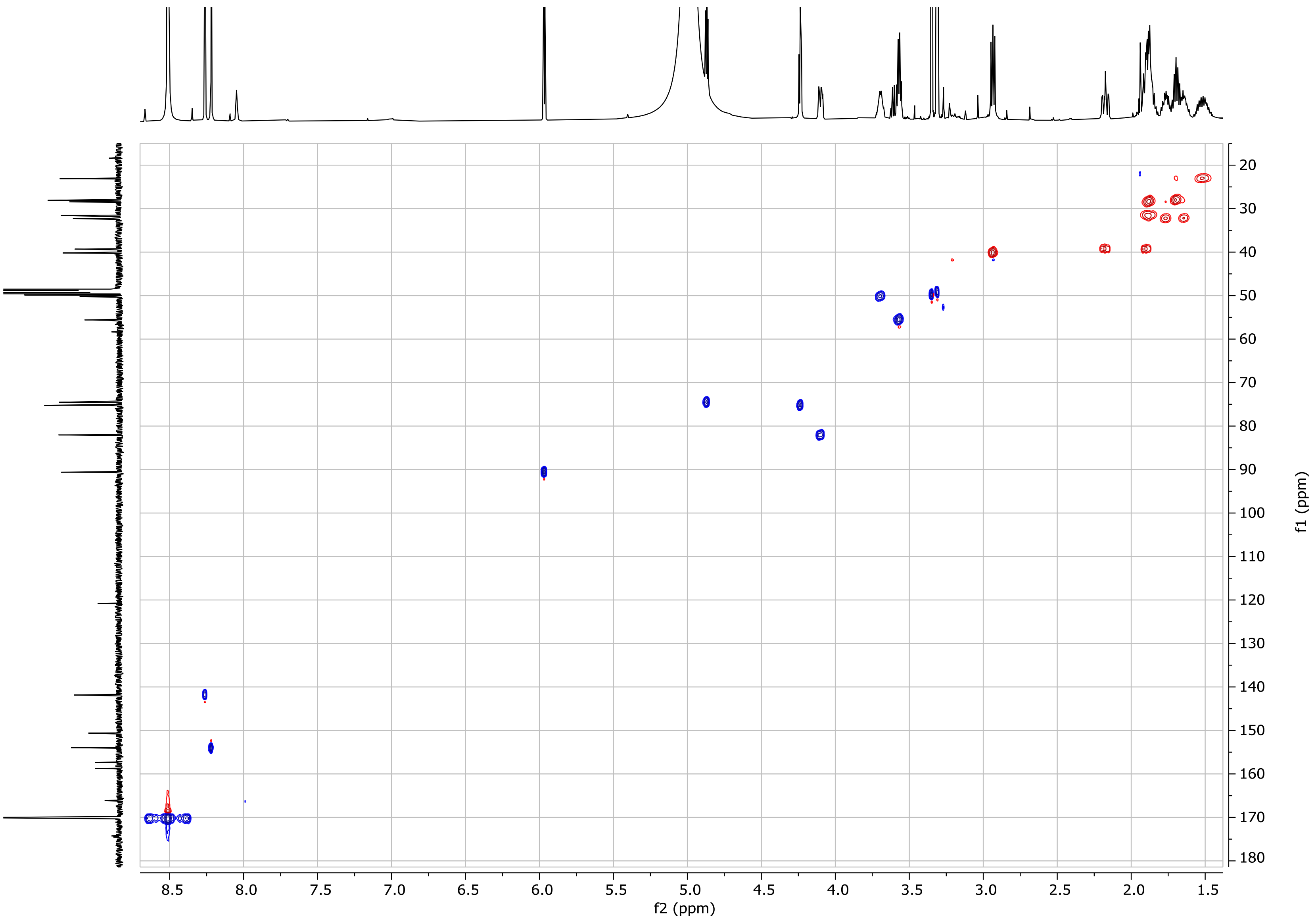

**Figure S22. HSQC spectrum of guanidinyl-sinefungin (4) (methanol-*d₄*).**

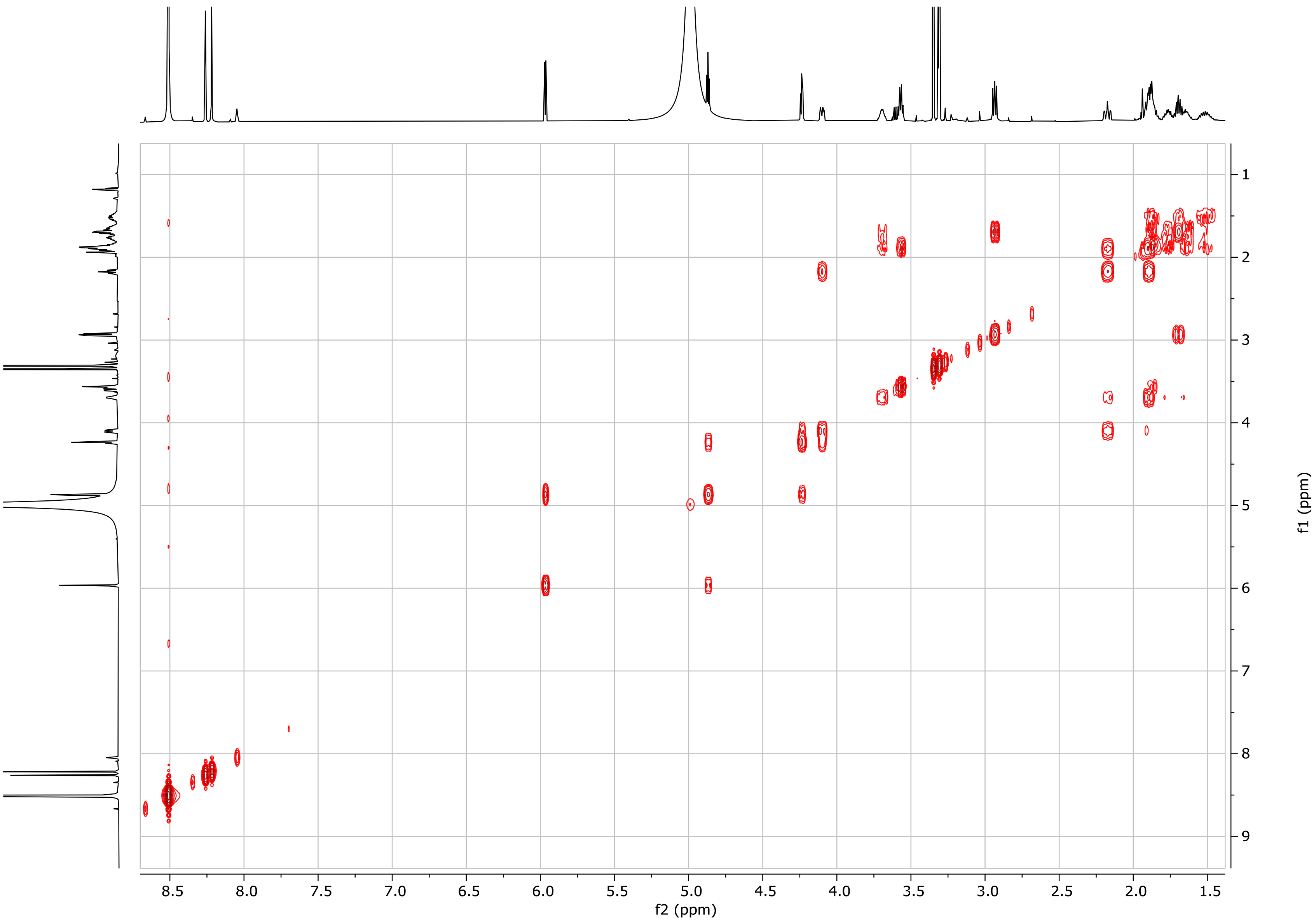

**Figure S23. COSY spectrum of guanidinyl-sinefungin (4) (methanol-*d*$_4$).**

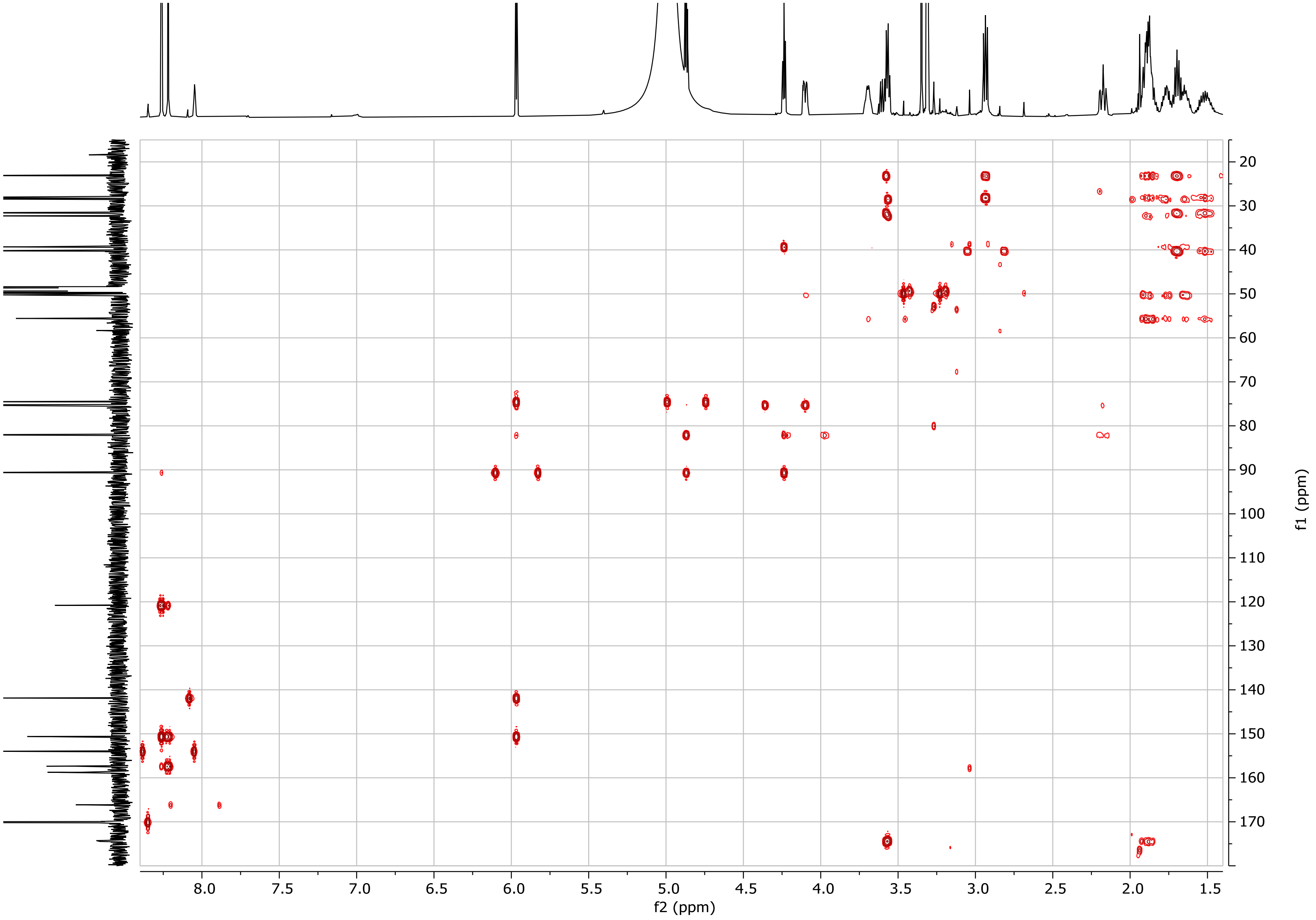

**Figure S24. HMBC spectrum of guanidinyl-sinefungin (4) (methanol-$d_4$).**

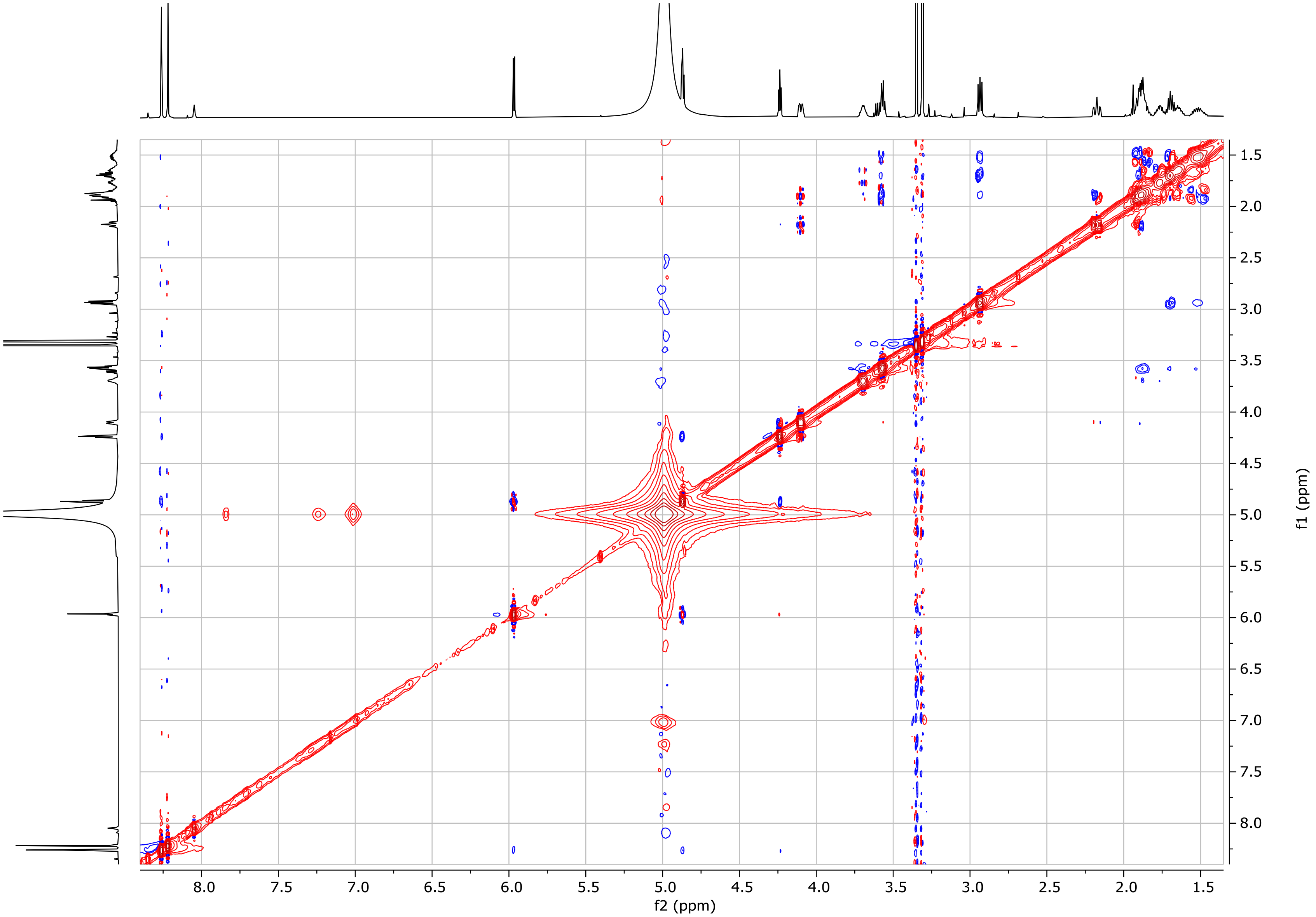

**Figure S25. NOESY spectrum of guanidinyl-sinefungin (4) (methanol-*d*$_4$).**

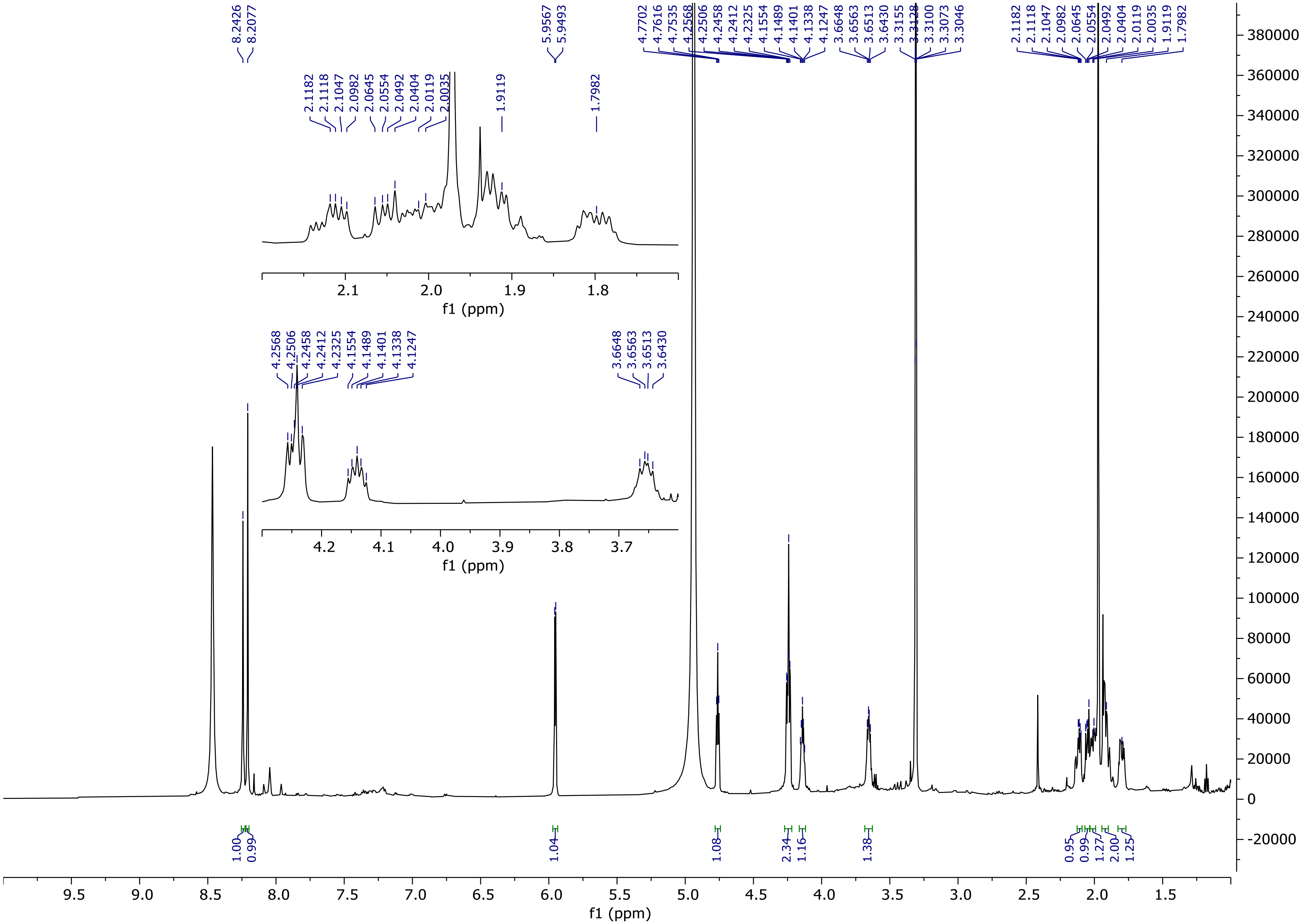

**Figure S26.** [1]**H NMR spectrum of guanidinyl-sinefungin lactam (5) (methanol-*d4*, 600 MHz).**

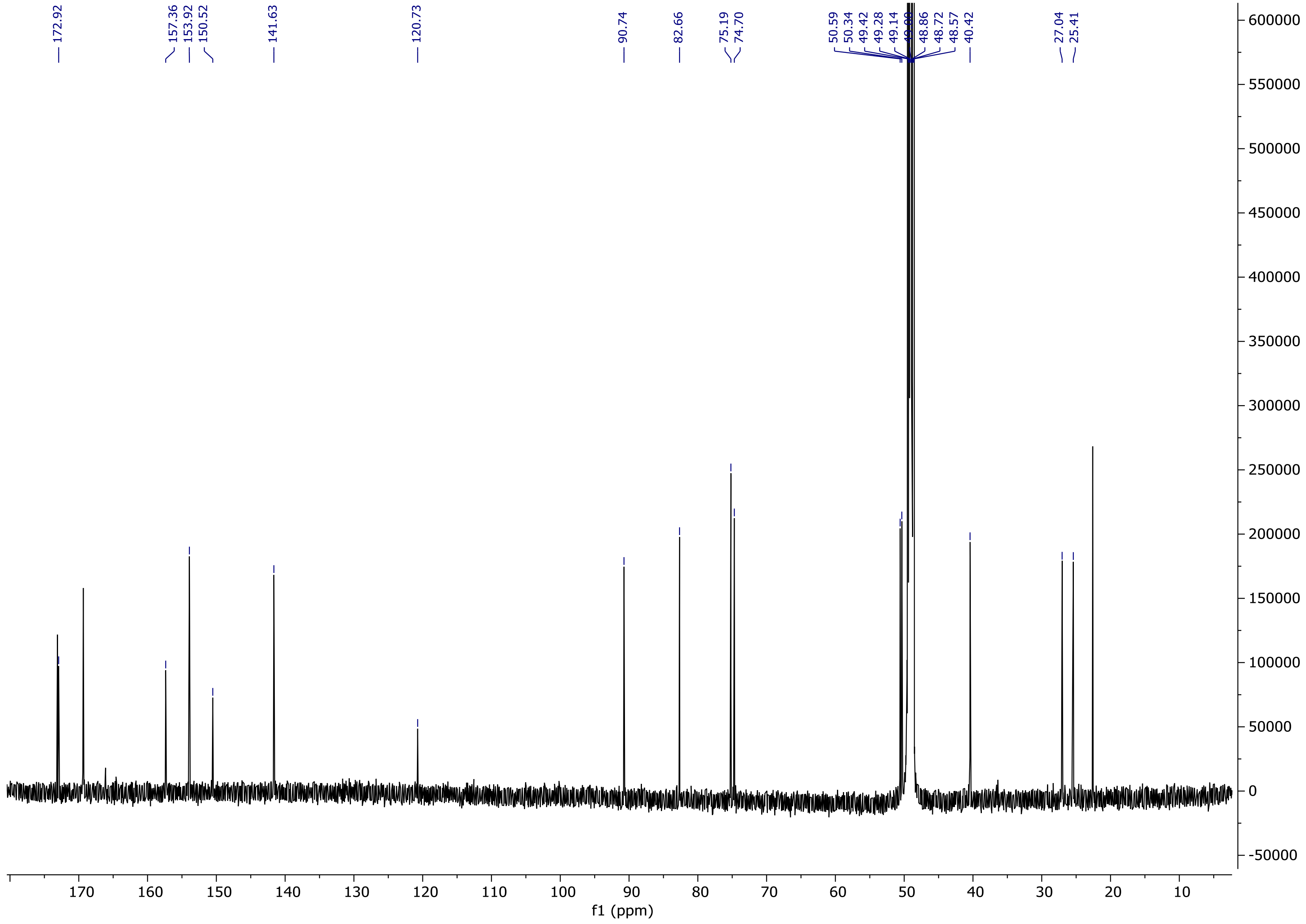

**Figure S27.** **$^{13}$C NMR spectrum of guanidinyl-sinefungin lactam (5) (methanol-*$d_4$*, 150 MHz).**

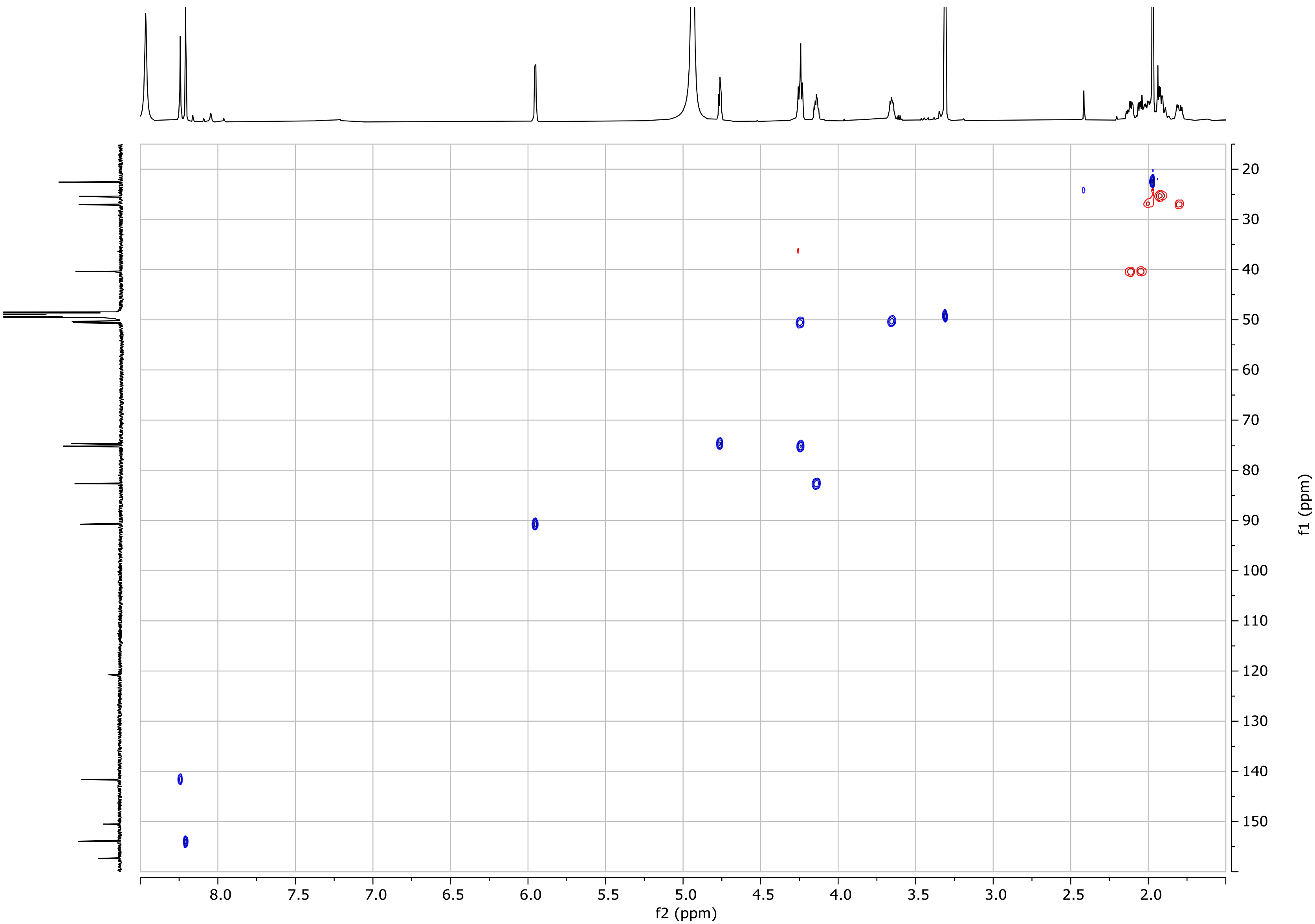

**Figure S28. HSQC spectrum of guanidinyl-sinefungin lactam (5) (methanol-*d*$_4$).**

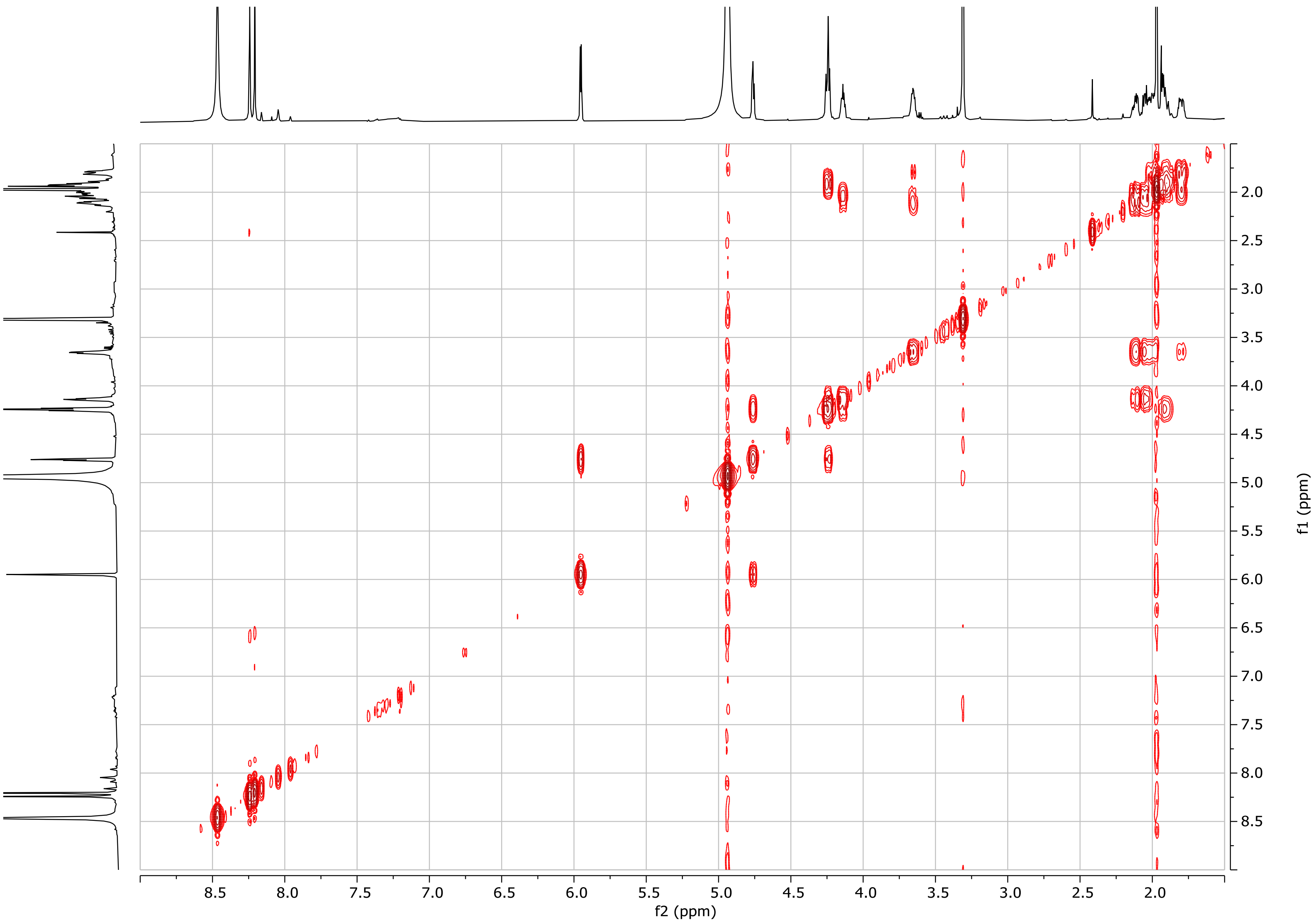

**Figure S29. COSY spectrum of guanidinyl-sinefungin lactam (5) (methanol-*d*$_4$).**

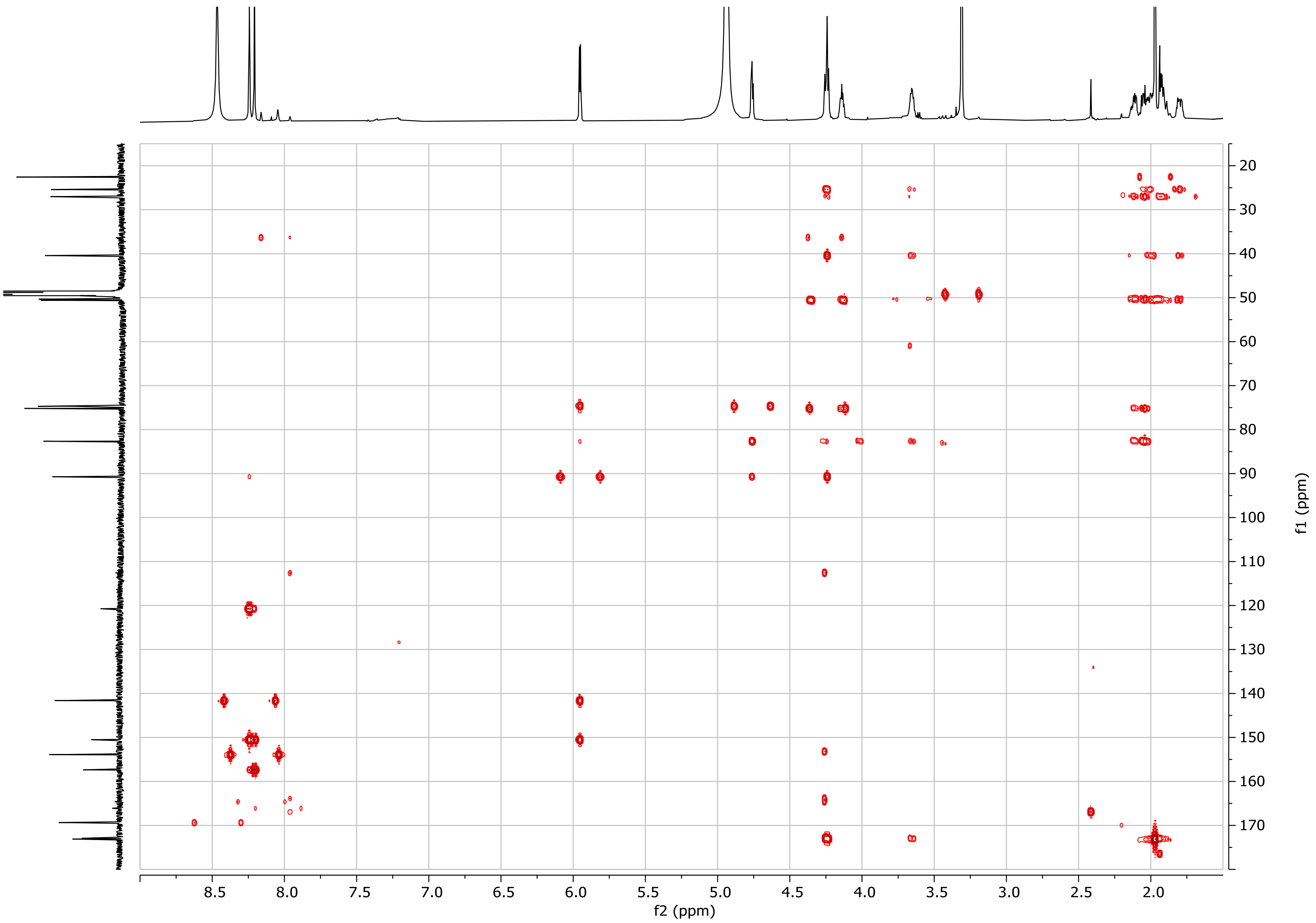

**Figure S30. HMBC spectrum of guanidinyl-sinefungin lactam (5) (methanol-*d*$_4$).**

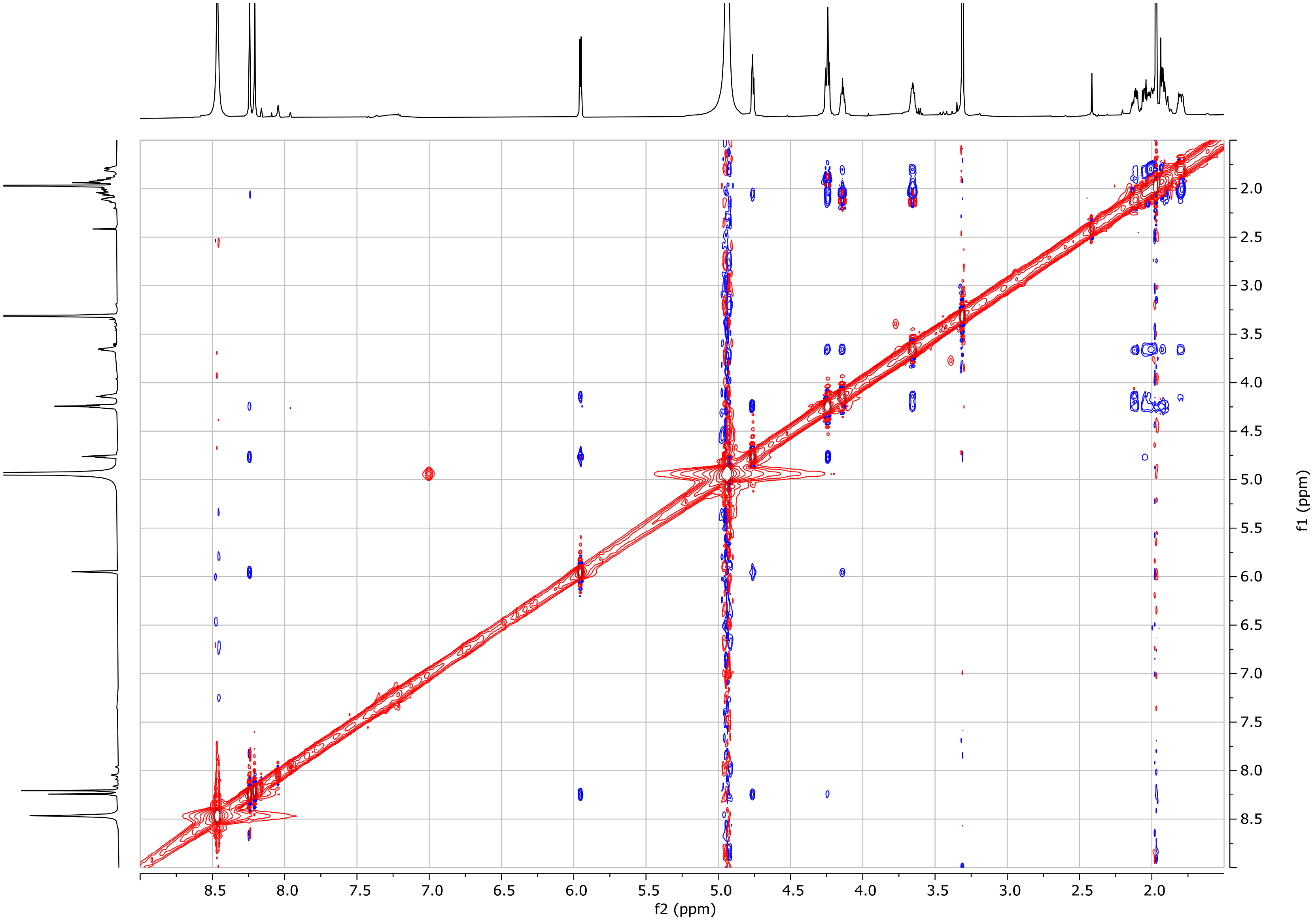

**Figure S31. NOESY spectrum of guanidinyl-sinefungin lactam (5) (methanol-*d₄*).**

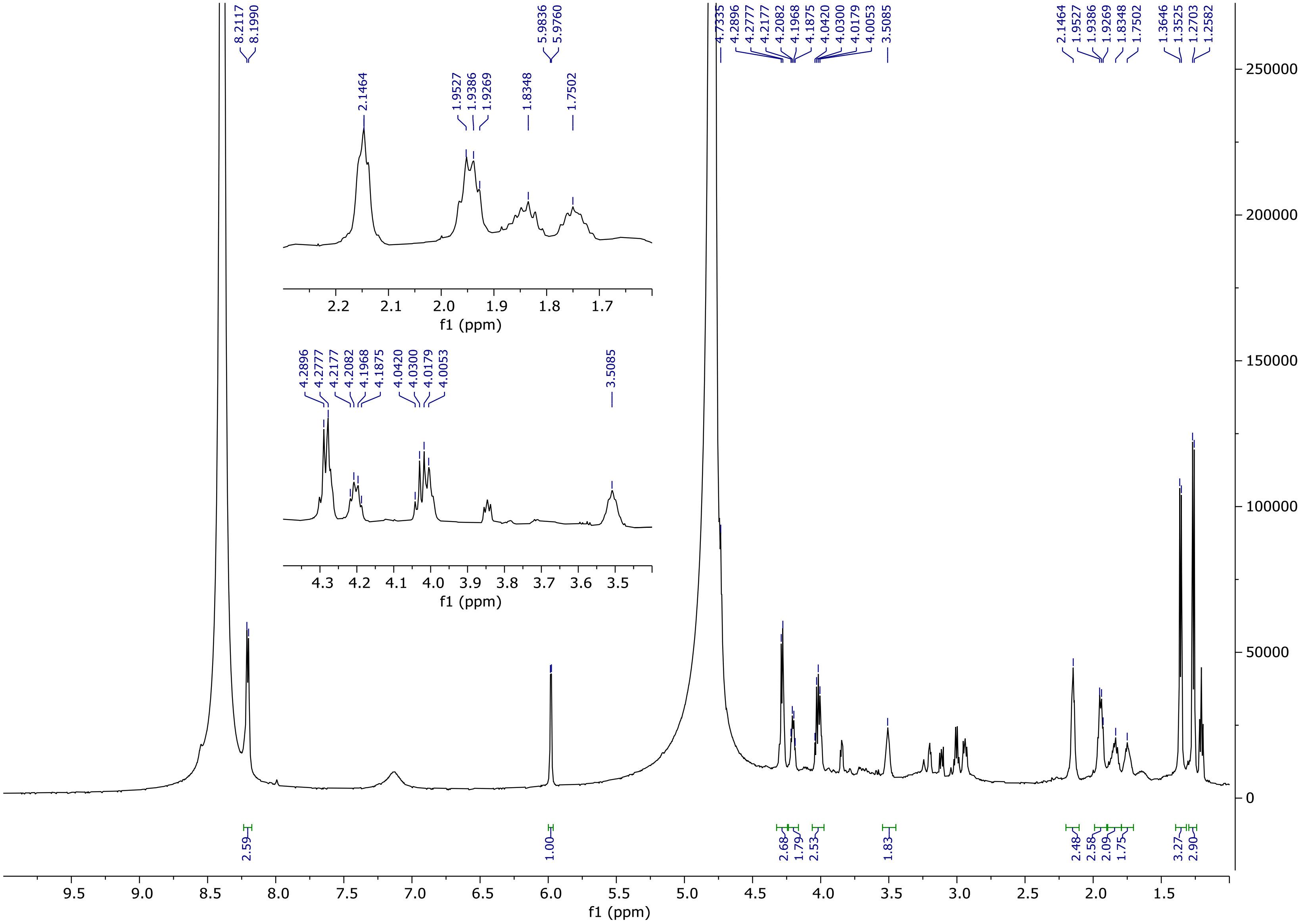

**Figure S32.** **¹H NMR spectrum of sinefungin AA (7) ($D_2O$, 600 MHz).**

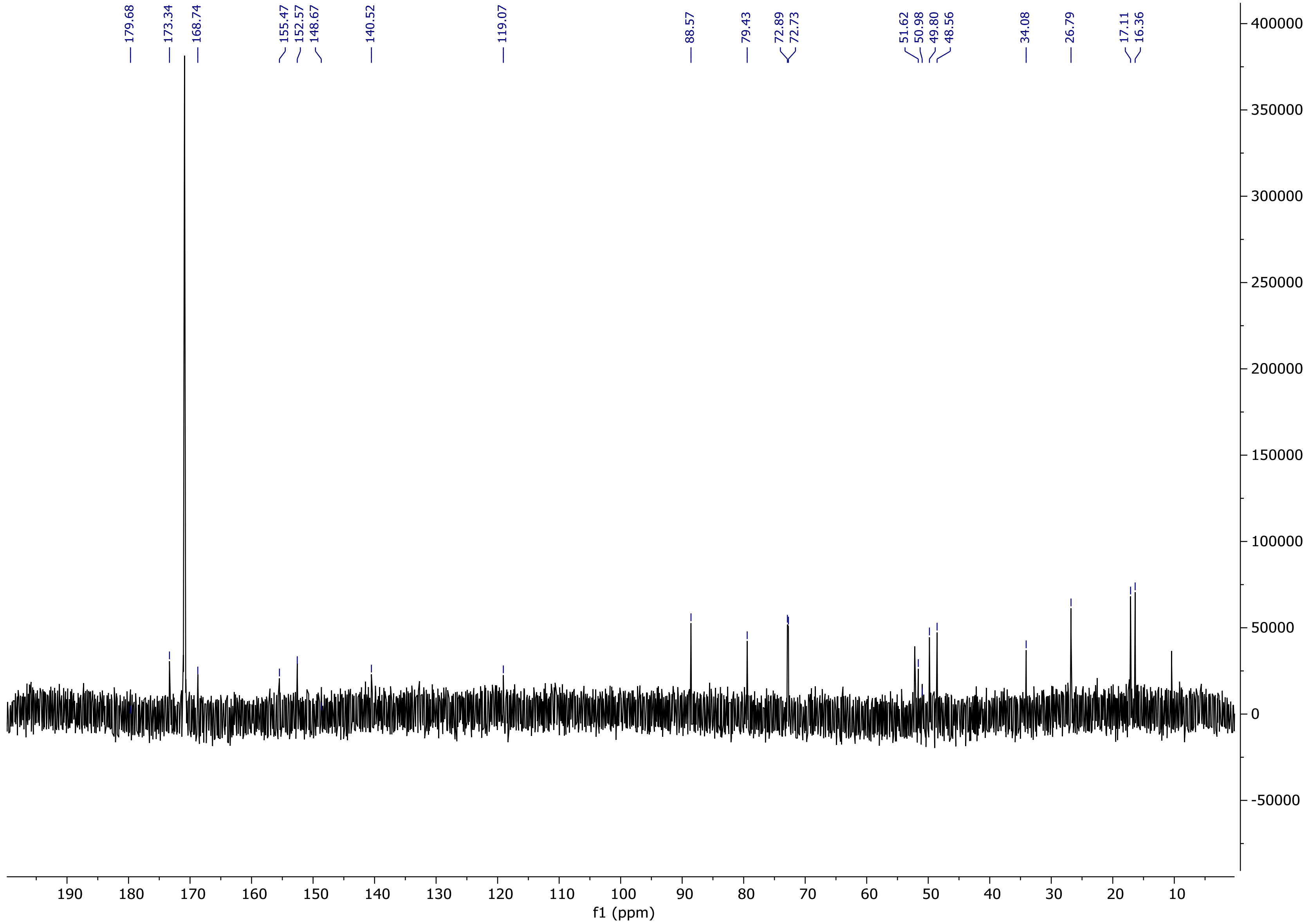

**Figure S33.** $^{13}$**C NMR spectrum of sinefungin AA (7) (D$_2$O, 150 MHz).**

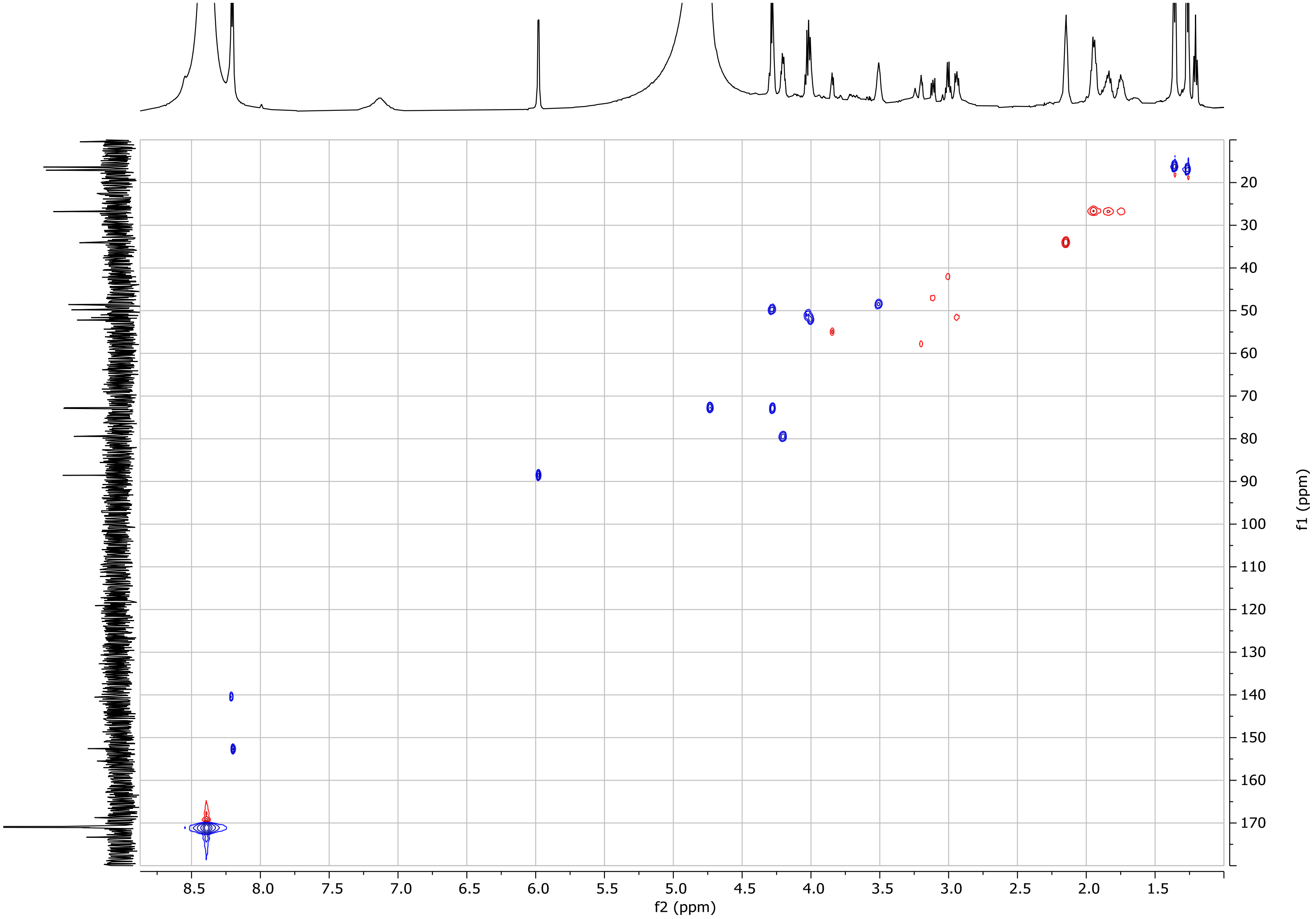

**Figure S34. HSQC spectrum of sinefungin AA (7) ($D_2O$).**

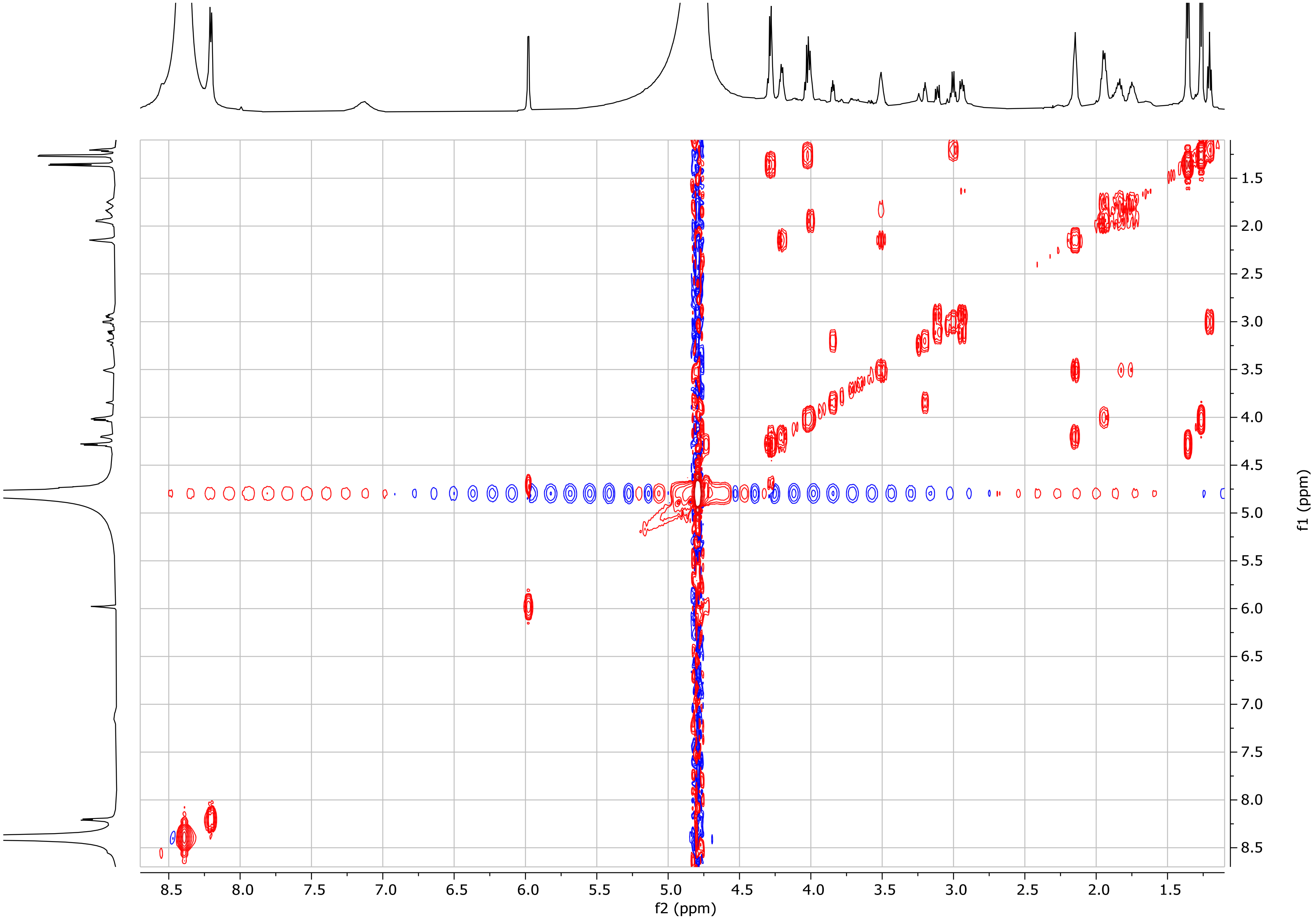

**Figure S35. COSY spectrum of sinefungin AA (7) ($D_2O$).**

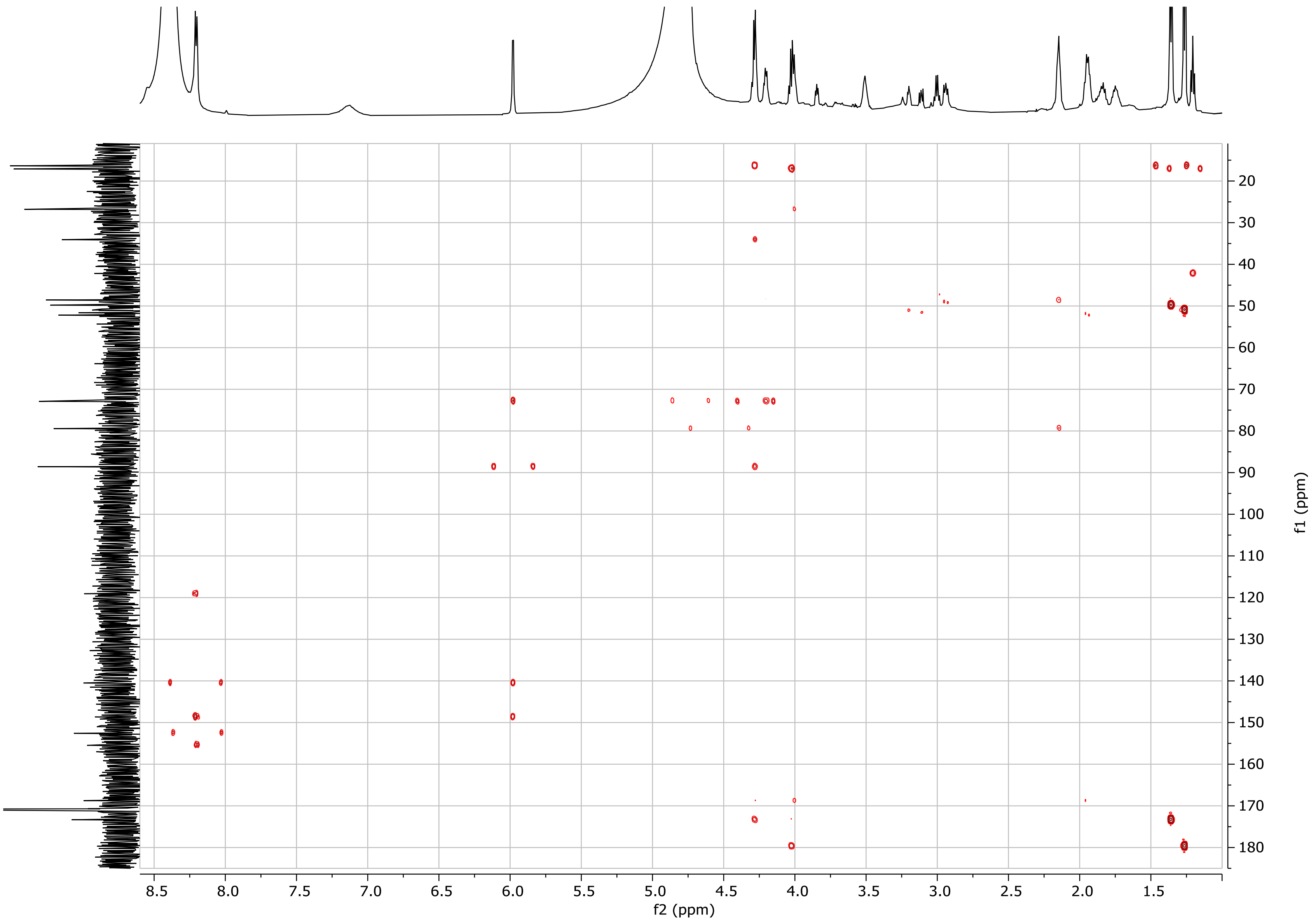

**Figure S36. HMBC spectrum of sinefungin AA (7) ($D_2O$).**

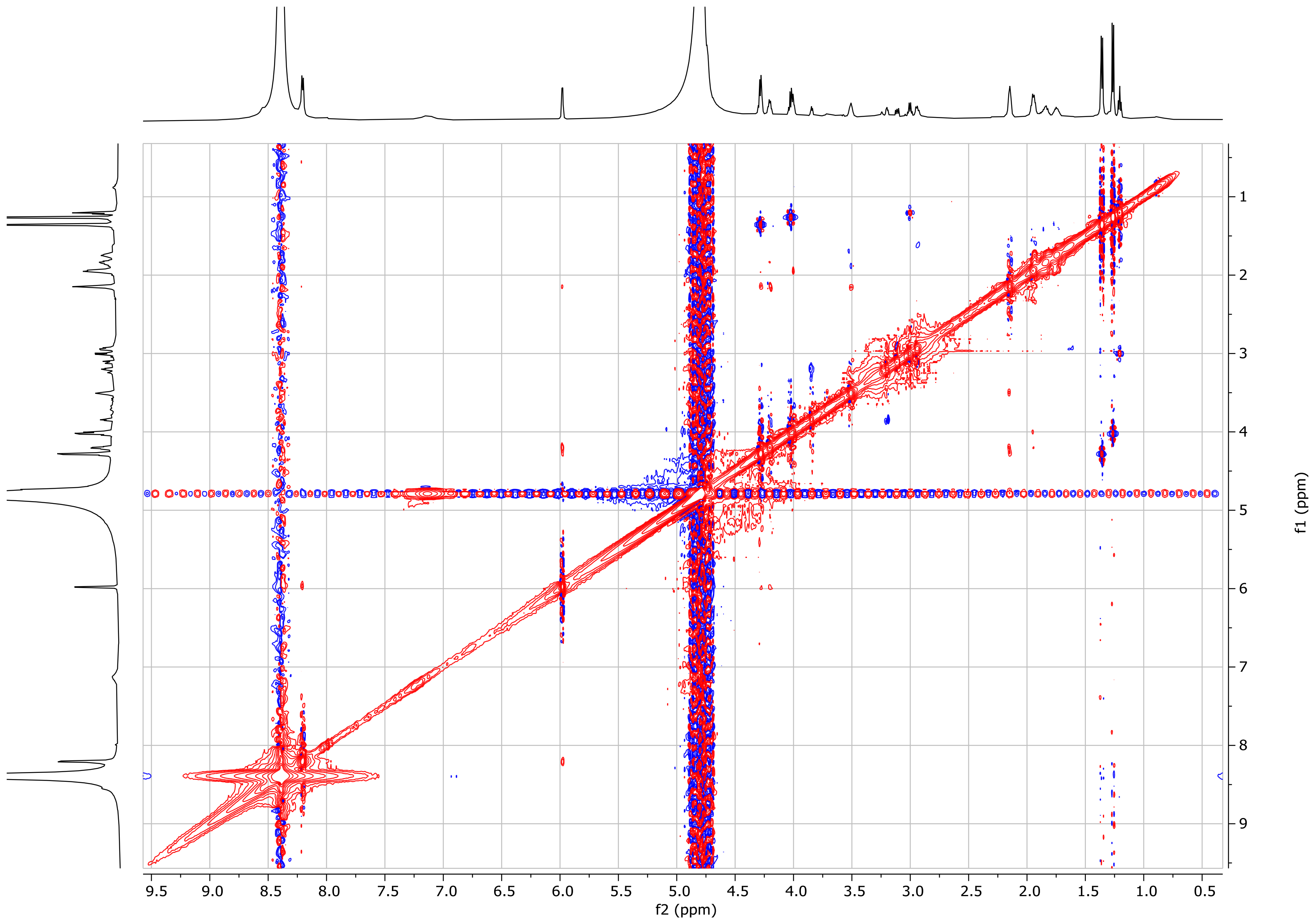

**Figure S37. NOESY spectrum of sinefungin AA (7) ($D_2O$).**

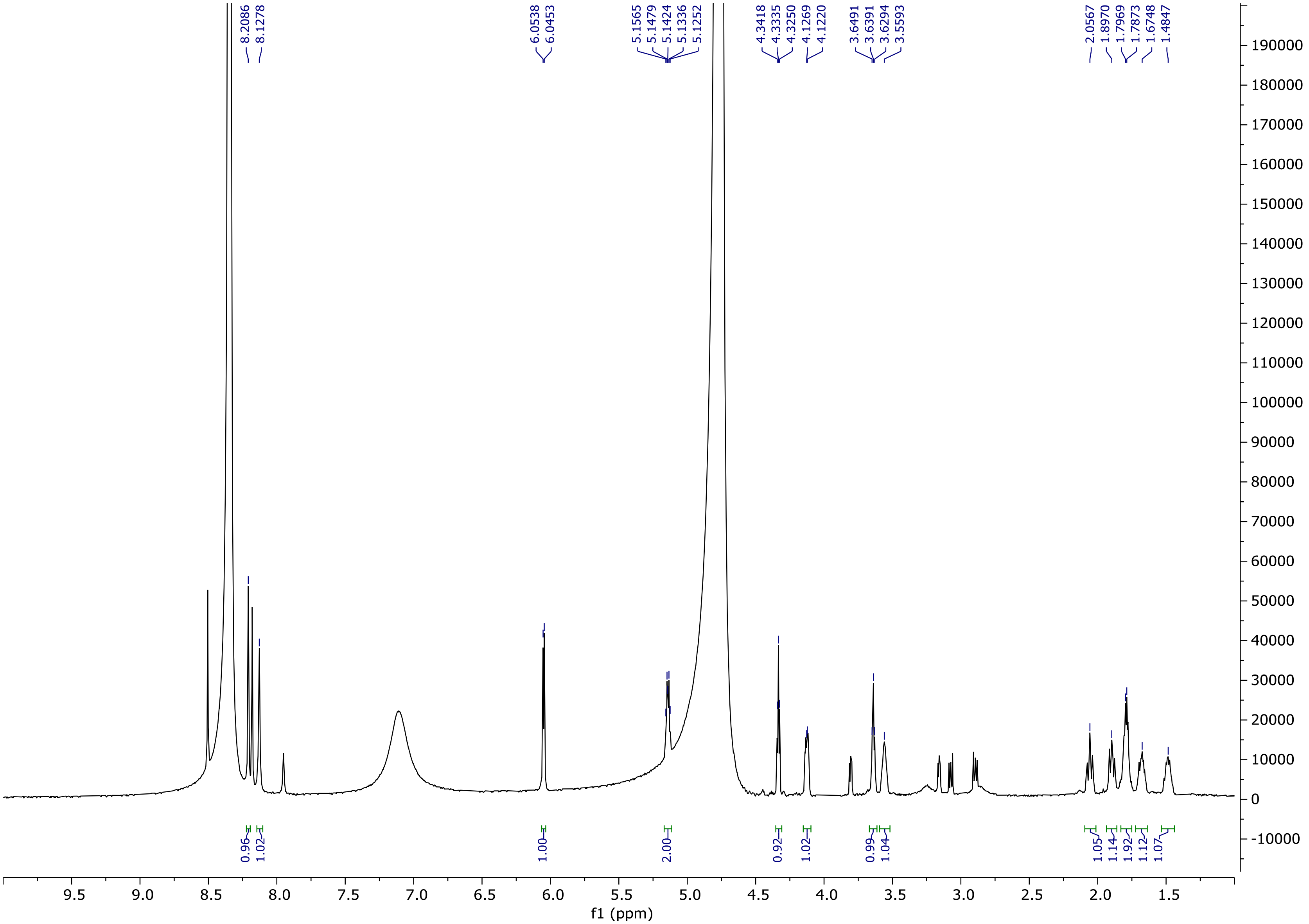

**Figure S38.** **$^{1}$H NMR spectrum of 2'-phosphoguanidinyl-sinefungin (8) ($D_2O$, 600 MHz).**

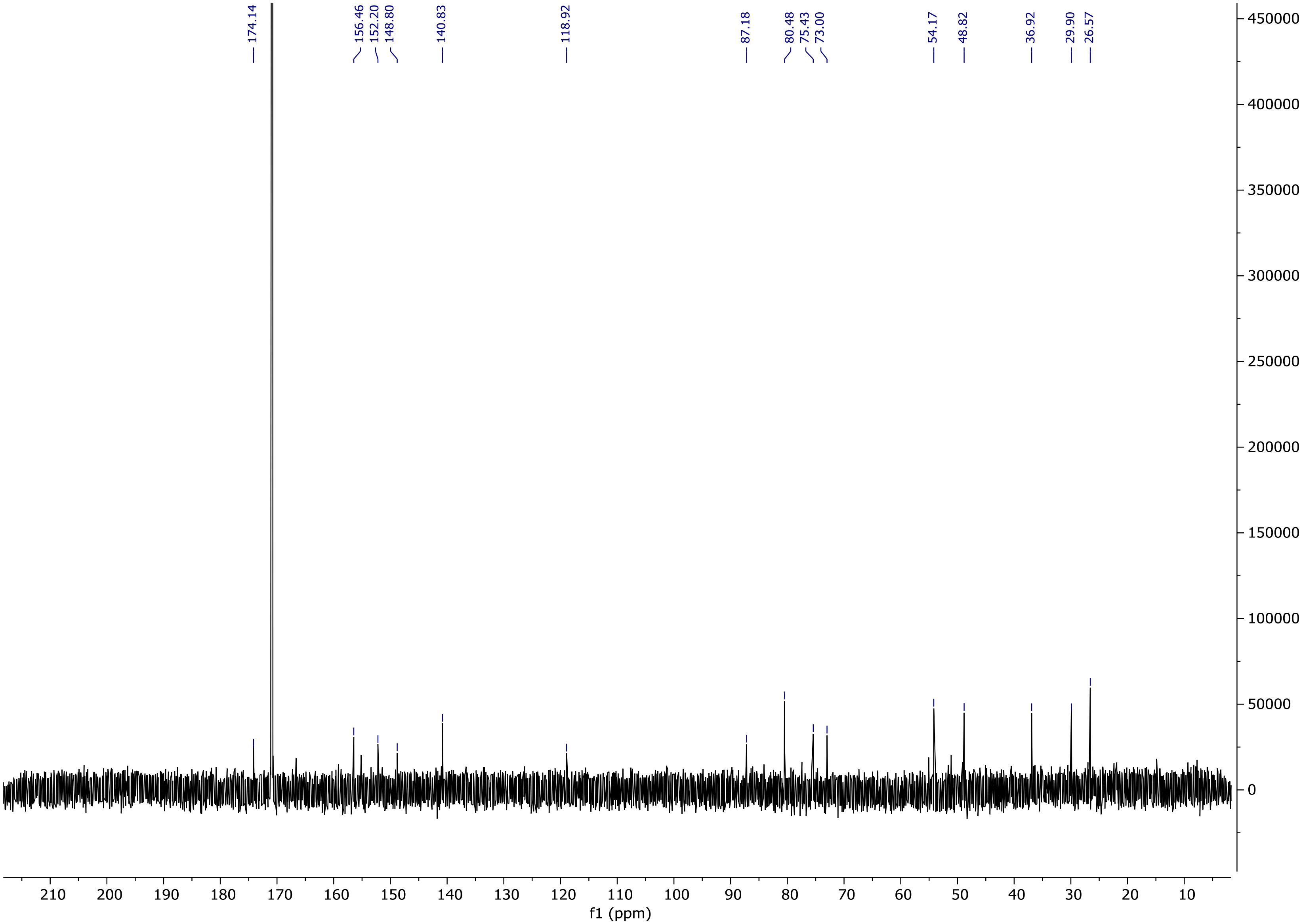

**Figure S39. $^{13}$C NMR spectrum of 2'-phosphoguanidinyl-sinefungin (8) ($D_2O$, 150 MHz).**

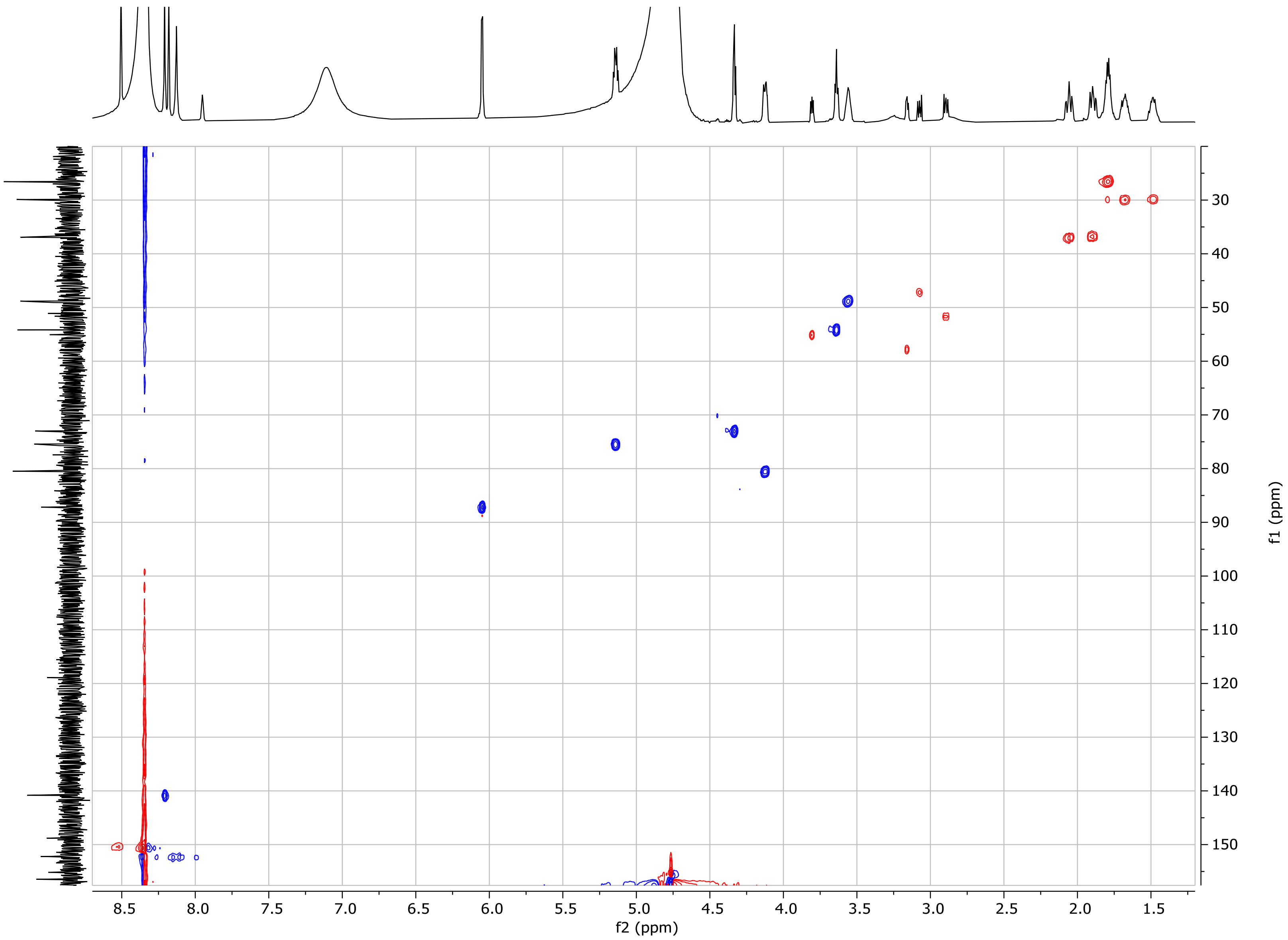

**Figure S40. HSQC spectrum of 2'-phosphoguanidinyl-sinefungin (8) ($D_2O$).**

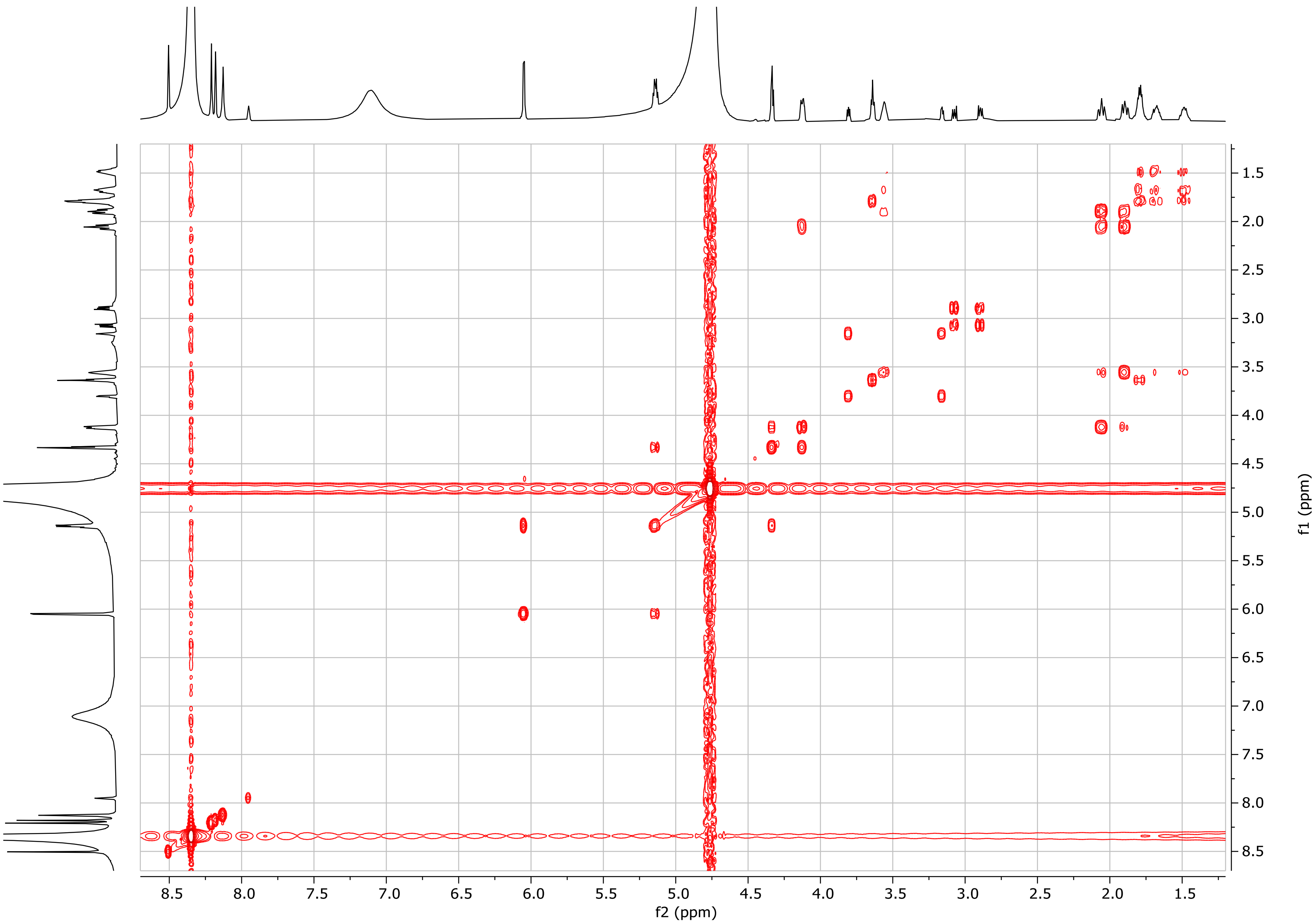

**Figure S41. COSY spectrum of 2'-phosphoguanidinyl-sinefungin (8) ($D_2O$).**

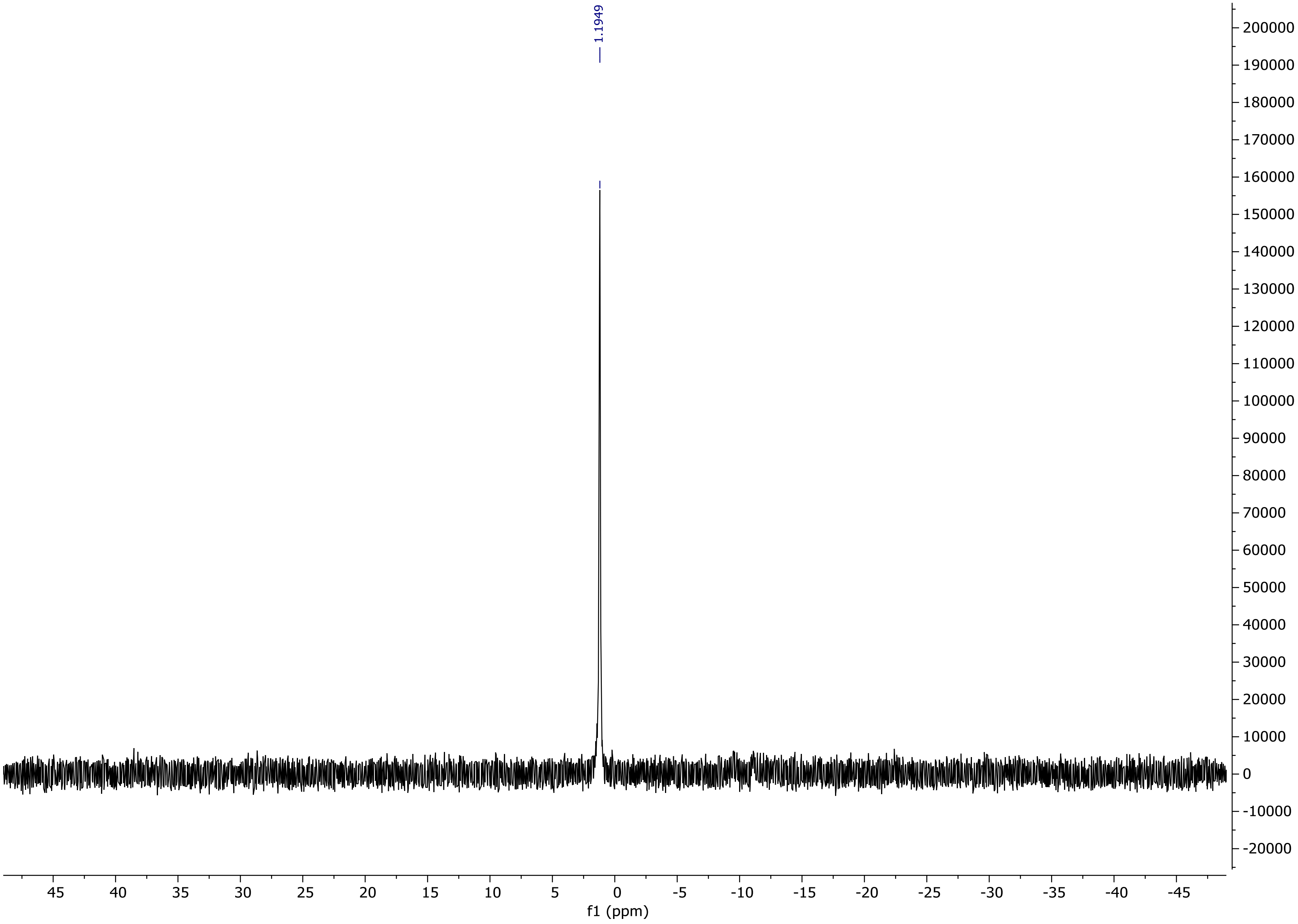

**Figure S42. $^{31}$P NMR spectrum of 2'-phosphoguanidinyl-sinefungin (8) ($D_2O$, 242 MHz).**

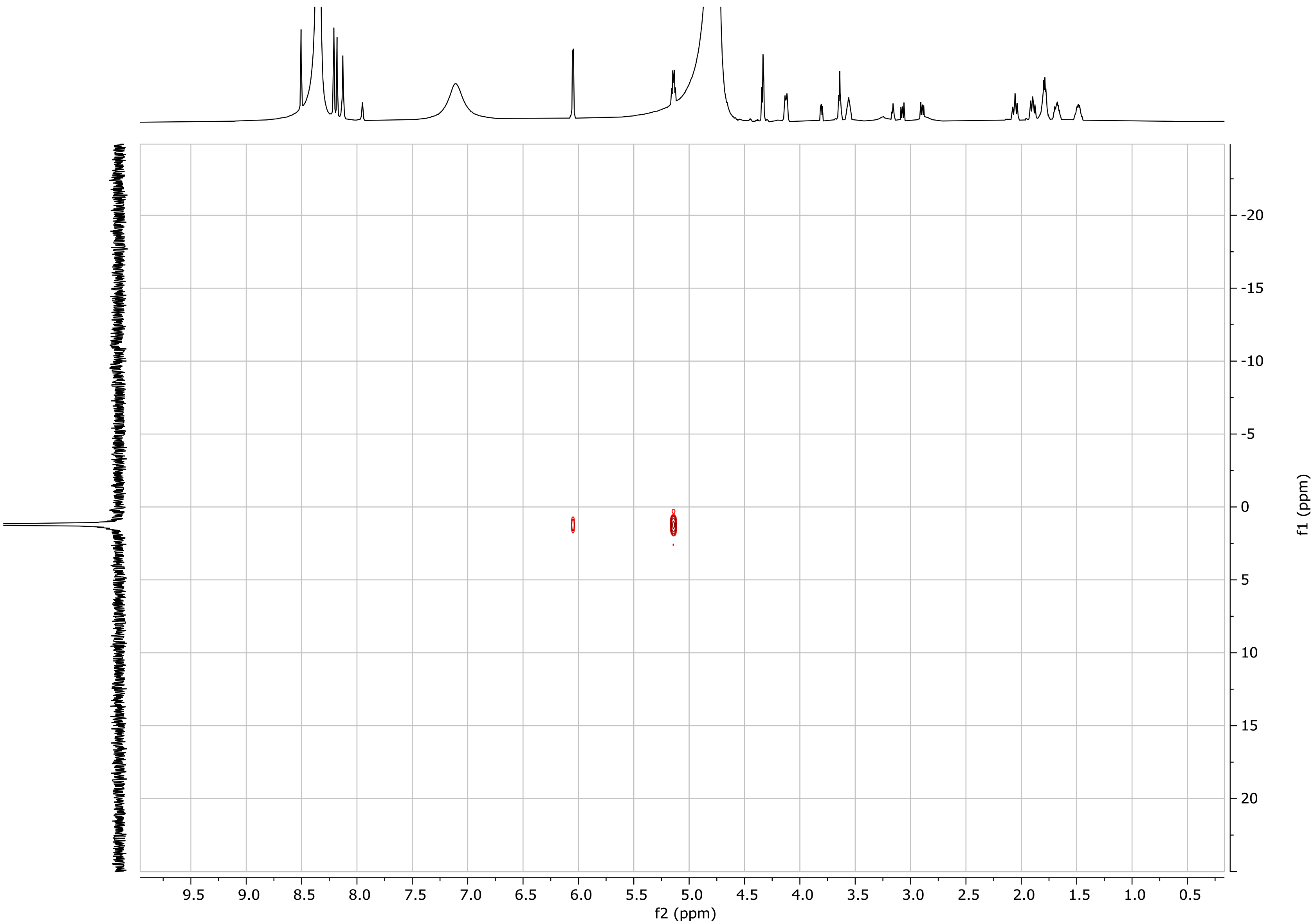

**Figure S43. $^{1}H$–$^{31}P$ HMBC spectrum of 2’-phosphoguanidinyl-sinefungin (8) ($D_2O$).**

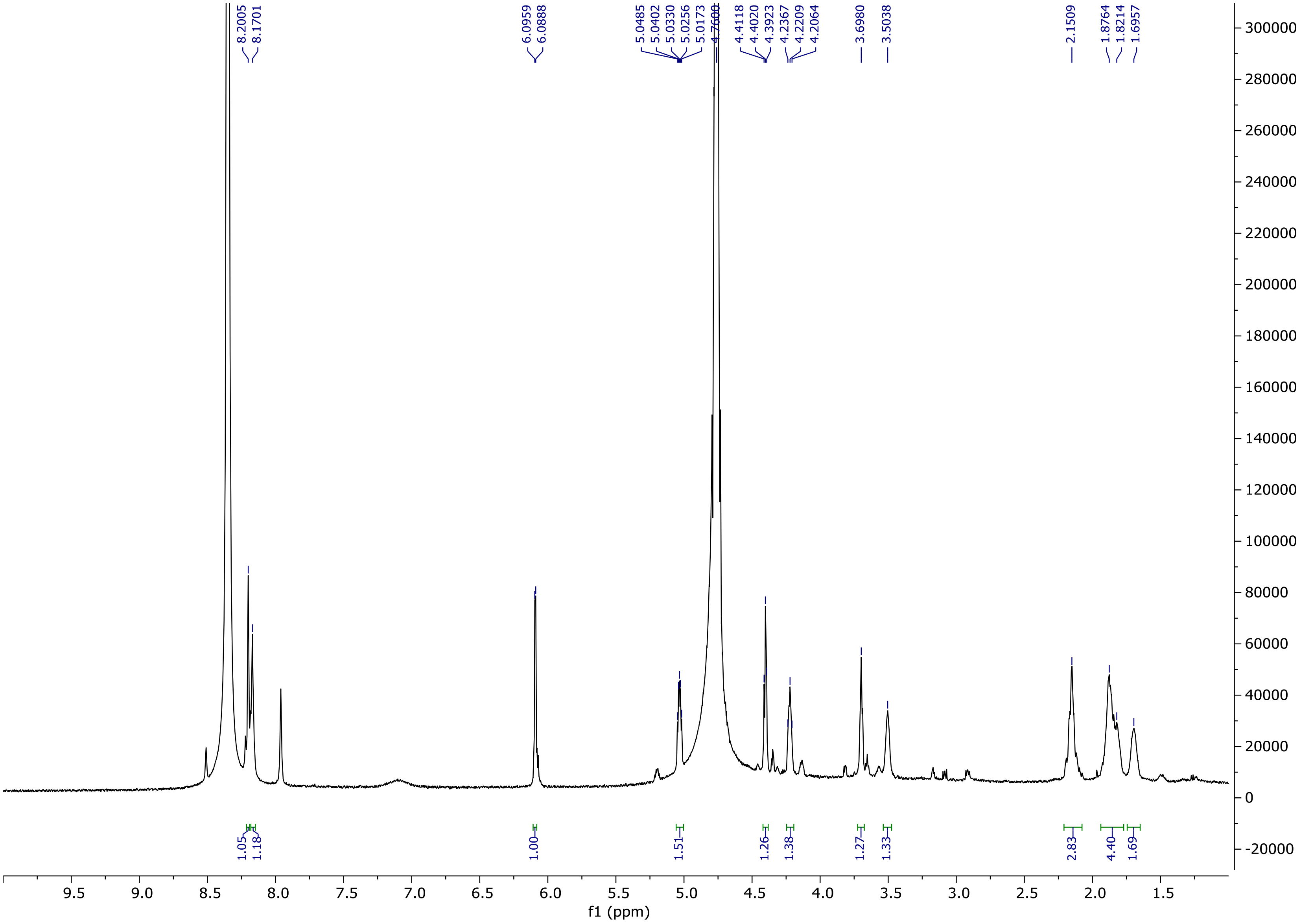

**Figure S44. [1]H NMR spectrum of 2'-phosphosinefungin (9) ($D_2O$, 150 MHz).**

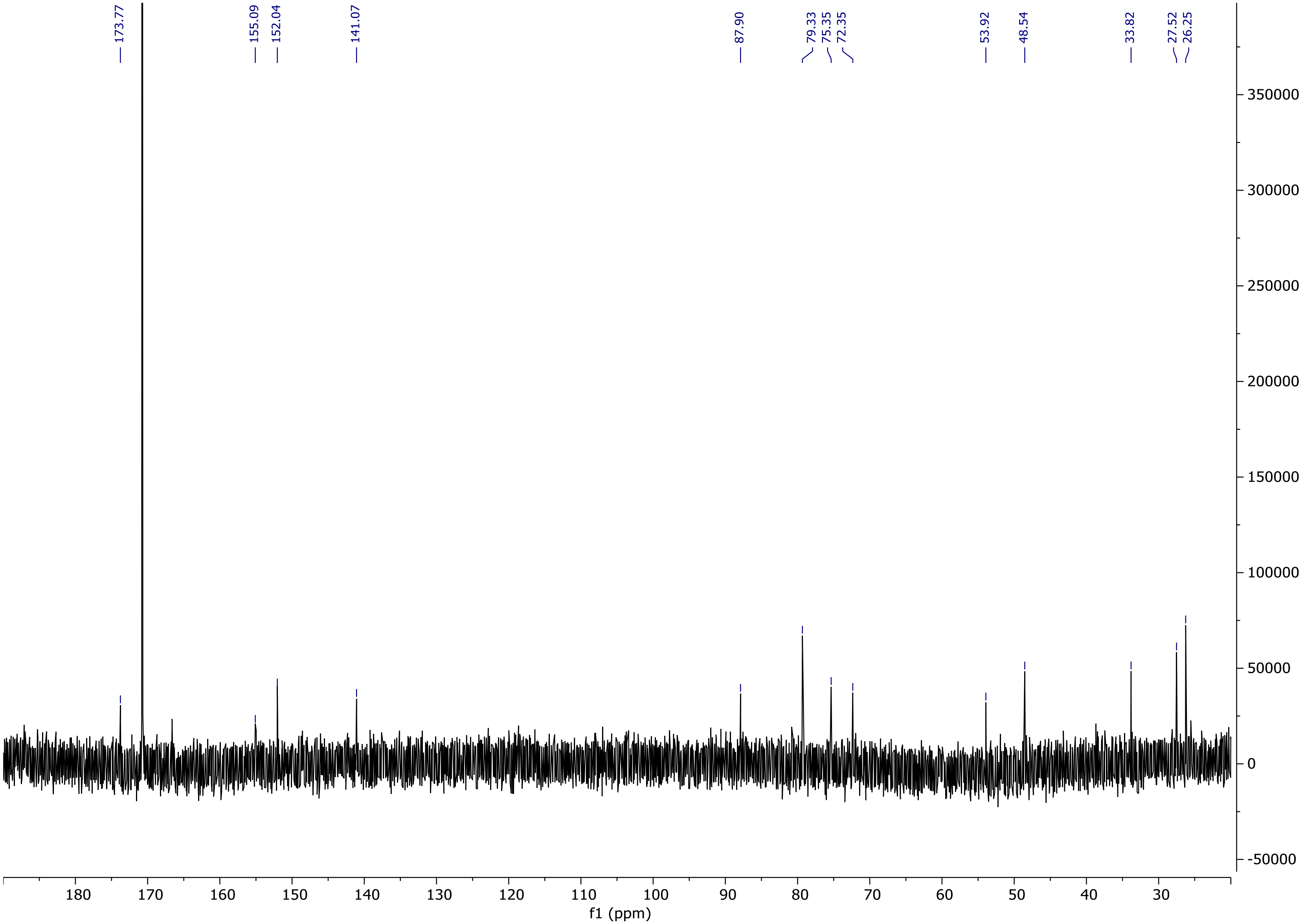

**Figure S45. $^{13}$C NMR spectrum of 2’-phosphosinefungin (9) ($D_2O$, 600 MHz).**

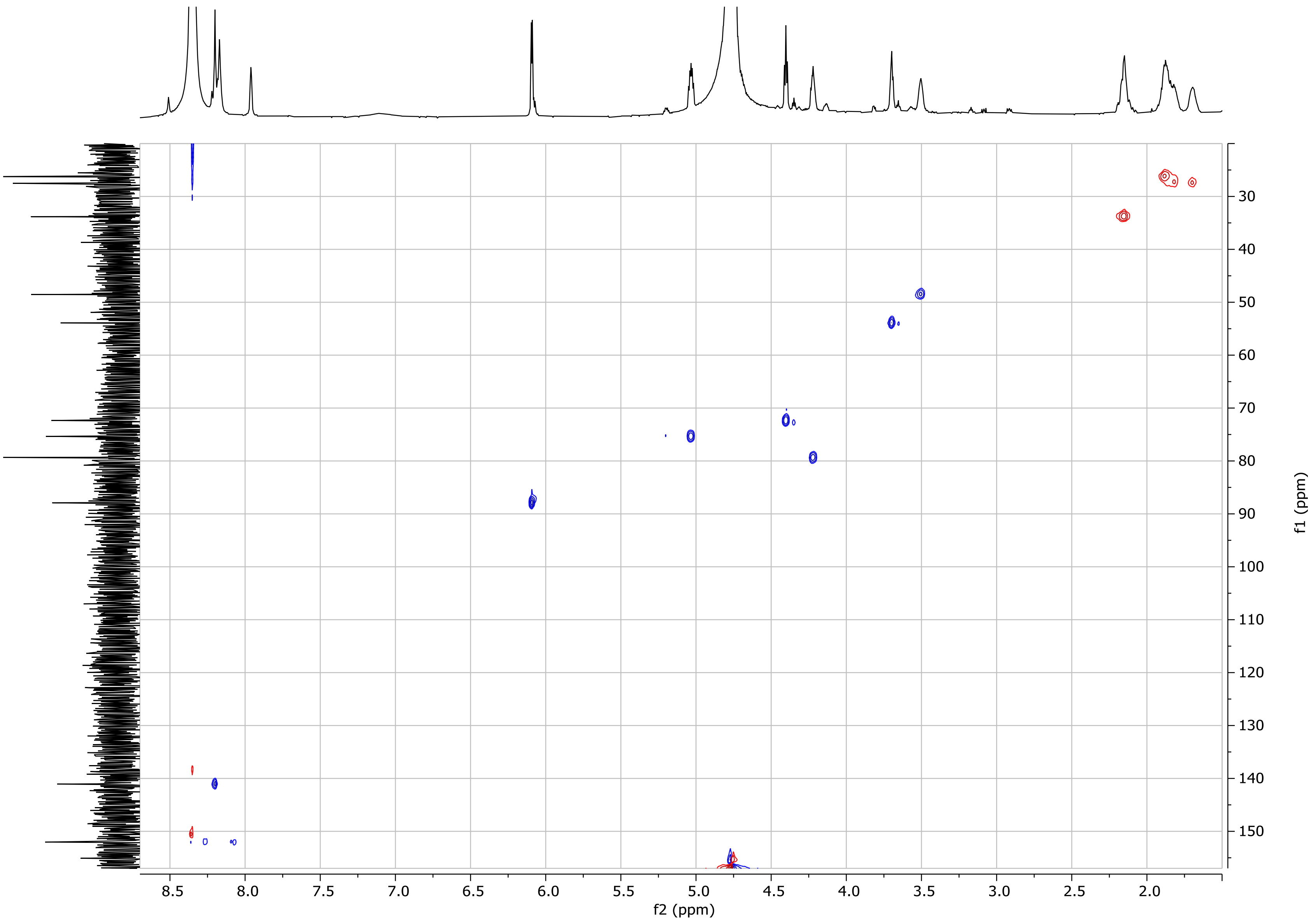

**Figure S46. HSQC spectrum of 2'-phosphosinefungin (9) ($D_2O$).**

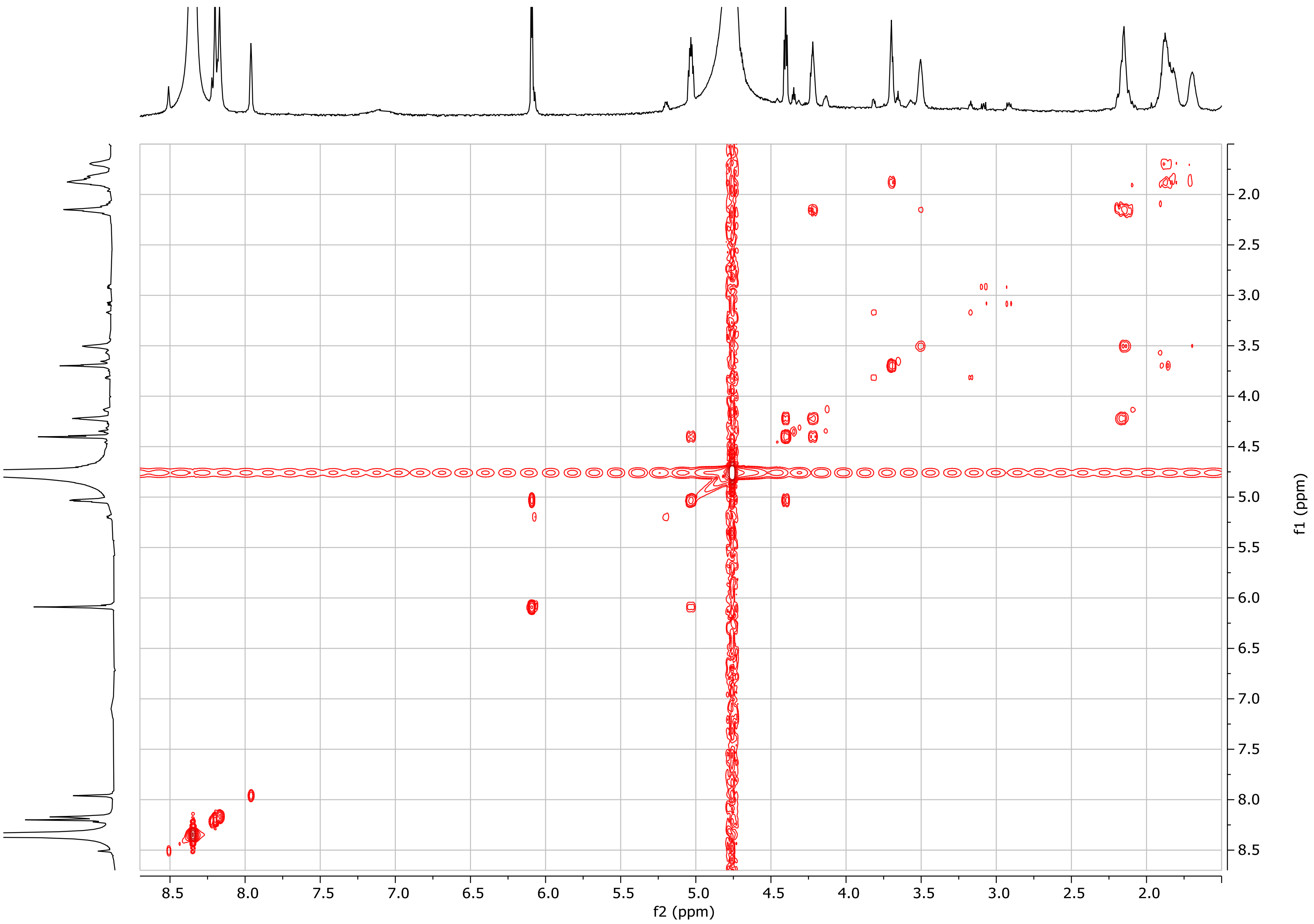

**Figure S47. COSY spectrum of 2'-phosphosinefungin (9) ($D_2O$).**

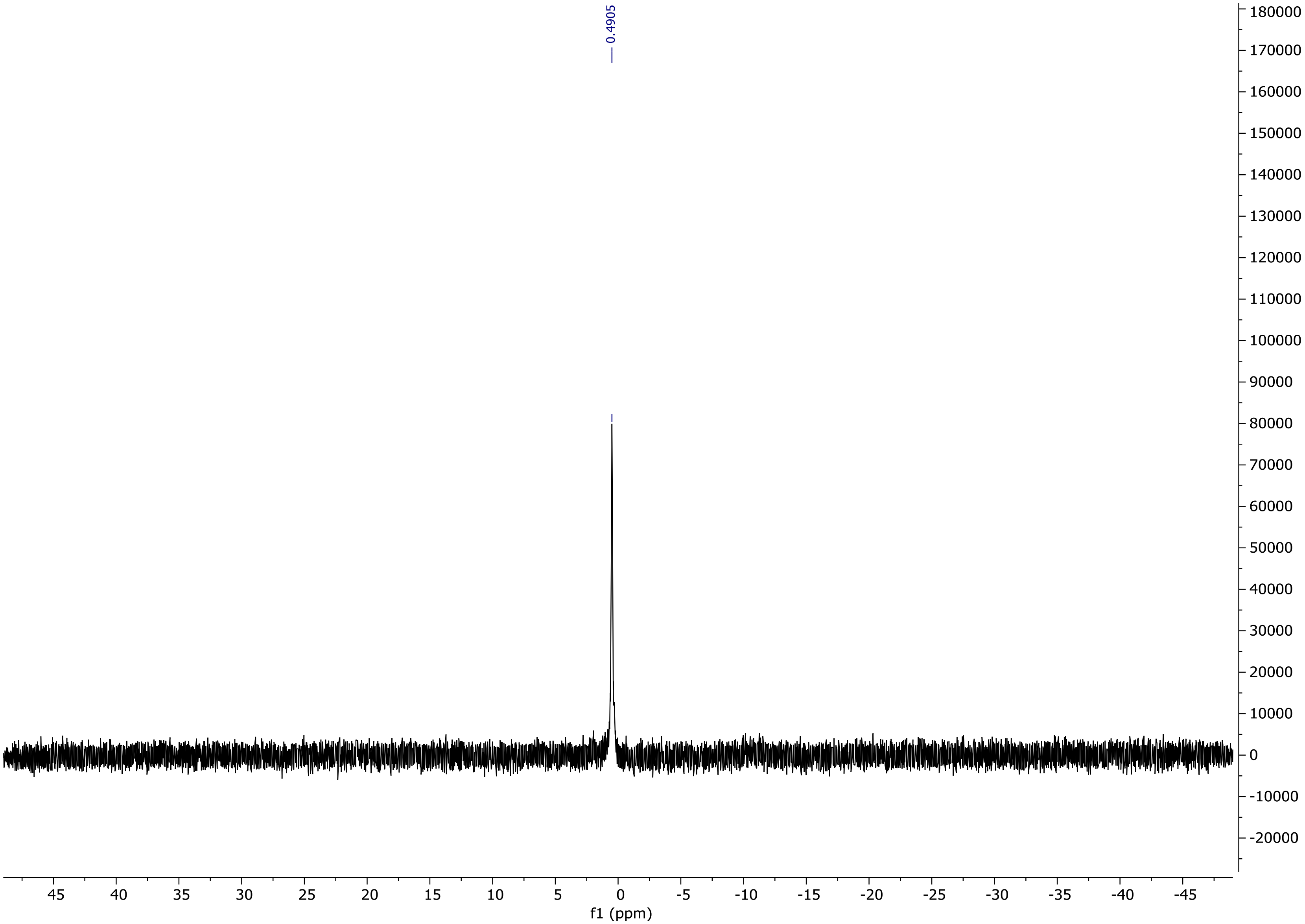

**Figure S48. $^{31}P$ NMR spectrum of 2'-phosphosinefungin (9) ($D_2O$, 242 MHz).**

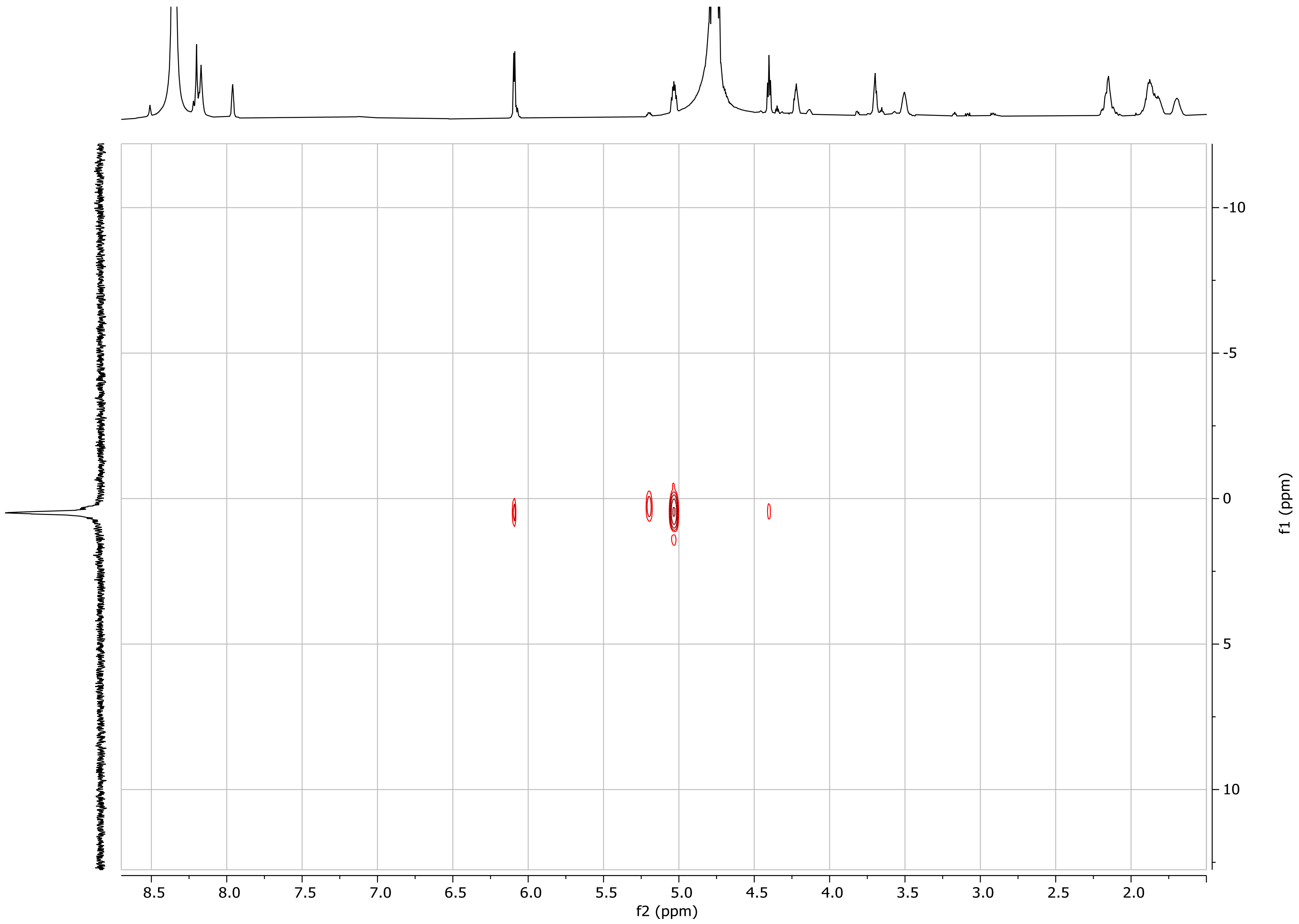

**Figure S49.** $^{1}H$–$^{31}P$ **HMBC spectrum of 2'-phosphosinefungin (9) ($D_2O$).**
